# Functional characterization and lineage analysis of broadly neutralizing human antibodies against dengue virus identified by single B cell transcriptomics

**DOI:** 10.1101/790642

**Authors:** Natasha D. Durham, Aditi Agrawal, Eric Waltari, Derek Croote, Fabio Zanini, Edgar Davidson, Mallorie Fouch, Olivia Smith, Esteban Carabajal, John E. Pak, Benjamin J. Doranz, Makeda Robinson, Ana M. Sanz, Ludwig L. Albornoz, Fernando Rosso, Shirit Einav, Stephen R. Quake, Krista M. McCutcheon, Leslie Goo

**Author notes:** Equal contribution (Natasha D. Durham, Aditi Agrawal). Department of Microbiology and Physiological Systems, University of Massachusetts Medical School, Worcester, United States (Natasha D. Durham). Lowy Cancer Research Center, University of New South Wales, Kensington, NSW, Australia.

## Abstract

Eliciting broadly neutralizing antibodies (bNAbs) against the four dengue virus serotypes (DENV1-4) that are spreading into new territories is an important goal of vaccine design. To delineate bNAb targets, we characterized 28 monoclonal antibodies belonging to expanded and hypermutated clonal families identified by transcriptomic analysis of single plasmablasts from DENV-infected individuals. Among these, we identified two somatically related bNAbs that potently neutralized DENV1-4. Mutagenesis studies revealed that the major recognition determinants of these bNAbs are in E protein domain I, distinct from the only known class of human bNAbs against flaviviruses with a well-defined epitope. B cell repertoire analysis from acute-phase peripheral blood suggested a memory origin and divergent somatic hypermutation pathways for these bNAbs, and a limited number of mutations was sufficient for neutralizing activity. Our study suggests multiple B cell evolutionary pathways leading to DENV bNAbs targeting a novel epitope that can be exploited for vaccine design.

## Introduction

Dengue virus (DENV) is an enveloped, positive-stranded RNA virus belonging to the *Flavivirus* genus, which includes clinically significant human pathogens such as Yellow Fever virus (YFV), Japanese encephalitis virus (JEV), West Nile virus (WNV), and Zika virus (ZIKV). DENV is transmitted to humans via *Aedes* mosquitoes, whose global distribution places half of the world’s population at risk for infection (Kraemer et al., 2019; Messina et al., 2019). Each year, the four phylogenetically and antigenically distinct DENV serotypes (DENV1-4) cause approximately 400 million infections (Bhatt et al., 2013). Additionally, increased global trade, connectivity, and climate change have fueled the expansion of DENV1-4 into new territories (Kraemer et al., 2019; Messina et al., 2014).

Approximately 20% of DENV-infected individuals develop a mild febrile illness, of which 5% to 20% progress to potentially fatal severe disease, characterized by bleeding, plasma leakage, shock, and organ failure (Guzman & Harris, 2015; Khursheed et al., 2013; Thein, Leo, Lee, Sun, & Lye, 2011). Epidemiological studies have shown that pre-existing antibodies from a primary DENV infection are a risk factor for severe disease following subsequent infection with a heterologous DENV serotype (Katzelnick et al., 2017; Salje et al., 2018; Sangkawibha et al., 1984). This is partly attributed to the prevalence of cross-reactive antibodies from the initial infection that can bind, but not neutralize the secondary heterologous virus. Instead, these non-neutralizing antibodies have the potential to facilitate viral uptake into Fc gamma receptor-expressing target cells in a process known as antibody-dependent enhancement (ADE) (Guzman & Harris, 2015; Halstead, 2014). Recent studies of clinical cohorts demonstrated that the risk of severe disease following secondary infection is greatest when pre-existing titers of cross-reactive antibodies fall within a narrow, intermediate range (Katzelnick et al., 2017; Salje et al., 2018). To limit the potential for ADE, an effective vaccine must therefore elicit durable and potent neutralizing antibodies of high titer against DENV1-4 simultaneously. However, the viral and host determinants leading to such bNAbs against flaviviruses are poorly understood.

All of the leading DENV vaccine candidates in clinical development are based on a tetravalent strategy (Scherwitzl, Mongkolsapaja, & Screaton, 2017), which assumes that the use of representative viral strains from each serotype will elicit a balanced and potent polyclonal antibody response to minimize the risk of ADE. However, the suboptimal efficacy and safety profile of a recently licensed DENV vaccine has been partly attributed to its inability to generate a balanced neutralizing antibody response to all four serotypes (Hadinegoro et al., 2015). Additionally, there may be important antigenic differences between circulating and lab-adapted strains (Lim et al., 2019; Raut et al., 2019), as well as among strains even within a given serotype (Bell, Katzelnick, & Bedford, 2019; Katzelnick et al., 2015). Antigenic mismatch between vaccine and circulating strains impacted vaccine efficacy (Juraska et al., 2018), highlighting the importance of rational selection of vaccine components. An alternative strategy, largely exemplified by vaccine development efforts for HIV (Kwong & Mascola, 2018) and respiratory syncytial virus (RSV) (Crank et al., 2019), relies on identifying antibodies with desirable properties and precisely defining their epitopes to guide epitope-based vaccine design (Graham, Gilman, & McLellan, 2019). For antigenically diverse viruses such as DENV, a conserved epitope-based vaccine strategy to elicit a broad and potent monoclonal neutralizing antibody response could mitigate the challenge of selecting representative vaccine strains.

The main target of flavivirus neutralizing antibodies is the envelope (E) glycoprotein, which consists of three structural domains (DI, DII, DIII), and is anchored to the viral membrane via a helical stem and transmembrane domain. The E proteins direct many steps of the flavivirus life cycle, including entry, fusion, and assembly of new virus particles (Pierson & Diamond, 2012). Flaviviruses bud into the endoplasmic reticulum lumen as immature particles with a spiky surface on which E proteins associate into sixty heterotrimers with a chaperone protein, prM (Prasad et al., 2017; Y. Zhang et al., 2003; Y. Zhang, Kaufmann, Chipman, Kuhn, & Rossmann, 2007). Within the low pH environment of the trans-golgi network, E proteins undergo conformational changes that allow furin-mediated cleavage of prM (Yu et al., 2008), resulting in the release of mature infectious virions with a smooth surface densely coated with ninety E homodimers (Kostyuchenko et al., 2016; Kuhn et al., 2002; Mukhopadhyay, Kim, Chipman, Rossmann, & Kuhn, 2003; Sirohi et al., 2016; X. Zhang et al., 2013). The dense arrangement of E proteins on the virion surface is important for antigenicity, as many potently neutralizing human antibodies against flaviviruses target quaternary epitopes spanning multiple E proteins (de Alwis et al., 2012; Hasan et al., 2017; Kaufmann et al., 2010; Rouvinski et al., 2015; Teoh et al., 2012).

Recent advances in monoclonal antibody isolation and characterization (Boonyaratanakornkit & Taylor, 2019; Corti & Lanzavecchia, 2014) have accelerated the identification of bNAbs, including those against flaviviruses. Examples include antibodies d488 (Li et al., 2019) and m366 (Hu et al., 2019), which were cloned from B cells of rhesus macaques receiving an experimental DENV vaccine and from healthy flavivirus-naive humans, respectively, and mAb DM25-3, which was isolated from a mouse immunized with a mature form of DENV2 virus-like particles ((Shen et al., 2018). Although these antibodies are cross-reactive against DENV1-4, they demonstrated only moderate potency. E protein residues involved in d488 binding lie at the interface of the M protein and the E protein ectodomain (Li et al., 2019), while those for m366.6 binding appear to be located at the dimerization interface between DII and DIII (Hu et al., 2019). Residue W101 within the DII fusion loop was identified to be important for recognition by mAb DM25-3 (Shen et al., 2018). Attempts to engineer mouse antibodies with increased breadth and potency against DENV1-4 have also been described (Deng et al., 2011; Shi et al., 2016; Tharakaraman et al., 2013).

Only a few naturally occurring human bNAbs against flaviviruses have been characterized. Many of these antibodies target epitopes consisting of the DII fusion loop as well as the adjacent bc loop in DII in some cases (Smith et al., 2013; Tsai et al., 2013; Xu et al., 2017). Although antibodies recognizing the highly conserved fusion loop can demonstrate broad reactivity to all DENV serotypes and related flaviviruses, their neutralizing potency is often limited due to this epitope being largely inaccessible, especially on mature virions (Cherrier et al., 2009; Nelson et al., 2008; Shen et al., 2018; Stiasny, Kiermayr, Holzmann, & Heinz, 2006). To date, the most well-characterized class of human antibodies with broad and potent neutralizing activity against flaviviruses targets a conserved, quaternary epitope spanning both E monomeric subunits within the dimer. A subset of these E dimer epitope (EDE)-specific bNAbs potently neutralize not only DENV1-4, but also ZIKV, owing to the high conservation of the EDE, which overlaps the prM binding site on E (Barba-Spaeth et al., 2016; Dejnirattisai et al., 2015; Rouvinski et al., 2015). The exciting discovery of the EDE class of bNAbs highlights the potential for an epitope-focused flavivirus vaccine strategy.

As multiple specificities are likely required to provide maximum coverage of diverse circulating viral variants (Bell et al., 2019; Doria-Rose et al., 2012; Goo, Jalalian-Lechak, Richardson, & Overbaugh, 2012; Katzelnick et al., 2015; Keeffe et al., 2018; Kong et al., 2015), in this study, we aimed to define novel sites on the flavivirus E protein that can be targeted by bNAbs. By characterizing 28 monoclonal antibodies from the plasmablasts of two DENV-infected individuals, we identified J8 and J9, clonally related bNAbs that neutralized DENV1-4 in the low picomolar range. The major recognition determinants for J8 and J9 were in E protein DI, distinct from previously characterized bNAbs. Analysis of the corresponding B cell repertoire revealed divergent evolution of J8 and J9, suggesting multiple evolutionary pathways to generate bNAbs within this lineage. Our work identifies both viral and host determinants of the development of DENV bNAbs that can guide immunogen design and evaluation.

## Results

### Identification of cross-reactive neutralizing antibodies from clonally expanded plasmablasts of DENV-infected individuals

We previously profiled the single-cell transcriptomics of peripheral blood mononuclear cells (PBMCs) from six dengue patients and four healthy individuals (Zanini et al., 2018). In two DENV-infected patients (013 and 020), we identified 15 clonal families comprising a total of 38 unique paired heavy (VH) and light (VL) chain IgG1 plasmablast sequences, some of which were hypermutated (1.67% to 10.77% for VH, 0.67% to 7.22% for VL; Figure S1). One clonal family (CF) included members found in both individuals (antibodies B10, M1, and D8 from CF1, Figure S1), suggesting convergent evolution, which has been described for the antibody response to distinct viruses, including flaviviruses (Parameswaran et al., 2013; Robbiani et al., 2017), Ebola virus (Davis et al., 2019) and HIV (Scheid et al., 2011; Wu et al., 2011). To functionally characterize these monoclonal antibodies (mAbs), we successfully cloned 36 paired VH and VL sequences into expression vectors, and transfected mammalian cells for small scale (96-well) recombinant IgG1 production. We detected secreted IgG in the transfection supernatants for 28 of 36 mAbs, which were tested for binding to DENV2 recombinant soluble E protein (rE) and reporter virus particles (RVPs), as well as for neutralizing activity against a panel of flavivirus RVPs, including DENV1-4, ZIKV, and WNV (Figure S1). Seventeen of 28 mAbs bound to either DENV2 rE (n = 1), or RVPs (n = 7), or both (n = 9). None of the mAbs neutralized ZIKV, but all 28 neutralized at least one DENV serotype, 21 mAbs neutralized two or more DENV serotypes, and one mAb neutralized WNV in addition to DENV.

### Binding profile of mAbs

For further characterization, we selected six mAbs for larger scale production and IgG purification based on their ability to neutralize at least four of the six RVPs tested. These included two mAb clonal variants found in both patients (B10 from patient 020 and M1 from patient 013), two (C4, J9) and one (L8) patient 013 and patient 020 mAbs, respectively, that neutralized DENV1-4, and the only mAb that neutralized WNV (I7 from patient 020). We first confirmed binding activity at a single antibody concentration (5 μg/ml) by ELISA. Consistent with our pilot screen using crude IgG-containing supernatant (Figure S1), I7, M1, B10, and L8 bound to both rE and RVPs, while C4 and J9 bound to RVPs only (Figure 1A & B), suggesting that these mAbs target epitopes preferentially displayed on the intact virion. As incubation at higher temperatures has been shown to improve exposure of some epitopes (Dowd, Jost, Durbin, Whitehead, & Pierson, 2011; Lok et al., 2008; Sukupolvi-Petty et al., 2013), we performed the ELISA at both ambient temperature and 37°C. For most antibodies, incubation at 37°C resulted in a modest but consistent increase in RVP binding (Figure 1B). To evaluate the relative binding of the mAbs to rE and RVP, we also performed a dose-responsive indirect ELISA at ambient temperature (Figure 1C). Binding curves revealed robust binding of mAbs L8, B10, M1, and I7 to DENV2 rE (EC50 range of 0.4 to 23 ng/ml) while J9, C4 and the EDE mAbs C10 and B7 displayed little to no binding to rE even at high antibody concentrations (up to 200 μg/mL). All mAbs showed binding to DENV2 RVP to varying extents, with relatively high EC50 values for J9 (200 ng/ml), J8 (213 ng/ml), and C4 (1200 ng/ml), suggesting limited affinity maturation (Figure 1D).

**Figure 1.**
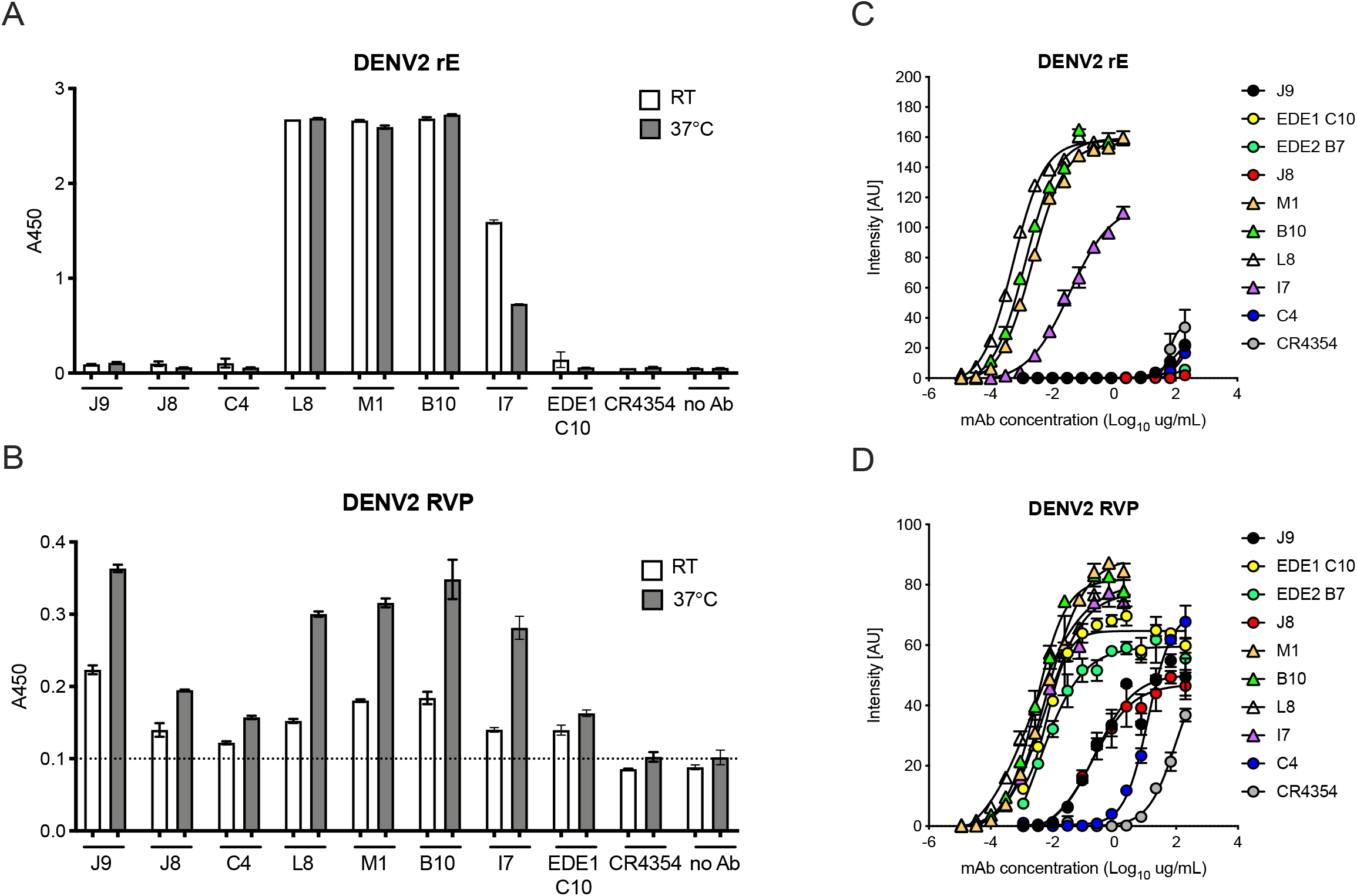
Binding profile of mAbs. A single dilution of the mAbs indicated on the x-axis was tested for binding to DENV2 **(A)** soluble recombinant E protein (rE) and **(B)** reporter virus particles (RVPs) at room temperature (RT) and 37°C by ELISA. The y-axis shows absorbance values at 450 nm. Error bars indicate the range of values obtained in duplicate wells. Data are representative of 3 independent experiments. The dotted horizontal line in (B) indicates the average A450 values obtained for negative control WNV-specific mAb CR4354. Representative dose-response binding curves of the indicated mAbs to DENV2 **(C)** rE and **(D)** RVPs at room temperature. The y-axis shows binding signal intensity in arbitrary units (AU). Error bars indicate the standard deviation (SD) of triplicate spots within one well of the microarray. Binding curves are representative of two independent experiments.

### Neutralization potency of mAbs

We next performed dose-response neutralization assays to obtain IC_50_ values (mAb concentration at which 50% of virus infectivity was inhibited). Antibodies M1, B10, and L8 displayed modest (average IC_50_ range of 379 to 796 ng/ml) and incomplete neutralization of DENV1-4, with ∼10% to ∼50% infectivity persisting at the highest mAb concentration tested (10 μg/ml) (Figures 2A-B). Incomplete neutralization is commonly observed for cross-reactive DII fusion loop-specific antibodies, and likely represents structurally heterogeneous virions on which the epitope is not displayed frequently enough for antibodies to bind at a stoichiometry sufficient for neutralization (Nelson et al., 2008; Pierson et al., 2007). Antibody I7 displayed an unusual neutralization profile as it did not neutralize DENV4, and its potency against DENV1-3 was lower than that against the more antigenically distant WNV. Although mAb C4 completely neutralized DENV1-4, it did so with modest potency, especially against DENV4 (IC_50_ > 1000 ng/ml). The most potent mAb we identified was J9, which despite relatively weak binding (Figure 1), completely neutralized DENV1-4 with average IC_50_ values of 6 ng/ml, 30 ng/ml, 15 ng/ml, and 39 ng/ml, respectively. A previously characterized subgroup of the EDE class of bNAbs, which includes mAb EDE1 C10 can neutralize not only DENV1-4, but also ZIKV (Barba-Spaeth et al., 2016). J9 showed a high specificity for DENV, with no activity against ZIKV, and up to ∼60-fold greater potency against some DENV serotypes compared to EDE1 C10 (Figure 2B). Depending on the serotype, the average neutralization potency of J9 against DENV1-4 was also up to 15-fold higher than that of bNAb EDE2 B7, which belongs to another EDE subgroup with poor neutralizing activity against ZIKV (Barba-Spaeth et al., 2016).

**Figure 2.**
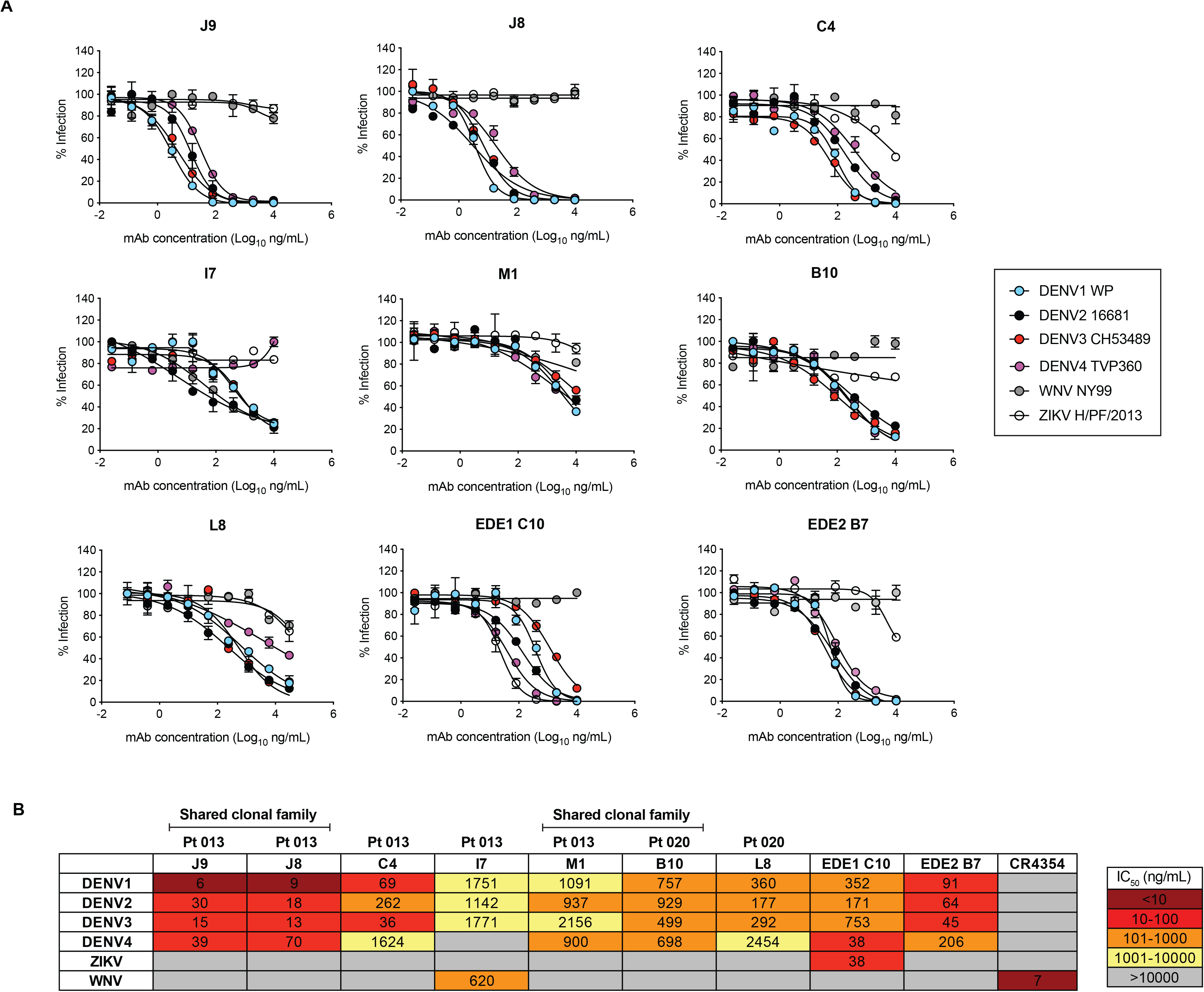
Neutralization profile of mAbs. **(A)** Representative mAb dose-response neutralization curves against DENV1-4, ZIKV, and WNV RVPs. Infectivity levels were normalized to those observed in the absence of antibody. Data points and error bars indicate the mean and range, respectively. Results are representative of at least 3 independent experiments, each performed in duplicate. **(B)** MAb concentrations resulting in 50% inhibition of infectivity (IC_50_) from dose-response neutralization experiments described in (A). Values represent the mean of at least 3 independent experiments, each performed in duplicate. The heatmap indicates neutralization potency, as defined in the key. Grey boxes indicate that 50% neutralization was not achieved at the highest mAb concentration tested (10 μg/ml). The patient (Pt) from which mAbs were isolated are indicated above each mAb name. MAbs from shared clonal families are indicated.

The broad and potent neutralizing activity of J9 prompted us to re-evaluate our pilot results obtained for J8, a somatic variant with no binding or neutralizing activity in our screen with crude IgG-containing supernatant (Figure S1). When we repeated the cloning, expression, and purification of J8 IgG, we observed similar binding (Figure 1) and neutralization (Figure 2) profiles to J9. When tested as Fab fragments, J9 and J8 were still able to potently neutralize DENV, unlike C4 and EDE1 C10, which failed to achieve 50% neutralization at the highest mAb concentration tested (Figure S2). J9 and J8 also potently neutralized contemporary DENV1-4 isolates with IC_50_ values < 50 ng/ml (Figure S3). Additionally, in contrast to C4 and the cross-reactive DII fusion loop-specific mouse mAb E60 (Goo, VanBlargan, Dowd, Diamond, & Pierson, 2017; Nelson et al., 2008; Oliphant et al., 2006), but similar to EDE bNAbs (Dejnirattisai et al., 2015), J9 and J8 potently neutralized DENV regardless of virion maturation state (Figure S4), which can indirectly modulate epitope exposure (Cherrier et al., 2009; Goo et al., 2019; Nelson et al., 2008) and has been shown to be distinct among circulating versus lab-adapted strains (Raut et al., 2019). We also tested the ability of mAbs to mediate neutralization after virus attachment to cells, which is characteristic of many potently neutralizing antibodies against flaviviruses (Goo et al., 2019; Nybakken et al., 2005; Sukupolvi-Petty et al., 2010; Vogt et al., 2009; Xu et al., 2017). When added after virus attachment to Raji-DCSIGNR cells, C4 failed to inhibit 40-50% of infection at the highest mAb concentration tested (300 μg/ml) (Figure 3). In contrast, J9, J8, and EDE1 C10 potently inhibited DENV2 infection both pre- and post-virus attachment to cells.

**Figure 3.**
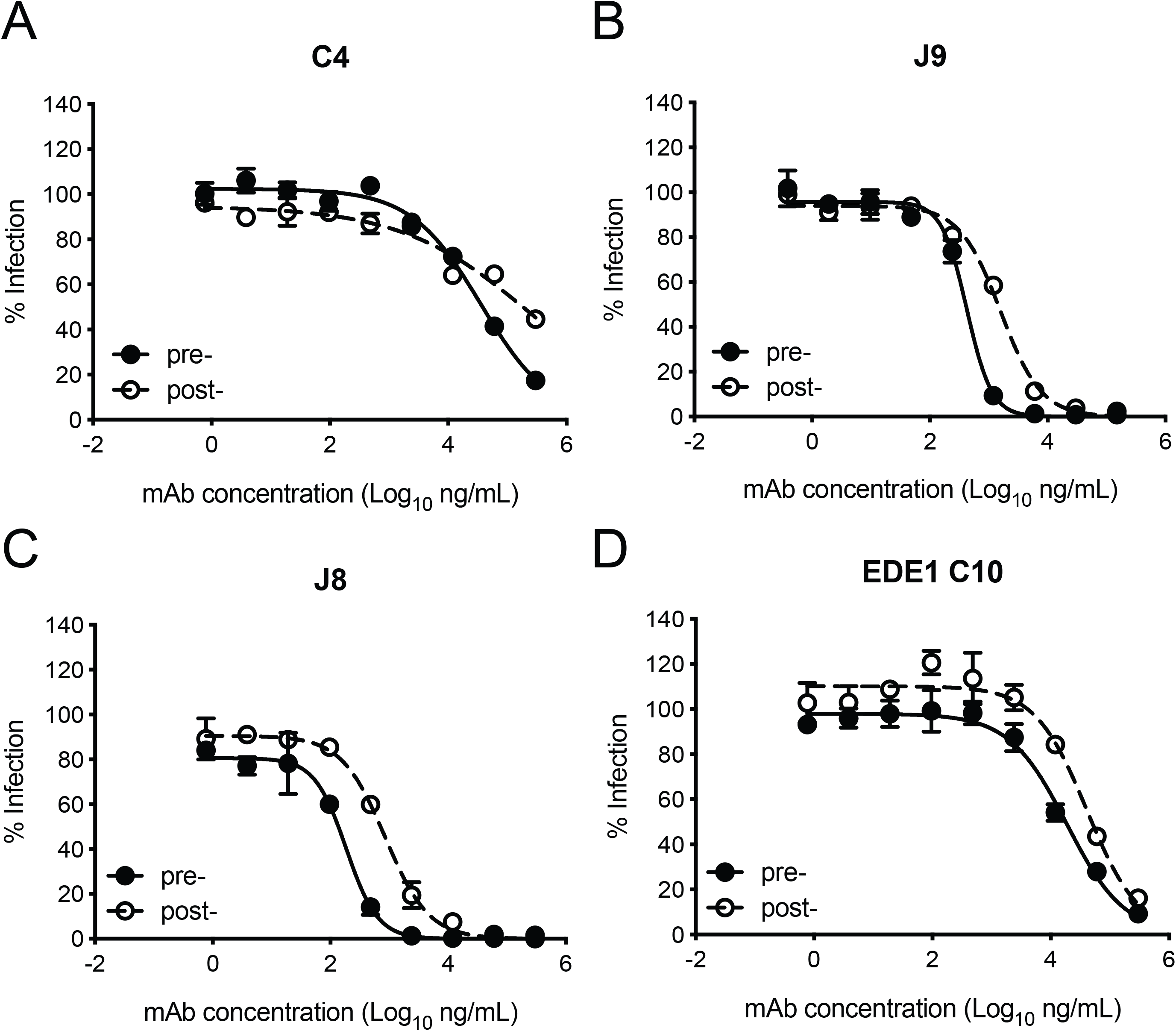
Mechanism of neutralization. Representative dose-response curves for pre- and post-attachment neutralization of DENV2 16681 RVPs by the indicated mAbs. Results are representative of at least 2 independent experiments each performed in duplicate. Data points and error bars indicate the mean and range, respectively.

### ADE potential of mAbs

*In vitro*, antibodies can mediate ADE of infection in cells expressing Fc gamma receptor (FcγR) at sub-neutralizing concentrations (Pierson et al., 2007). Recent studies in humans have also demonstrated that the risk of severe dengue disease following secondary infection is greatest within a range of intermediate titers of pre-existing DENV-specific antibodies, while higher titers are protective against symptomatic infection (Katzelnick et al., 2017; Salje et al., 2018). Thus, eliciting potently neutralizing antibodies is desirable to limit the concentration range within which ADE can occur. We measured the ADE potential of a subset of the mAbs identified above in K562 cells, which have been used extensively to study ADE of flaviviruses as they express FcγR and are poorly permissive for infection in the absence of antibody (Littaua, Kurane, & Ennis, 1990). As expected, all DENV-specific mAbs mediated ADE of DENV2 infection to varying extents (Figure S5). We measured the antibody concentration at which the highest level of ADE was observed (peak enhancement titer). Consistent with their high neutralization potencies, the average peak enhancement titer of J9 and J8 for DENV2 (3 ng/ml) was approximately 27-fold and 480-fold lower than that of mAbs EDE1 C10 (80 ng/ml) and C4 (1467 ng/ml), respectively (Figures S5A and S5D). For J9, J8, and EDE bNAbs, DENV2 neutralization occurred beyond the peak enhancement titer, with no infectivity observed at high antibody concentrations. In contrast, for C4 and L8, which neutralized DENV relatively weakly (Figure 1), ADE of DENV2 was still detected at the highest concentration (10 μg/ml) tested (Figure S5A). Even at high concentrations, L8 also mediated ADE of ZIKV (Figure S5B) and WNV (Figure S5C), suggesting binding, but not neutralizing activity against these viruses (Figure 1). Consistent with their ability to recognize ZIKV, EDE bNAbs enhanced ZIKV infection at sub-neutralizing concentrations (Figure S5B). J9 and J8 did not facilitate ADE of ZIKV nor WNV infection, suggesting lack of binding to these flaviviruses. Given their high neutralization potencies against DENV1-4, J9 and J8 represent desirable antibodies to elicit as their ADE potential is restricted to a narrow range of low antibody concentrations, beyond which neutralization is observed.

### Epitope specificity of mAbs

To identify amino acid residues required for mAb recognition, we screened a shotgun alanine-scanning mutagenesis library of DENV2 E protein variants for mAb binding by flow cytometry (Davidson & Doranz, 2014). MAb binding profiles to this entire library are summarized in Table S1. MAbs M1 and L8 demonstrated loss of binding to variants encoding W101A and F108A mutations within the DII fusion loop (Figure 4A). Similar to a subset of previously described fusion loop-specific antibodies (Cherrier et al., 2009; Smith et al., 2013; Tsai et al., 2013), M1 recognition also depended on residue G106 within the fusion loop and residue 75 on the adjacent bc loop (Figure 4A). Specificity for the fusion loop epitope likely contributes to the inability of these mAbs to neutralize completely, even at high concentrations (Figures 2A-B), as previously described (Dowd et al., 2011; Nelson et al., 2008). The I7 epitope involved DII residues Q256 and G266 (Figure 4B), which are conserved among many flaviviruses (indicated by yellow squares in Figure S6A alignment) and are important for recognition by the recently described cross-reactive mAb (d448) isolated from vaccinated rhesus macaques (Li et al., 2019). Unlike I7, d448 neutralized DENV4, but not WNV (Li et al., 2019). Despite testing different temperature and pH conditions, we did not detect C4 binding to WT DENV2 in this flow cytometry-based assay (data not shown) and were thus unable to screen against the mutant library. For J9, most E protein mutations that reduced binding by >80% relative to WT (shown in bold in Table S1) were found in DI (R2, I4, K47, S145, H149, N153, T155), but G102 within DII fusion loop and DII N242, as well as DIII N366 were also important (Figure 4C). These mutations minimally affected EDE1 C10 binding (Figure 4C).

**Figure 4.**
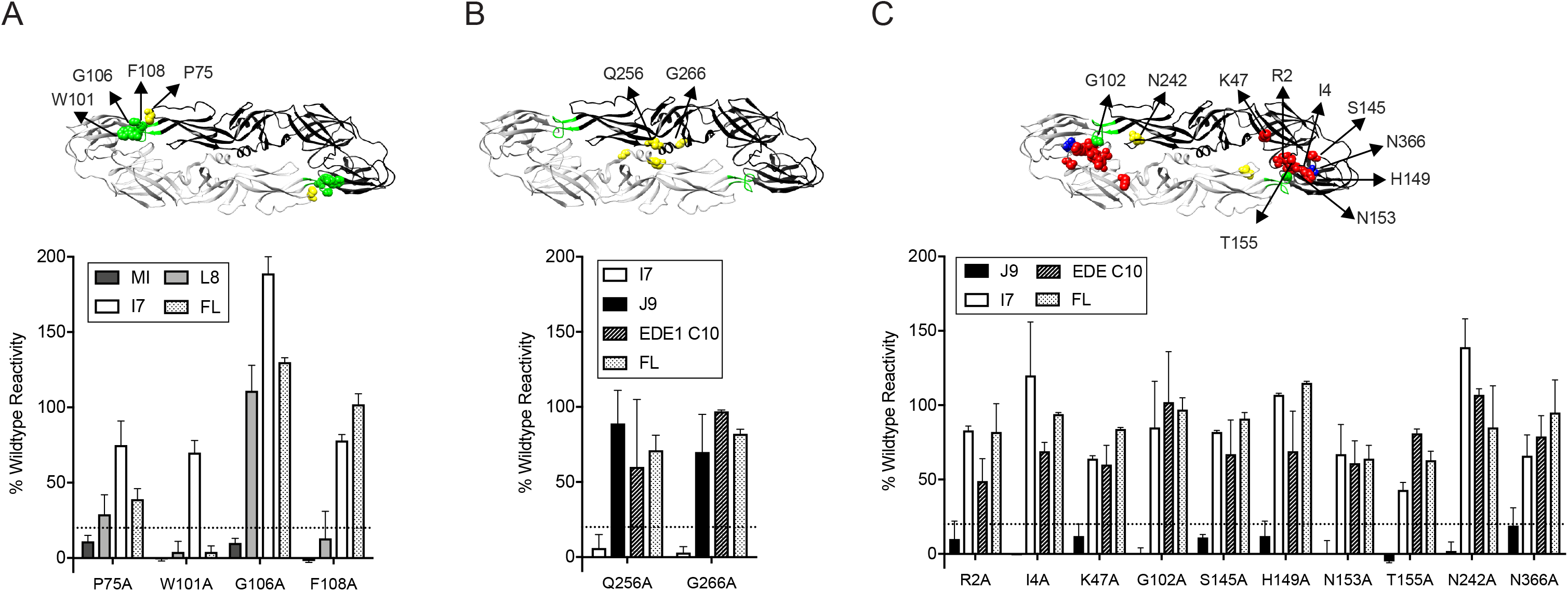
Critical E protein residues for mAb binding. Individual alanine mutation of a subset of DENV2 E residues decreased binding by mAbs **(A)** M1 and L8, **(B)** I7 or **(C)** J9 as shown, but did not affect binding by other mAbs, including EDE1 C10 and a previously screened control mAb (FL) targeting the fusion loop (unpublished). Error bars represent the mean and range (half of the maximum-minus-minimum values) of at least two biological replicates. The dotted horizontal line indicates 80% reduction in mAb binding reactivity to mutant compared to wildtype DENV2 E. Above each graph, residues involved in binding of the corresponding mAb are highlighted on the ribbon structure of one of the monomers (black) within the DENV2 E dimer (PDB: 1OAN). Residues in DI, DII, DIII, and DII fusion loop are indicated in red, yellow, blue, and green, respectively.

For further epitope mapping of J9, one of the most potent bNAbs we identified, we first attempted to select for neutralization escape viral variants but were unsuccessful after six serial passages in cell culture under mAb selection pressure. We noted that J9 neutralized DENV1-4 but not ZIKV (Figure 1). As an alternative epitope mapping approach, we generated a panel of DENV2 RVP variants encoding individual mutations at solvent accessible E protein residues on the mature DENV virion that are identical or chemically conserved across representative DENV1-4 strains but differ from ZIKV (Figure S6A). These residues in DENV2 16681 E were substituted for analogous residues in ZIKV H/PF/2013. We also included a subset of alanine mutants identified by our binding screen to confirm their importance for neutralization. In total, we generated 34 DENV2 RVP variants encoding individual mutations throughout the E protein (Figure S6B); 31 of these variants retained sufficient infectivity (Figure S7) for neutralization studies. Most individual mutations displayed minimal (< 2-fold) effects on J9 neutralization potency (Figure S8A). Two DI mutations, K47T and V151T resulted in a modest 4-fold increase in IC_50_ (Figure S8A). Consistent with our binding screen, we confirmed that mutation at residue N153 or T155, each of which results in a loss of a potential N-linked glycosylation site, abrogated J9 neutralization, while mutation at N242 resulted in a 5-fold reduction in potency (Figures S8A). Although our binding screen suggested that individual G102A and S145A mutations contributed to J9 recognition, they had limited effects (∼2-fold) on neutralization potency (Figure S8A).

Because we observed only modest effects with single mutations, we next generated DENV2 RVPs encoding the K47T and V151T mutations in combination, as well as 8 additional pairs of mutations at a subset of the above 34 residues selected based on their proximity to each other on the E dimer structure (Figure S6C). Seven of 8 DENV2 RVP variants encoding paired mutations retained infectivity (Figure S7). Five combinations of paired mutations displayed similarly modest (up to 4-fold) effects on J9 neutralization potency as when these mutations were tested individually (Figure S8A). However, in combination, K47T+V151T reduced J9 potency by almost 100-fold. In combination with the F279S mutation in DI, K47T also resulted in a 16-fold average reduction in J9 potency (Figure S8A). Figures 5A-B highlight key residues that reduced J9 neutralization potency, either alone or in combination, as identified from our screen against the entire panel of single and double mutants (Figure S8).

**Figure 5.**
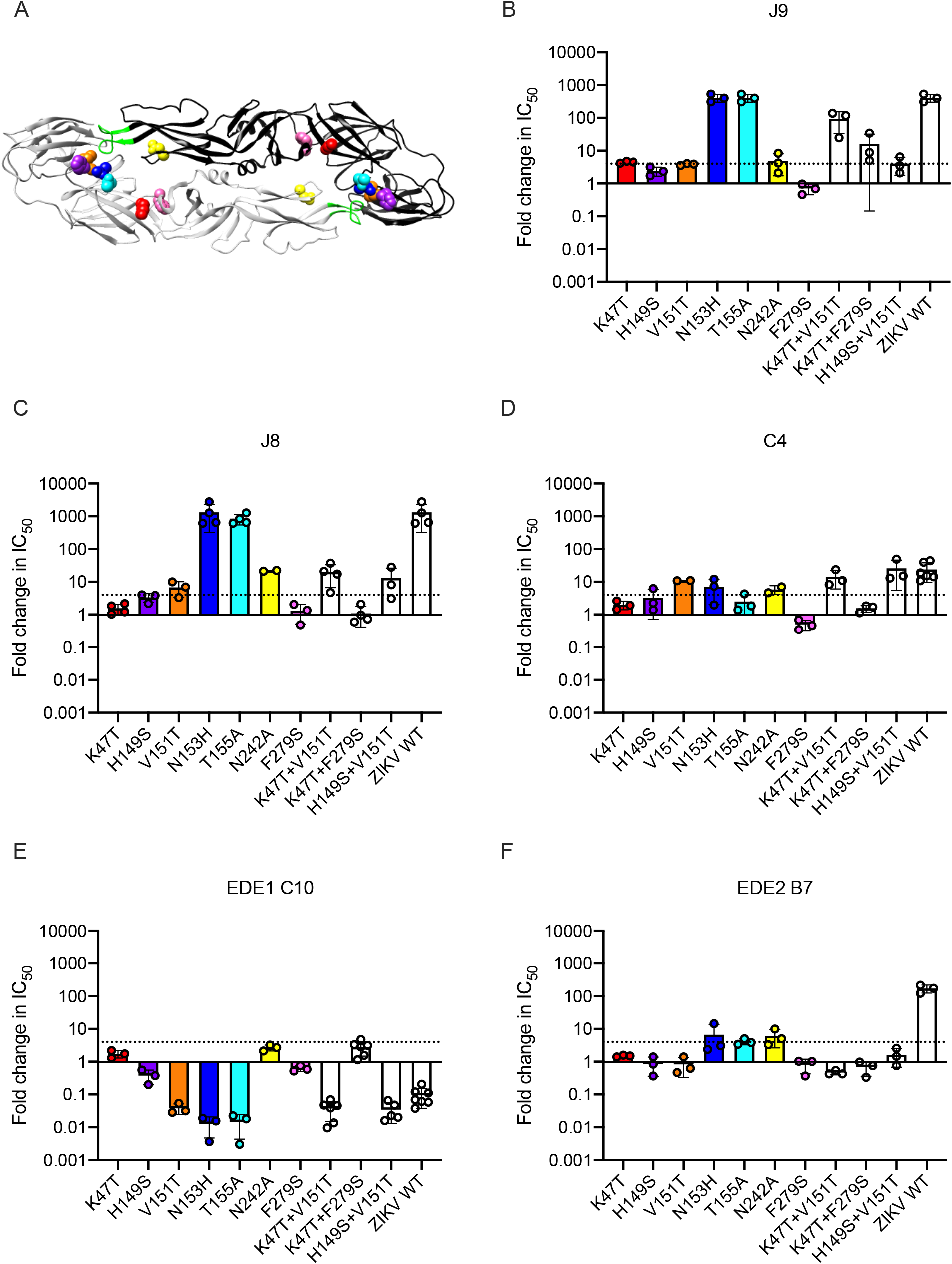
Critical E protein residues for mAb neutralization. **(A)** Ribbon structure of the DENV2 E dimer (PDB: 1OAN) with one monomer in black and the other in grey. The conserved DII fusion loop is shown in green. Colored spheres indicate residues that contribute to J9 recognition identified in Figure S8A and summarized in (B). Bar graphs depict the mean fold change in IC_50_ values against the DENV2 E variants indicated on the x-axis relative to DENV2 wildtype RVPs for mAbs **(B)** J9, **(C)** J8, **(D)** C4, **(E)** EDE1 C10, and **(F)** EDE2 B7. For each mAb, wildtype ZIKV RVPs were included as controls. Mean values were obtained from 2 to 7 independent experiments represented by data points. Error bars indicate the standard deviation (n > 2 experiments) or range (n = 2 experiments). Bar colors correspond to those of spheres in (A) to indicate location within the E dimer. The dotted horizontal line indicates a 4-fold increase in IC_50_ value relative to wildtype DENV2.

As seen for J9, individual mutations at residues N153 and N155, which together encode a potential N-linked glycosylation site, abrogated the neutralizing activity of the somatically related bNAb, J8 (Figure 5C). To varying extents, a similar set of individual and paired mutations that reduced J9 neutralization potency, including V151T, N242A, K47T+V151T, and H149S+V151T, also reduced J8 and C4 neutralization potency (Figures 5C-D). However, individual mutations at some DII and DIII residues that did not impact J9 neutralization potency (Figure S8A) did modestly increase resistance to neutralization by J8 (Figure S8B) and C4 (Figure S8C), respectively by 4-to 7-fold. Interestingly, the V151T mutation alone or in combination with either K47T or H149S increased neutralization potency of EDE1 C10 by 25-fold (Figure 5E). These mutations had minimal (< 2-fold) effects on the neutralizing activity of EDE2 B7 (Figure 5F) as well as patient 013 polyclonal serum (Figure S9), suggesting that they did not globally alter antigenicity. As previously shown (Dejnirattisai et al., 2015; Rouvinski et al., 2015), mutation at residue N153 or N155 improved and disrupted EDE1 C10 and EDE2 B7 recognition, respectively (Figures 5E-F). Overall, these results suggest that the recognition determinants of J9, J8, and C4, are distinct from those targeted by EDE-specific and other previously characterized bNAbs against DENV1-4 (Dejnirattisai et al., 2015; Hu et al., 2019; Li et al., 2019; Smith et al., 2013; Tsai et al., 2013; Xu et al., 2017).

### Lineage analysis reveals memory origin and divergent evolution of bNAbs

To gain insight into the development of bNAbs J9 and J8, we processed PBMCs of patient 013 (from which these bNAbs were identified) obtained four days post-fever onset, and performed next generation sequencing (NGS) of the B cell receptor (BCR) repertoire. This patient experienced acute secondary infection with DENV4 (Zanini et al., 2018). Given its greater junctional diversity compared to light chain, we focused our analysis on the heavy chain repertoire, which is sufficient to identify clonal relationships (Zhou & Kleinstein, 2019). We have recently shown that PBMC stimulation in a polyclonal, BCR-independent manner can selectively expand antigen-specific memory B cells (Waltari, McGeever, Friedland, Kim, & McCutcheon, 2019). Accordingly, we obtained 8-fold more unique VH sequences from stimulated PBMCs with a greater representation of IgG over IgM clonal families compared to unstimulated PBMCs (Table S2). Compared to previously described healthy BCR repertoire data (Waltari et al., 2019), VH1-69, VH3-30, VH3-30-3, VH4-34, VH4-39 and VH4-59 were the most dominant across both unstimulated and stimulated PBMC VH families in patient 013 (average >= 5% of repertoire; Figure S10).

We found 579 and 43,179 VH sequences related to J9/J8 in unstimulated and stimulated PBMCs, respectively (0.4% and 3.5% of total reads, respectively, Table S3). For lineage construction (Figure 6A), we included sequences that met one of three criteria: 1) highest numbers of unique molecular identifier (UMI) counts (>35 in PBMC repertoire and >150 in stimulated PBMC repertoire IgG sequences and >15 in IgA sequences), 2) < 5% somatic hypermutation, or 3) 97% identity to J9 or J8. The J9/J8 lineage derived from recombination of IGHV1-69 with IGHD2-2 and IGHJ5 with no CDRH3 insertions or deletions (Figure 6B). The majority of the clonal family members were of the IgG_1_ subtype, with no IgM identified having a UMI count >2 and only a small percentage of IgA (1.8% of stimulated PBMC relatives; Figure 6A, triangles).

**Figure 6.**
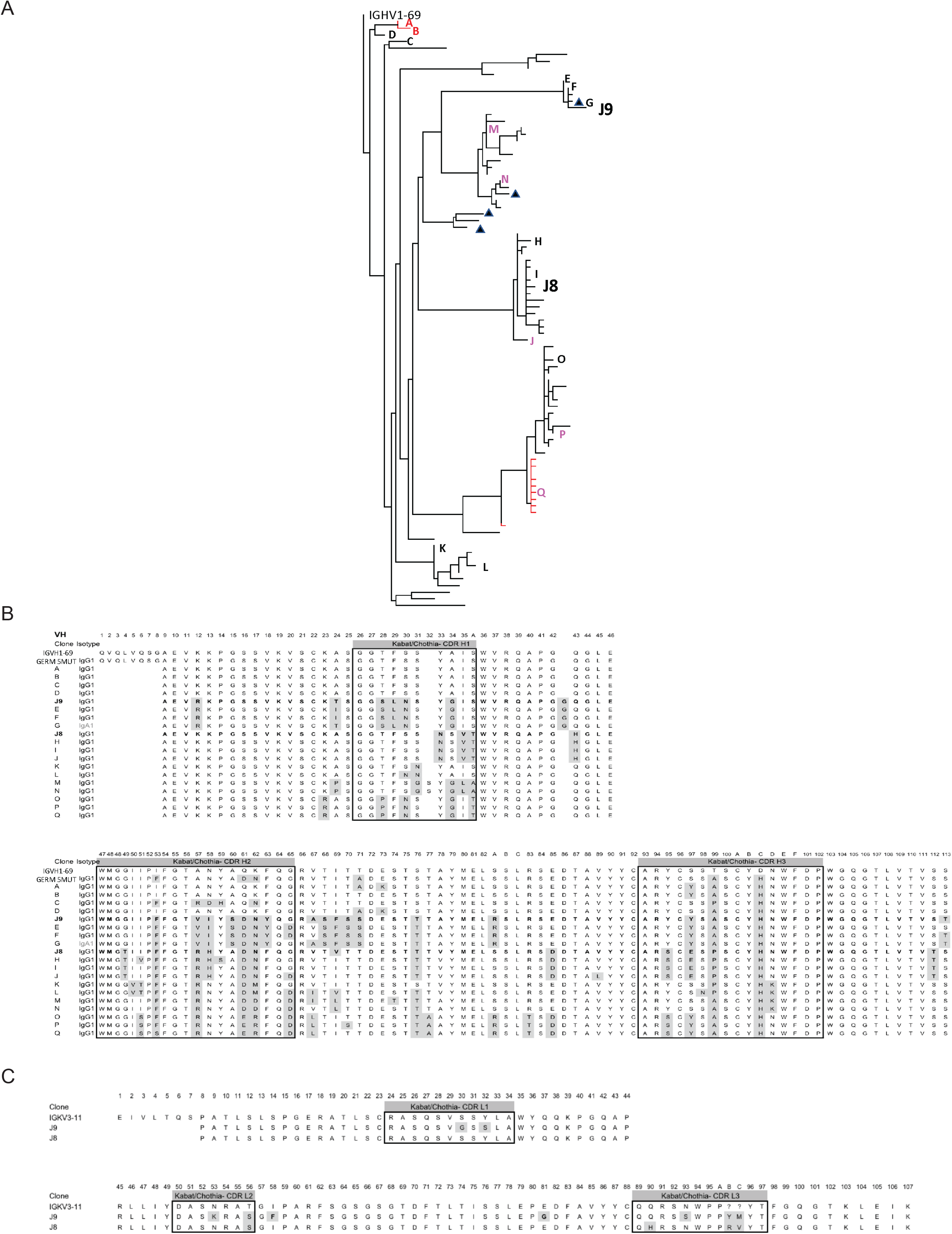
Lineage analysis of the J9/J8 clonal family. **(A)** Maximum likelihood phylogeny of mAbs related to J9 and J8 found in the BCR repertoire of patient 013 created using the HLP19 model in IgPhyML. VH germline sequence (IGHV1-69*05 + IGHD2-2 + IGHJ5) is shown at the top. Red letters and tips indicate sequences found in unstimulated PBMCs, black letters and tips indicate sequences found in stimulated PBMCs, and magenta letters indicate sequences found both in unstimulated and stimulated PBMCs. Letters correspond to sequences shown in the alignment in (B). Triangles next to tips indicate IgA sequences. (**B**) Heavy chain alignment of selected mAbs within the J9/J8 lineage found in the BCR repertoire of patient 013. Letters correspond to sequences shown in (A), with isotype indicated. The germline sequence (IGHV1-69 + IGHD2-2 + IGHJ5) is shown first, followed by a constructed sequence with 5 amino acid changes shown as “GERM 5mut.” Kabat numbering is shown on top. Amino acid changes are highlighted in gray, and boxes indicate the CDR regions. (**C**) Light chain alignment of J8 and J9. The germline sequence (IGKV3-11 + IGKJ2) is shown first, with Kabat numbering shown on top. Amino acid changes are highlighted in gray, and boxes indicate the CDR regions.

We identified clones with a 100% match at the nucleotide (nt) level to J9 and J8 in the stimulated PBMC repertoire (UMI counts of 9 and 14, respectively), and related clones identical in both unstimulated and stimulated PBMC repertoires throughout the various branches of the lineage (branch tips labeled J, Q, M, N and P in Figure 6A). Overall, the repertoire showed a rapid expansion of class switched IgG with numerous nt point mutations from germline VH, strongly suggesting both J8 (27 nt) and J9 (28 nt) plasmablasts derived from memory B cells from a prior infection, consistent with previous studies (Priyamvada et al., 2016; Xu et al., 2016). Among this acute-phase repertoire, we did observe less mutated IgG clones A (3 nt), B (5 nt), and C (5 nt) early in the lineage (Figure 6A), which could represent antibodies derived from a *de novo* immune response, or from less mutated memory clones. Finally, the divergent evolution of J9 and J8 suggested multiple somatic hypermutation pathways within this lineage leading to bNAbs.

### VH and VL maturation contributes to broadly neutralizing activity

As described above, J8 and J9 VH derived from V-D-J recombination of IGHV1-69 with IGHD2-2 and IGHJ5. Although we did not analyze the light chain repertoires, both J8 and J9 used the same founder germline IGKV3-11 and IGKJ2 genes with identical CDR lengths and no convergent mutations from germline (Figure 6C). To investigate the contribution of somatic hypermutation (SHM) on broadly neutralizing activity, we generated a panel of recombinant IgG variants, confirmed proper folding (Figure S11), and tested them for neutralizing activity. As expected, recombinant J8 and J9 IgGs expressing fully germline VH and VL had no neutralizing activity (Figures 7A-D). Similarly, J8 and J9 IgG expressing germline VH paired with the corresponding mature VL, and vice versa, did not neutralize DENV1-4, suggesting that both VH and VL SHM contributed to neutralizing activity.

**Figure 7.**
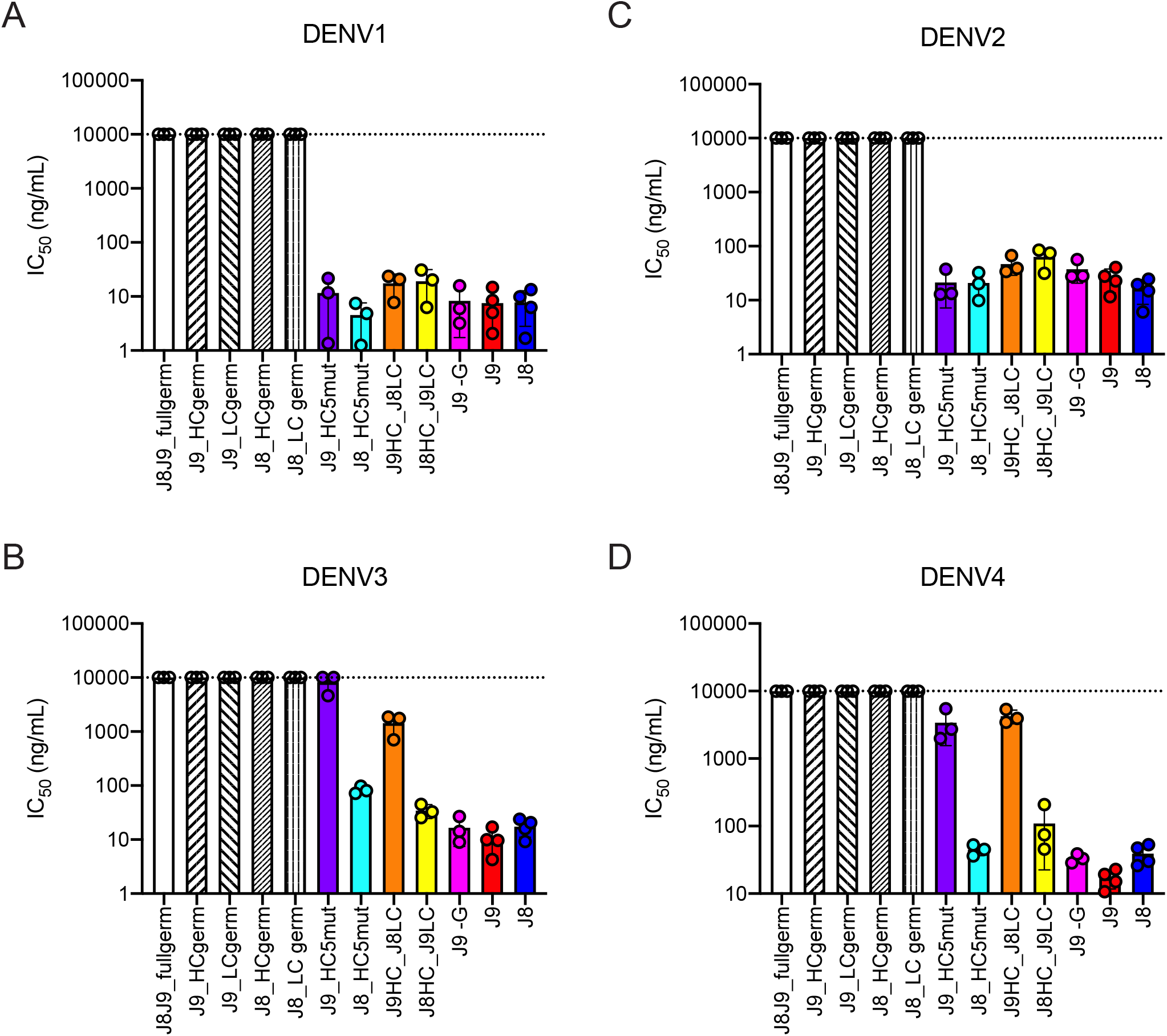
Contribution of VH and VL SHM to J9 and J8 neutralizing activity. Summary of IC_50_ values of J9 and J8 IgG variants against **(A)** DENV1, **(B)** DENV2, **(C)** DENV3, and **(D)** DENV4 RVPs. Bars represent mean IC_50_ values obtained from 3 to 4 independent experiments indicated by data points for each mAb. Error bars show the SD. Values at the dotted horizontal line indicates that 50% neutralization was not achieved at the highest IgG concentration tested (10 µg/ml). IC_50_ values for fully mature J9 and J8 are shown in red and blue bars, respectively. J8J9_full germ: germline J8/J9 heavy chain (HC) paired with J8/J9 light chain (LC); J9_HCgerm: J9 germline HC paired with J9 mature LC; J9LC_germ: J9 mature HC paired with J9 germline LC; J8_HCgerm: J8 germline HC paired with J8 mature LC; J8_LCgerm: J8 mature HC paired with J9 germline LC; J9_HC5mut: J9 HC with 5 mutations indicated in Figure 6B paired with J9 mature LC; J8_HC5mut: J8 HC with 5 mutations indicated in Figure 6B paired with J8 mature LC; J9HC_J8LC: J9 mature HC paired with J8 mature LC; J8HC_J9LC: J8 mature HC paired with J9 mature LC; J9-G: J9 HC with a single glycine deletion in FR2 paired with mature J9 LC.

Several mutations occurred early in the J8/J9 lineage, including CDR-H2 I53F, CDR-H3 T99A/P and D100cH (clones labeled A, B, and C in Figure 6A, and alignments in Figure 6B), and were retained throughout the continued VH somatic hypermutation. To investigate the VH SHM requirements for broadly neutralizing activity, we generated “J9 5mut” and “J8 5mut” variants containing the above three early VH mutations (I53F, T99P, D100cH) and two additional CDR-H2 mutations (Q61D, K62N) common across different lineage branches (Figures 6A and 6B). We paired J8 and J9 5mut VH with the corresponding mature VL to generate recombinant IgGs. These five CDR-H2 and CDR-H3 mutations were sufficient for broadly neutralizing activity of J8 against DENV1-4 (Figures 7A-D) and of J9 against DENV1 and DENV2 only (Figures 7A-B). Compared to fully mature J9, J9 5mut displayed reduced neutralization potency against DENV3 and DENV4 (Figures 7C-D), suggesting that additional J9 VH mutations were required for neutralization of these viruses. J9 VL mutations also played a role in neutralization of DENV3 and DENV4 as chimeric IgG expressing J9 heavy chain with J8 light chain displayed less potent neutralization of these viruses (Figures 7C-D). Finally, although the FR2 of J9 VH contained a glycine insertion not present in related clones (Figure 6B), this insertion was not necessary for neutralizing activity (Figure 7A-D).

## Discussion

A safe and effective vaccine to protect against DENV remains elusive, largely due to the challenge of eliciting antibodies that can potently neutralize all four viral serotypes simultaneously to minimize the risk of ADE. For antigenically diverse viruses, more than one antibody specificity may be required to provide optimal coverage of diverse circulating variants (Bell et al., 2019; Doria-Rose et al., 2012; Goo et al., 2012; Katzelnick et al., 2015; Keeffe et al., 2018; Kong et al., 2015). Although cross-reactive antibodies against flaviviruses have been described, very few display potent neutralizing activity (Barba-Spaeth et al., 2016; Dejnirattisai et al., 2015; Xu et al., 2017). The epitope for one of these bNAbs (SiGN-3C) is not well defined but involves 2 residues within DII fusion loop and one in DIII (Xu et al., 2017). Detailed epitope mapping studies have been performed only for the EDE class of bNAbs, which recognize a quaternary epitope spanning both monomers within the E protein dimer (Barba-Spaeth et al., 2016; Rouvinski et al., 2015). This epitope involves five main regions on the E protein: *b* strand, fusion loop, and *ij* loop on DII; glycan loop on DI; and the DIII A strand. In this study, we functionally characterized 28 mAbs identified to be clonally expanded and somatically hypermutated by transcriptomic analyses of single plasmablasts from two individuals acutely infected with DENV (Zanini et al., 2018). Among these, we identified bNAbs J9 and J8, which potently neutralized all four DENV serotypes and recognize an epitope with major determinants in DI. The location of residues important for J9 and J8 recognition (Figure 5A), and the ability of these bNAbs to bind virus particles but not soluble E protein (Figure 1) suggest a quaternary epitope. Alternatively, the epitope may be localized to the E monomer, but is preferentially displayed on virus particles, as previously described for a DENV1-specific mAb (Fibriansah et al., 2014). Nevertheless, our epitope mapping results demonstrated that the recognition determinants for J9/J8 are distinct from EDE bNAbs because E protein mutations that reduced neutralization potency of J9/J8 either increased or did not alter neutralization potency of EDE1 and EDE2 antibodies, respectively (Figure 5). Thus, our study defines a new vulnerable site on the DENV E protein that can be exploited for immunogen design to elicit bNAbs.

A common strategy to isolate and characterize virus-specific mAbs involves sorting hundreds of single B cells from immune donors followed by reverse-transcription (RT)-PCR to isolate paired VH/VL genes for recombinant IgG production and functional characterization (Dejnirattisai et al., 2015; Robbiani et al., 2017; Rogers et al., 2017). In some cases, memory B cells that specifically bind viral antigens are first enriched by staining with fluorescently labeled antigen (Robbiani et al., 2017; Rogers et al., 2017; Scheid et al., 2009; Woda & Mathew, 2015; Wu et al., 2010). Alternatively, single B cells are cultured, and secreted antibodies are directly screened for function (Walker et al., 2009). Although these methods have successfully identified many human bNAbs, including those against flaviviruses (Dejnirattisai et al., 2015; Smith et al., 2013; Tsai et al., 2013; Xu et al., 2012; Xu et al., 2017), they involve labor intensive steps. Instead of screening a large panel of candidate antibodies, we leveraged transcriptomic analyses of single plasmablasts from acute secondary DENV infection to focus our screen on clonally expanded and somatically hypermutated B cells (Zanini et al., 2018), which are likely to encode antigen-specific and affinity matured antibodies. Using this approach, we successfully identified highly potent mAbs capable of neutralizing all four DENV serotypes. It is unclear whether this bioinformatics-based approach to identify DENV bNAbs is fortuitous as acute DENV infection has been shown to induce a rapid and massive expansion of plasmablasts, many of which can neutralize multiple DENV serotypes (Priyamvada et al., 2016; Wrammert et al., 2012; Xu et al., 2012), or whether it is applicable to the rapid identification of highly functional antibodies against other viruses.

J9 and J8 are somatic IgG variants isolated from the same patient (013) who had an acute secondary infection with DENV4 (Zanini et al., 2018). NGS of the B cell repertoire and phylogenetic analysis of the J9/J8 lineage revealed divergent evolution of these bNAbs (Figure 6A), suggesting multiple SHM pathways to generate bNAbs against the J9/J8 epitope, which is encouraging for vaccine design. Antibody lineage divergence and parallel evolution leading to multiple bNAbs within the same individual has also been described in the context of HIV infection (MacLeod et al., 2016). J9 and J8 bNAbs derived from IGVH1-69 and IGVK3-11 germline genes. Consistent with a previous study of DENV-infected individuals (Appanna et al., 2016), many of the DENV-specific mAbs originating from plasmablasts of patient 013 also derived from IGVH1-69 (Table S3), which is commonly used among bNAbs against other viruses such as influenza and Hepatitis C virus (Chen, Tzarum, Wilson, & Law, 2019). Many of these bNAbs can achieve neutralization breadth and potency with limited SHM (Lingwood et al., 2012; Tzarum et al., 2019). Despite a moderately high degree of SHM for J8 VH (9.9% at the nucleotide level, Figure S1), five early amino acid mutations in CDR-H2 and CDR-H3 were sufficient for neutralization breadth and potency (Figure 7). As the paired VL had a low (1.4%) level of SHM, this observation suggests a relatively limited maturation pathway to a highly functional antibody.

To our knowledge, all human bNAbs against flaviviruses identified in the context of natural infection so far, including J9 and J8, were isolated from plasmablasts of individuals sequentially exposed to at least two different DENV serotypes (Dejnirattisai et al., 2015; Xu et al., 2012; Zanini et al., 2018). Sequential infection with heterologous DENV serotypes or HIV strains has been shown to broaden and strengthen the polyclonal neutralizing antibody response (Cortez, Odem-Davis, McClelland, Jaoko, & Overbaugh, 2012; Patel et al., 2017; Tsai et al., 2015; Tsai et al., 2013). One proposed model is that low affinity, cross-reactive antibody secreting B-cell clones elicited by primary DENV exposure are reactivated during secondary infection to undergo further affinity maturation resulting in antibodies with more broad and potent neutralizing activity (Patel et al., 2017). Although the relatively high level of SHM already present in J9 and J8 at day 4 post-fever onset following secondary DENV infection suggests a recall response, repertoire analysis from earlier time points would be required to determine whether the memory B cell clones from which these bNAbs were derived underwent further SHM to achieve neutralization breadth and potency.

It is also unclear whether the specificities and functions of plasmablast-derived mAbs present during acute infection confer long-lived protection from infection and pathogenesis. Interestingly, despite the presence of plasmablast-derived bNAbs such as J9 and J8, and those belonging to the EDE class during acute secondary infection, the donors from which these antibodies were isolated subsequently developed severe dengue disease (Dejnirattisai et al., 2015; Zanini et al., 2018). Indeed, the rapid and massive plasmablast activation following acute DENV infection is coincident with the onset of severe symptoms and has been proposed to contribute to immunopathology (Wrammert et al., 2012). Alternatively, despite broad and potent *in vitro* neutralizing activity against a surrogate panel of DENV1-4 strains, these bNAbs may not efficiently neutralize circulating infecting strains. Finally, it is possible that these bNAbs make up a minor component of the overall polyclonal antibody response, as supported by our finding that J9/J8-like mAbs minimally contribute to the overall neutralizing activity of patient 013 serum (Figure S9). Moreover, BCR repertoire analysis revealed that though expanded, the J9/J8 clonal family is not the largest in this donor (Table S3), at least not in the acute phase sample tested. The complex interplay among B cells and antibodies of different specificities and functions present in sera, and their impact on immunity and pathogenesis warrant further study.

## Acknowledgements

We thank the cohort participants and staff; Ted Pierson for providing Raji-DCSIGNR cells and constructs for RVP production; Anna Sellas, Gorica Margulis, Esther Ho, and Purnima Ravisankar for lab management and support; Peter Kim and Don Ganem for helpful discussion; Erick Matsen and Duncan Ralph for valuable comments on the manuscript.

Molecular graphics and analyses of the DENV2 E dimer were performed with UCSF Chimera, developed by the Resource for Biocomputing, Visualization, and Informatics at the University of California, San Francisco, with support from NIH P41-GM103311.

This work was funded by the Chan Zuckerberg Biohub (NDD, AA, EW, FZ, OS, EC, JEP, SRQ, KMM, LG); NSF Graduate Research Fellowship and the Kou-I Yeh Stanford Graduate Fellowship (DC); Catalyst Award from Dr. Ralph & Marian Falk Medical Research Trust and the Stanford Bio-X Interdisciplinary Initiatives Seed Grants Program (SE); Stanford Advanced Residency Training at Stanford Fellowship Program (MR); NIH contract HHSN272201400058C (BJD); and Fred Hutchinson Cancer Research Center (LG).

## Competing interests

FZ, DC, MR, LG, SRQ, SH, KMM, and EW are inventors of the following patent application, which is co-owned by the Chan Zuckerberg Biohub and Stanford University: PCT patent application entitled ANTIBODIES AGAINST DENGUE VIRUS AND RELATED METHODS,

Serial no. PCT/US2019/045427, filed August 7, 2019.

## Methods

### Patient samples

The study was approved by the Stanford University Administrative Panel on Human Subjects in Medical Research (Protocol #35460) and the Fundación Valle del Lili Ethics committee in biomedical research (Cali,Colombia). All subjects, their parents, or legal guardians provided written informed consent, and subjects between 6 to 17 years of age and older provided assent. We collected blood samples from individuals who presented with symptoms compatible with dengue between 2016 and 2017 to the Fundación Valle del Lili in Cali, Colombia. Cohort details have been previously described (Zanini et al., 2018).

### Monoclonal antibodies

Plasmablast-derived variable heavy or light chain sequences (Zanini et al., 2018) were synthesized as gene fragments (Genewiz, San Francisco, CA; Integrated DNA Technologies, Coralville, IA) to include at least a 15 basepair overlap with the 5’ signal sequence and 3’ constant region of our human IgG1, kappa or lambda expression vectors described elsewhere (Waltari et al., 2019). For pilot ELISAs and neutralization assays using crude IgG-containing supernatant, paired heavy and light chain plasmids for each mAb were expressed in Expi293F cells (Cat# A14527, ThermoFisher Scientific, Waltham, MA) in a 96-well format. IgG levels were quantified by ELISA as described (Waltari et al., 2019). Antibodies with crude IgG expression levels < 0.5 ng/mL were excluded from further characterization. Antibodies selected for in-depth characterization (J9, J8, C4, B10, M1, L8 and I7), as well as control mAbs EDE1 C10 (Dejnirattisai et al., 2015), EDE2 B7 (Dejnirattisai et al., 2015), CR4354 (Kaufmann et al., 2010) were expressed by transient transfection of Expi-CHO-S cells (Cat# A29129; ThermoFisher Scientific). Variable heavy and light chain sequences of the above control mAbs used for gene synthesis and cloning into expression vectors were based on PDB IDs 4UT9, 4UT6, and 3N9G, respectively. Cell culture supernatant was clarified by centrifugation at 3900 xg for 30 min at 4°C, passed through a 0.22 µm filter, and IgG was purified on MabSelect SuRe resin (Cat# 17-5438-01; GE Healthcare, Chicago, IL). Other control mAbs used in this study were obtained commercially: Anti-Dengue Virus Type II Antibody, clone 3H5-1 (Cat# MAB8702; Millipore Sigma, Burlington, MA); Flavivirus group antigen Antibody (D1-4G2-4-15 (4G2)) (Cat# NBP2-52709-0.2mg; Novus Biologicals, Centennial, CO).

### Cells

Expi293F cells (Cat# A14527; ThermoFisher Scientific) were cultured in Expi293 Expression Medium (Cat# A1435101; ThermoFisher Scientific) according to the manufacturer’s instructions. Expi-CHO-S Cells (Cat# A29127; ThermoFisher Scientific) were cultured in ExpiCHO Expression Medium (Cat# A2910001; ThermoFisher Scientific). HEK-293T/17 cells (ATCC CRL-11268) were maintained in DMEM (Cat# 11965118; ThermoFisher Scientific) supplemented with 10% fetal bovine serum (Cat# FB-11; Omega Scientific, Inc.) and 100U/mL penicillin-streptomycin (Cat# 15140-122; ThermoFisher Scientific). Raji cells stably expressing DCSIGNR (Raji-DCSIGNR) (Davis et al., 2006) (provided by Ted Pierson, NIH) and K562 cells (ATCC Cat# CCL-243) were maintained in RPMI 1640 supplemented with GlutaMAX (Cat# 72400-047; ThermoFisher Scientific), 10% FBS and 100U/mL penicillin-streptomycin. All cells were maintained at 37°C in 5% CO_2_ unless otherwise stated. C6/36 cells (ATCC CRL-1660) were maintained in EMEM (ATCC Cat# 30-2003) supplemented with 10% FBS at 30°C in 5% CO_2_.

### Production of reporter virus particles (RVPs)

RVPs were produced by co-transfection of HEK-293T/17 cells with (i) a plasmid expressing a WNV subgenomic replicon encoding GFP in place of structural genes (Pierson et al., 2006), and (ii) a plasmid encoding C-prM-E structural genes from the following viruses: DENV1 Western Pacific (WP) (Ansarah-Sobrinho, Nelson, Jost, Whitehead, & Pierson, 2008), DENV2 16681 (Ansarah-Sobrinho et al., 2008), WNV NY99 (Pierson et al., 2006), and ZIKV H/PF/2013 (Dowd et al., 2016). Briefly, 8 x 10^5 HEK-293T/17 cells pre-plated in a 6-well plate were co-transfected with a mass ratio of 1:3 replicon:C-prM-E plasmids using Lipofectamine 3000 (Cat# L3000-015; ThermoFisher Scientific). Four hours post-transfection, media was replaced with low-glucose DMEM (Cat# 12320-032; ThermoFisher Scientific) containing 10% FBS and 100U/mL penicillin-streptomycin (i.e. low-glucose DMEM complete) and cells were transferred to 30°C in 5% CO_2_. RVP-containing supernatant was harvested at 3, 4, and 5 days post-transfection, passed through a 0.22 µm filter, pooled, and stored at −80°C. DENV3 strain CH53489 RVPs (Cat# RVP-301; Integral Molecular, Philadelphia, PA) and DENV4 strain TVP360 RVPs (Cat# RVP-401; Integral Molecular) were obtained commercially. RVPs with increased efficiency of prM cleavage were produced as above by co-transfecting plasmids encoding the replicon, structural genes, and human furin (provided by Ted Pierson, NIH) at a 1:3:1 mass ratio. Where indicated, RVPs were concentrated by ultracentrifugation through 20% sucrose at 164,000 xg for 4 h at 4°C, resuspended in HNE buffer (5mM HEPES, 150mM NaCl, 0.1mM EDTA, pH 7.4), and stored at −80°C.

Infectious titers of RVPs were determined by infection of Raji-DCSIGNR cells. At 48 h post-infection, cells were fixed in 2% paraformaldehyde (Cat# 15714S; Electron Microscopy Sciences, Hatfield, PA), and GFP positive cells quantified by flow cytometry (Intellicyt iQue Screener PLUS, Sartorius AG, Gottingen, Germany).

### Production, titer and neutralization of fully infectious contemporary DENV1-4 isolates

The following DENV contemporary strains were used to infect C6/36 cells: DENV1 UIS 998 (Cat# NR-49713; BEI), DENV2 US/BID-V594/2006 (Cat# NR-43280; BEI), DENV3/US/BID-V1043/2006 (Cat# NR-43282; BEI), DENV4 Strain UIS497 (Cat# NR-49724; BEI). Virus-containing supernatant was collected at days 2-7 post-infection, filtered, and stored at −80 °C. Infectious titer was determined on Raji-DCSIGNR cells. At 48 h post-infection, intracellular staining was performed using BD Cytofix/Cytoperm Solution Kit (Cat# 554714; BD Biosciences, San Jose, CA) according to the manufacturer’s instructions. Mouse mAb 4G2 conjugated to Alexa Fluor 488 (Cat# A20181; Thermo Fisher Scientific) was used for intracellular staining to detect infected cells by flow cytometry (Intellicyt iQue Screener PLUS, Sartorius AG). Neutralization assays were performed as described below, using intracellular staining with Alexa Fluor 488-conjugated 4G2 to detect infected cells.

### Generation of E protein variants

The DENV2 16681 C-prM-E expression construct (Ansarah-Sobrinho et al., 2008) was used as a template for site-directed mutagenesis using the *Pfu* Ultra DNA polymerase system (Cat# 600380; Agilent Technologies, Santa Clara, CA) and primers generated by QuikChange® Primer Design (Agilent Technologies). The entire C-prM-E region was sequenced (Quintara, San Francisco, CA) to confirm the presence of the desired mutation(s).

### Shotgun mutagenesis epitope mapping

A DENV2 strain 16681 prM/E expression construct was subjected to high-throughput shotgun mutagenesis to generate a comprehensive mutation library, with each prM/E polyprotein residue mutated to alanine (with alanine residues to serine). In total, 559 DENV2 mutants were generated (99.6% coverage of the prM/E protein), sequence confirmed, and arrayed into 384-well plates (one mutation per well). For mAb library screening, plasmids encoding the DENV protein variants were transfected individually into human HEK-293T cells and allowed to express for 22 h before fixing cells in 4% paraformaldehyde (Electron Microscopy Sciences), and permeabilizing with 0.1% (w/v) saponin (Sigma-Aldrich, St. Louis, MA) in PBS plus calcium and magnesium (PBS++). Cells were incubated with purified mAbs (0.1-2.0 µg/mL) diluted in 10% NGS (Sigma-Aldrich) / 0.1% saponin, pH 9.0. MAb J9 was screened in unfixed cells that had been co-transfected with the prM/E library and furin expression plasmids, to decrease levels of prM in the cells. Before screening, the optimal concentration was determined for each antibody, using an independent immunofluorescence titration curve against wild-type prM/E to ensure that signals were within the linear range of detection and that signal exceeded background by at least 5-fold. Antibodies were detected using 3.75 µg/mL Alexa Fluor 488-conjugated secondary antibody (Jackson ImmunoResearch, West Grove, PA) in 10% NGS / 0.1% saponin. Cells were washed three times with PBS++ / 0.1% saponin followed by 2 washes in PBS. Mean cellular fluorescence was detected using a high throughput flow cytometer (Intellicyt iQue Screener Plus, Sartorius AG). Antibody reactivity against each mutant protein clone was calculated relative to reactivity with wild-type prM/E, by subtracting the signal from mock-transfected controls and normalizing to the signal from wild-type protein-transfected controls. The entire library data for each mAb was compared to the equivalent data from control mAbs. Mutations were identified as critical to the mAb epitope if they did not support reactivity of the test mAb (<20% of reactivity to WT prM/E) but supported reactivity of appropriate control antibodies (>70% of reactivity to WT prM/E). This counter-screen strategy facilitates the exclusion of DENV prM/E protein mutants that are mis-folded or have an expression defect (Davidson & Doranz, 2014; Paes et al., 2009).

### ELISA

High-binding 96-well plates (Cat# CLS3361; Millipore Sigma, Burlington, MA) were either coated directly with 500 ng/well of recombinant DENV2 16681 E protein (Cat# DENV2-ENV-500; The Native Antigen Company, Oxford, UK) or with 300 ng/well of murine mAb 4G2 for capture of concentrated and partially purified RVPs. Recombinant E or capture mAb was added in 100 µl 1X PBS and incubated at 4°C overnight. The following day, 300 µl 1% BSA in PBS blocking buffer (Cat# B0101; Teknova, Hollister, CA) was added for 1 h either at room temperature (RT) or 37°C. Plates were subsequently washed 6 times using 300 µl PBST (PBS + 0.05% Tween-20) and 100 µl of DENV2 RVPs diluted 1:10 in blocking buffer was added to wells coated with murine mAb 4G2 and incubated for 1 hr at RT and 37°C. Plates were washed 6 times and incubated with 5 µg/well of primary mAbs in 100µl blocking buffer for 1 h at room temperature or 37°C. Plates were washed 6 times and incubated with horseradish peroxidase (HRP)-conjugated mouse anti-human IgG Fc secondary antibody (Cat# 05-4220; ThermoFisher Scientific) diluted 1:1000 in 100 µl blocking buffer for 1 h at room temperature or 37°C. Plates were washed 6 times, and 100 μl TMB substrate (Cat# 34028; Thermo Fisher Scientific) was added at room temperature. The reaction was stopped after 6 min by adding 50 μl of 1N HCL. The absorbance at 450 nm was determined using a microplate reader (SpectraMax i3, Molecular Devices, San Jose, CA).

### Protein array printing and ELISA

Spotted protein arrays (5 x 6) were printed onto each well of Greiner high-binding 96-well plates (Cat# 655097, Thermo Fisher Scientific) using a sciFLeXARRAYER S12 (Scienion AG, Berlin). An array with 75x final concentration of sucrose-purified DENV2 RVPs and a final concentration of 180 µg/mL of recombinant DENV2 16681 E protein was printed alongside 10 µg/mL anti-human IgG Fc (Cat# 09-005-098; Jackson ImmunoResearch), and 1 µg/mL biotinylated kappa secondary (Cat# 2060-08, SouthernBiotech, Birmingham, AL). Probes were diluted 1:1 with D12 buffer (Cat# CBP-5436-25; Scienion AG) and printed at the final concentrations indicated above in triplicate spots from a 384-well source plate (Cat# CPG-5502-1; Scienion AG) chilled to dew point with 3 x 350 pL drops per spot at 60% humidity on each 96-well plate. Plates were cured overnight at 70% humidity before vacuum sealing.

For ELISAs, printed 96-well plates were washed once with binding buffer (0.5% BSA + 0.025% Tween in PBS) then blocked in 100 µl/well blocking buffer (3% BSA in PBS + 0.05% Tween-20 (PBST)). After 1 h, blocking buffer was removed and 100 µl/well of test mAbs diluted in binding buffer added and incubated overnight at 4°C. We tested twelve 3-fold serial dilutions of J9, J8, C4, EDE1 C10, EDE2 B7 and CR4354 starting at 200 µg/mL; and B10, M1, L8 and I7 starting at 2 µg/mL. The following day, plates were washed 3 times with PBST, and 100 µl/well of goat anti-human IgG Fc-BIOT (Cat# 2014-08; Southern Biotech) secondary antibody diluted 1:10,000 in binding buffer was added. After 1 h shaking incubation at room temperature, plates were washed 3 times, followed by 1 h shaking incubation at room temperature with 100 µl/well Pierce High Sensitivity Streptavidin-HRP (Cat# 21130, Thermo Fisher Scientific). Plates were again washed 3 times, developed for 20 min with 50 µl/well SciColor T12 (Cat# CD-5600-100; Scienion AG), then analyzed using sciREADER CL2 (Scienion AG). Dose-response binding curves were analyzed by non-linear regression with a variable slope (GraphPadPrism v8, GraphPad Software Inc., San Diego, CA).

### Neutralization and antibody-dependent enhancement assays

RVP stocks diluted to 5-10% final infectivity were incubated with 5-fold dilutions of mAb or heat-inactivated (65°C for 30 min) serum for 1 h at room temperature before addition of Raji-DCSIGNR cells (neutralization assays) or K562 cells (ADE assays). After 48 h incubation at 37°C, cells were fixed in 2% paraformaldehyde and GFP positive cells were quantified by flow cytometry (Intellicyt iQue Screener Plus, Sartorius AG). Dose-response neutralization curves were analyzed by non-linear regression with a variable slope (GraphPadPrism v8, GraphPad Software Inc.). Fab fragments were generated and purified from IgGs using the Pierce Fab Preparation Kit (Cat# PI44985; Thermo Scientific) and used in neutralization assays at 2x molar concentration relative to IgG.

Pre- and post-attachment neutralization assays were carried out as previously described (Xu et al., 2017) using Raji-DCSIGNR cells, DENV2 RVPs, and serial 5-fold dilutions of mAb starting at 150 µg/mL (J9 and J8) or 300 µg/mL (C4 and EDE1 C10). All cells, RVPs, mAbs, and media were pre-chilled to 4°C prior to use. For the pre-attachment assay, mAb dilutions were mixed with undiluted DENV2 RVPs for 1 h at 4°C followed by the addition of Raji-DCSIGNR cells and incubation for 1 h at 4°C. Cells were washed 3 times with media, resuspended in media and incubated for 48 h at 37°C. For the post-attachment assay, undiluted DENV2 RVPs were incubated with Raji-DCSIGNR cells for 1h at 4°C, washed three times with media, resuspended in fresh media and incubated with antibody dilutions. After 1 h at 4°C, cells were washed three times with media, resuspended in media and incubated for 48 h at 37°C. After 48 h at 37°C, cells from both pre- and post-attachment assays were fixed in 2 % paraformaldehyde. GFP positive cells were quantified by flow cytometry after 48h incubation, as described above.

### Preparation of PBMCs for BCR repertoire analysis

A 1 ml vial of PBMCs was thawed rapidly in a 37°C water bath, immediately diluted into 9 ml of B cell growth media containing Corning® DMEM [+] 4.5 g/L glucose, sodium pyruvate [-] L-glutamine (VWR International, Radnor, PA), 1x Pen/Strep/Glu and 10% ultralow IgG HI-FBS (Thermo Fisher Scientific), and pelleted at 350 xg for 5 min. The cells were resuspended in 1 mL of growth media and filtered through a 5 ml polystyrene tube with a cell strainer cap (Thomas Scientific, Swedesboro, NJ). One half of the PBMCs were transferred to the T25 flask with feeder cells and B cell stimulation media as described previously (Waltari et al., 2019) and the other half was spun down in a 1.5 ml Eppendorf tube at 8000 rpm for 5 min, resuspended in 600 µl RLT (Qiagen, Hilden, Germany) + beta-mercaptoethanol, allowed to lyse for 5 min, snap frozen on dry ice and stored at −80°C until RNA purification with the Qiagen AllPrep RNA/DNA kit (Qiagen). Immunoglobulin amplicon preparation, sequencing and BCR analysis were previously described (Waltari et al., 2019). Lineage trees were constructed using the IgPhyML package in the Immcantation pipeline (Hoehn, Fowler, Lunter, & Pybus, 2016) to construct a somatic hypermutation-optimized maximum likelihood phylogeny of the heavy chain sequences clonally related to J9 and J8.

## Supplemental Information

**Table S1.**
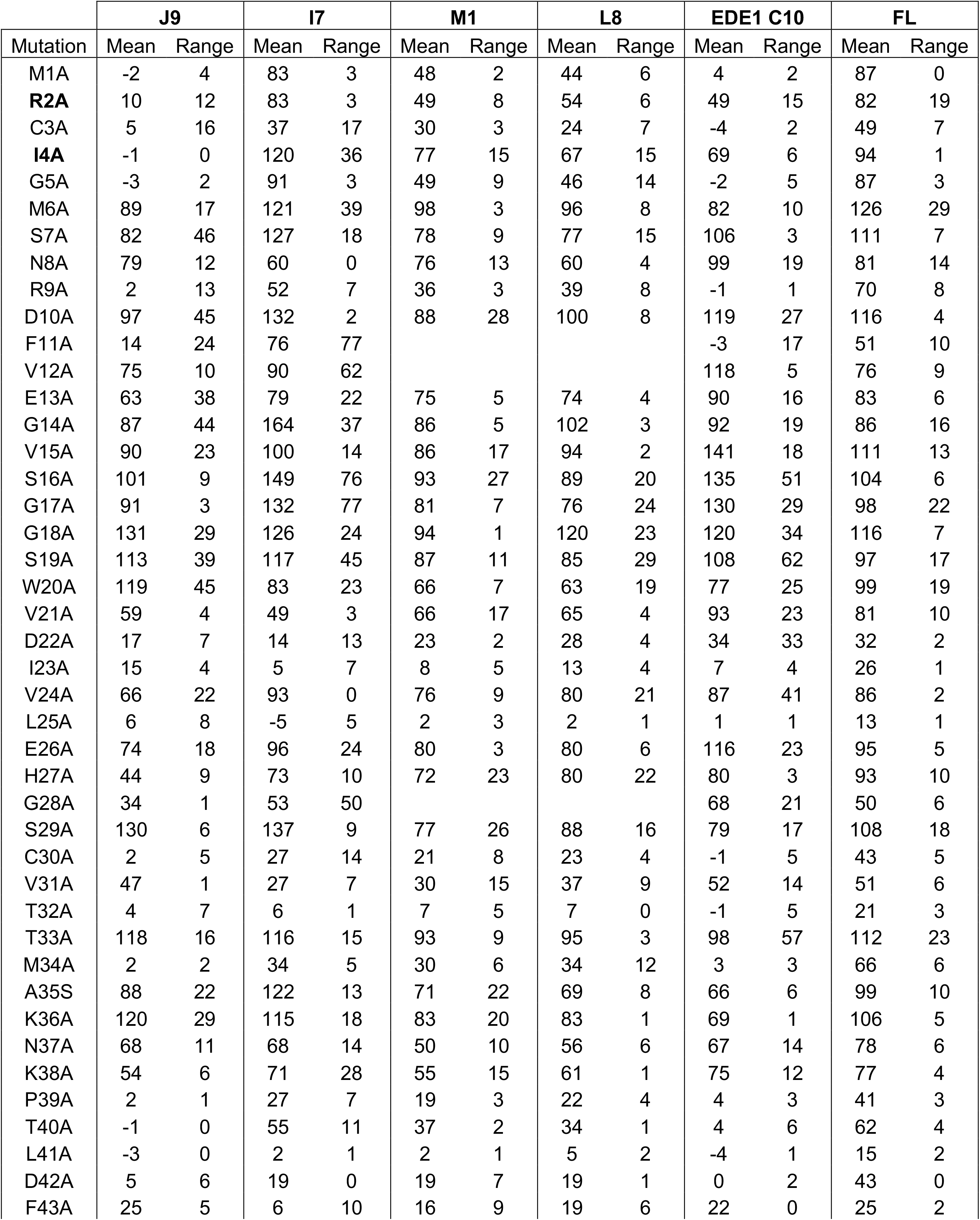
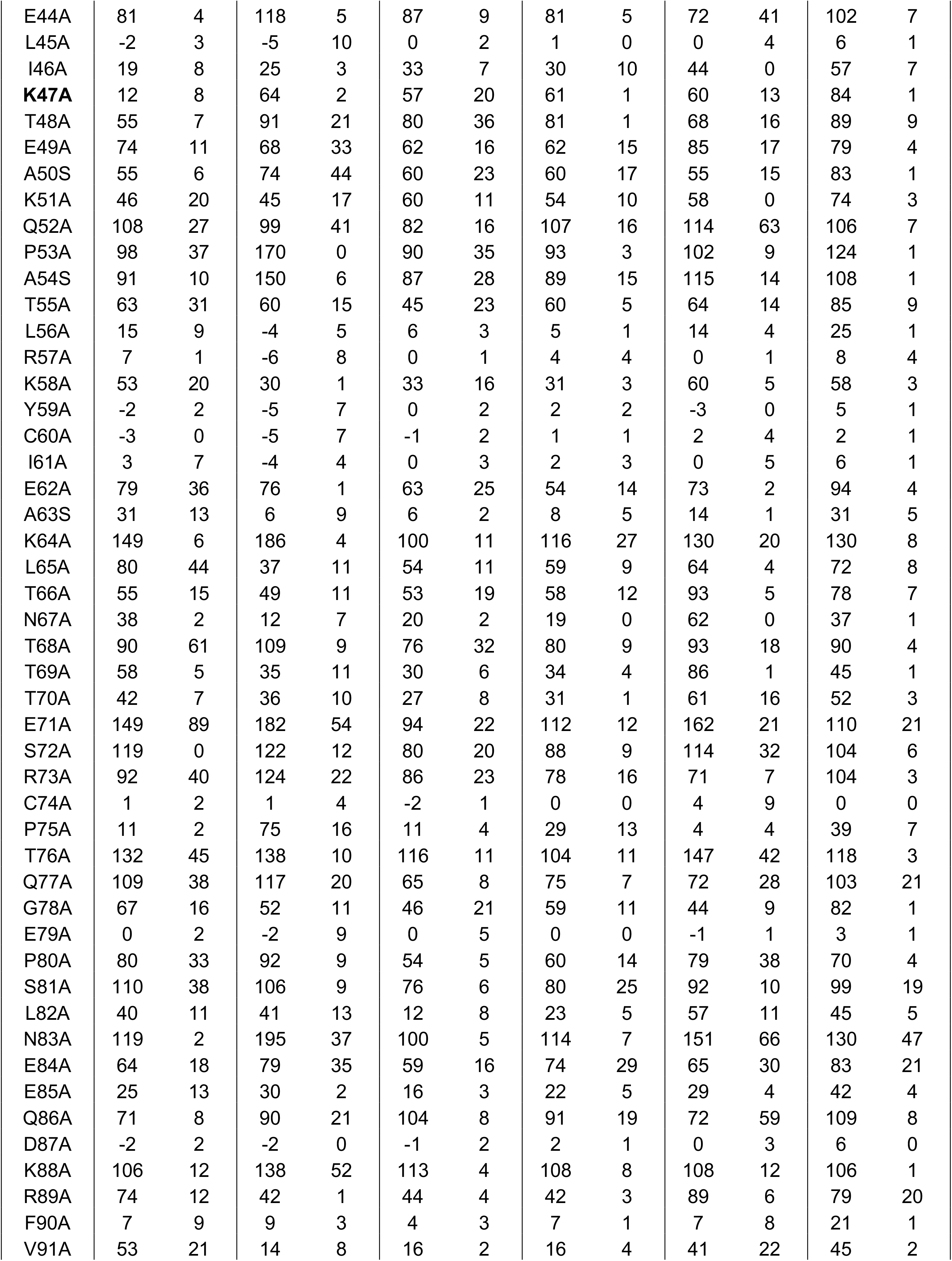
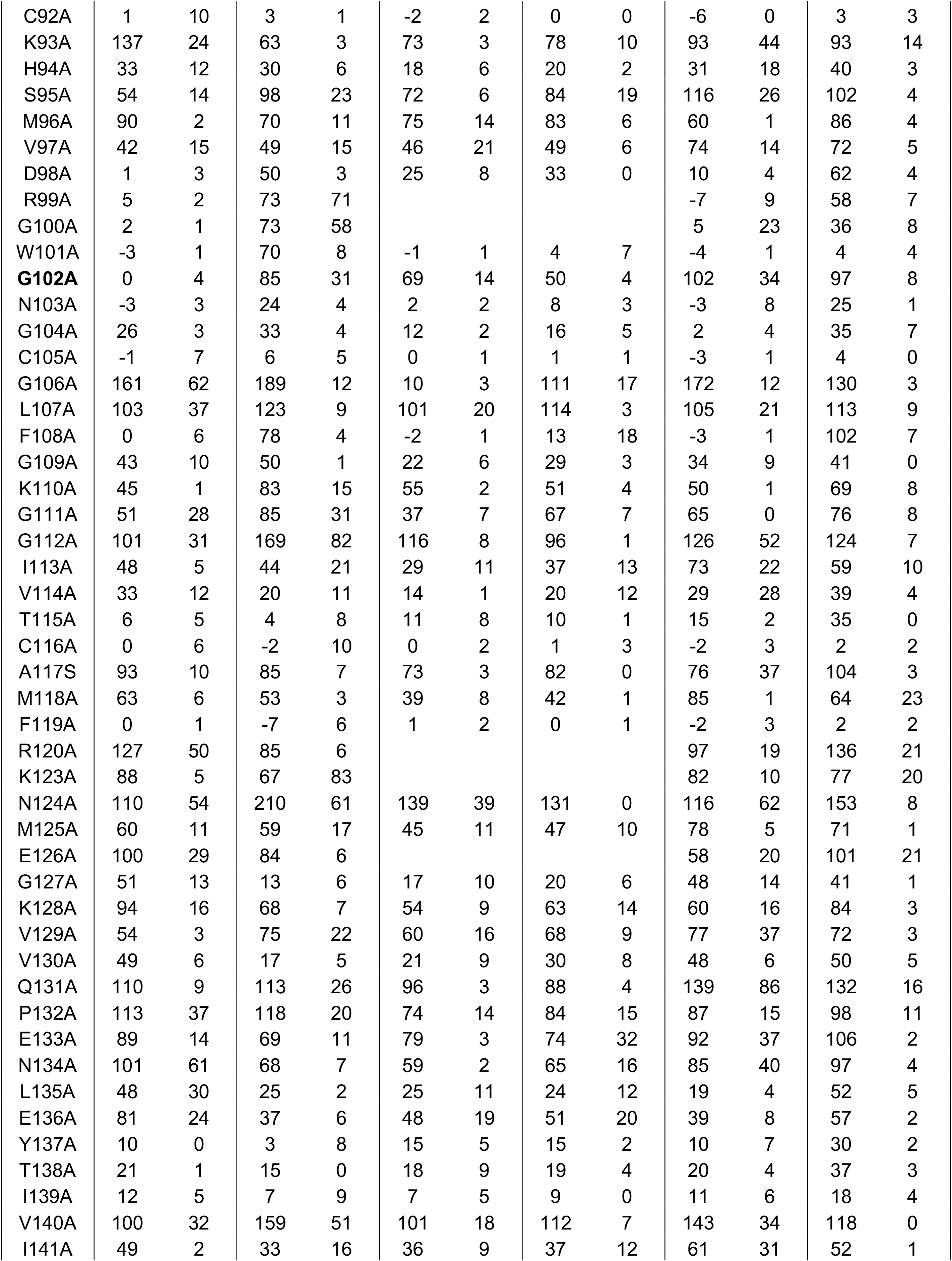
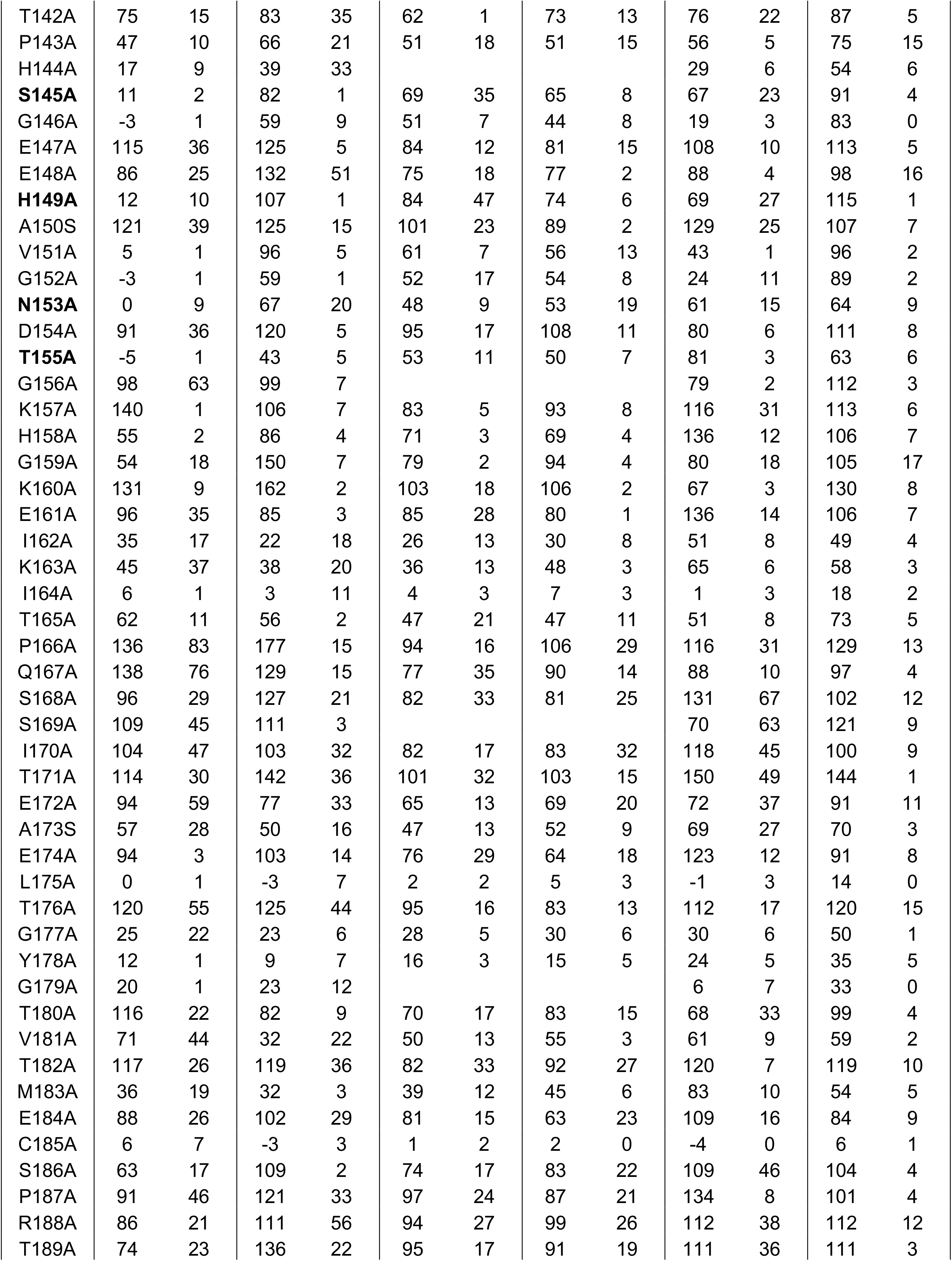
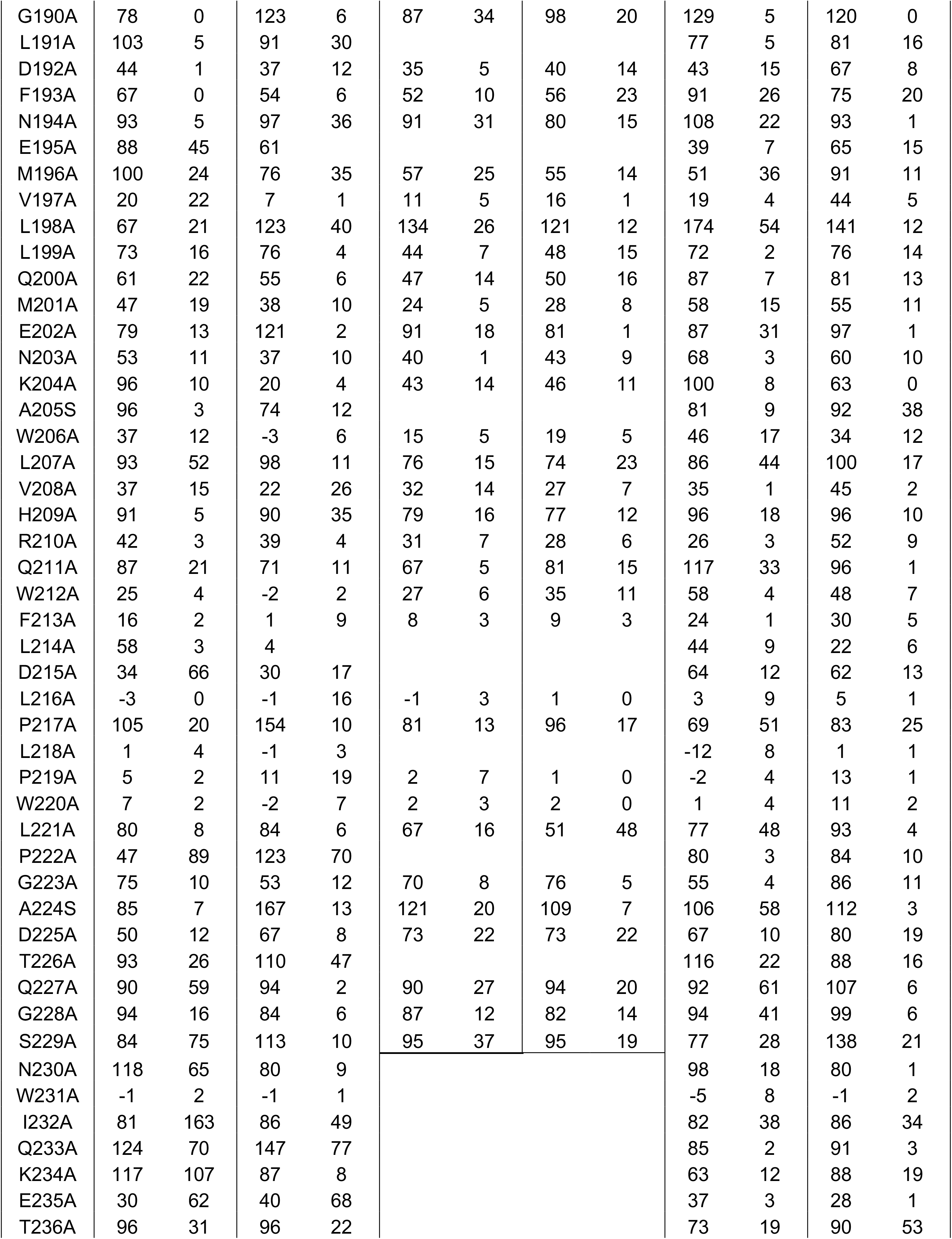
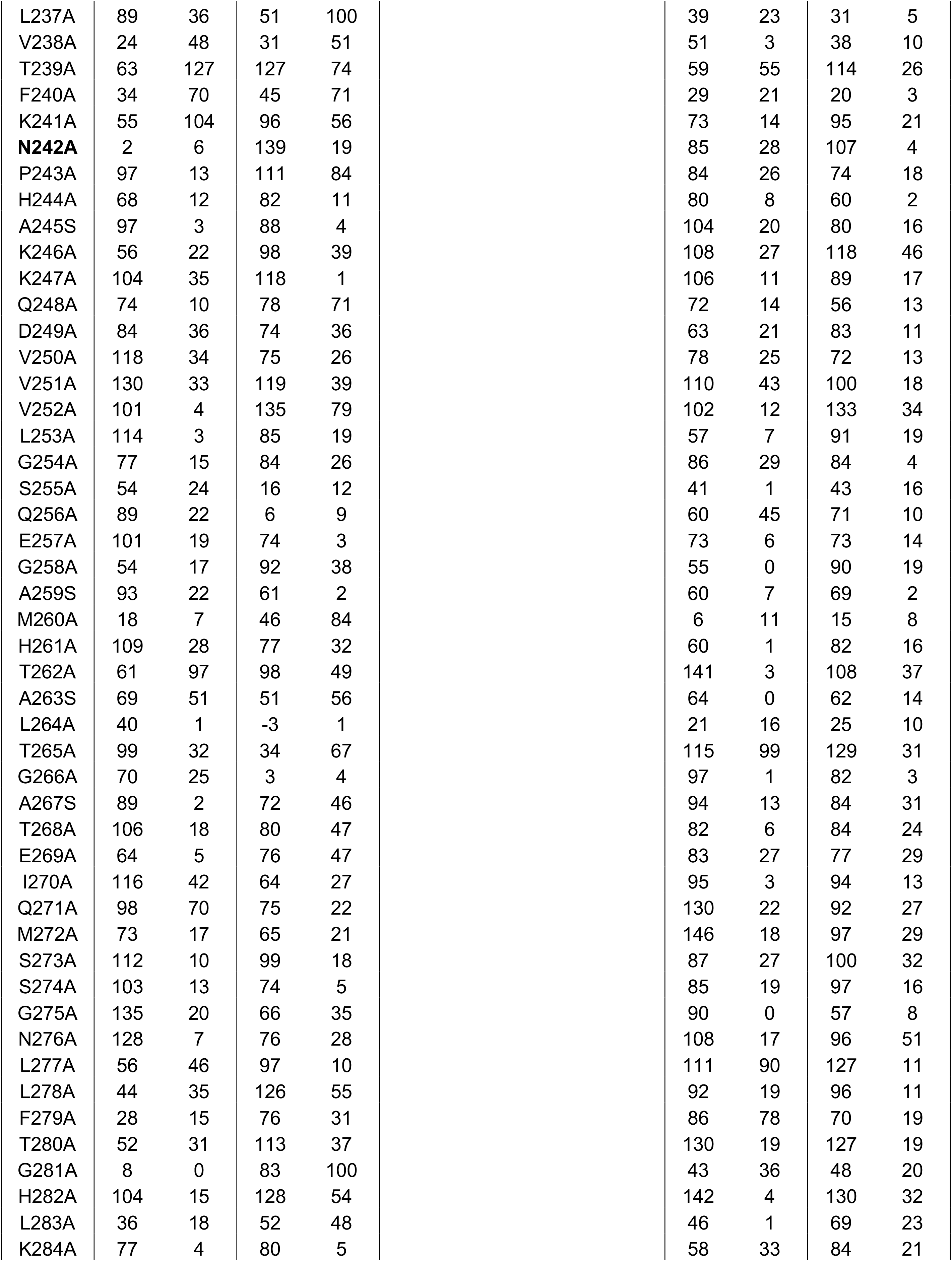
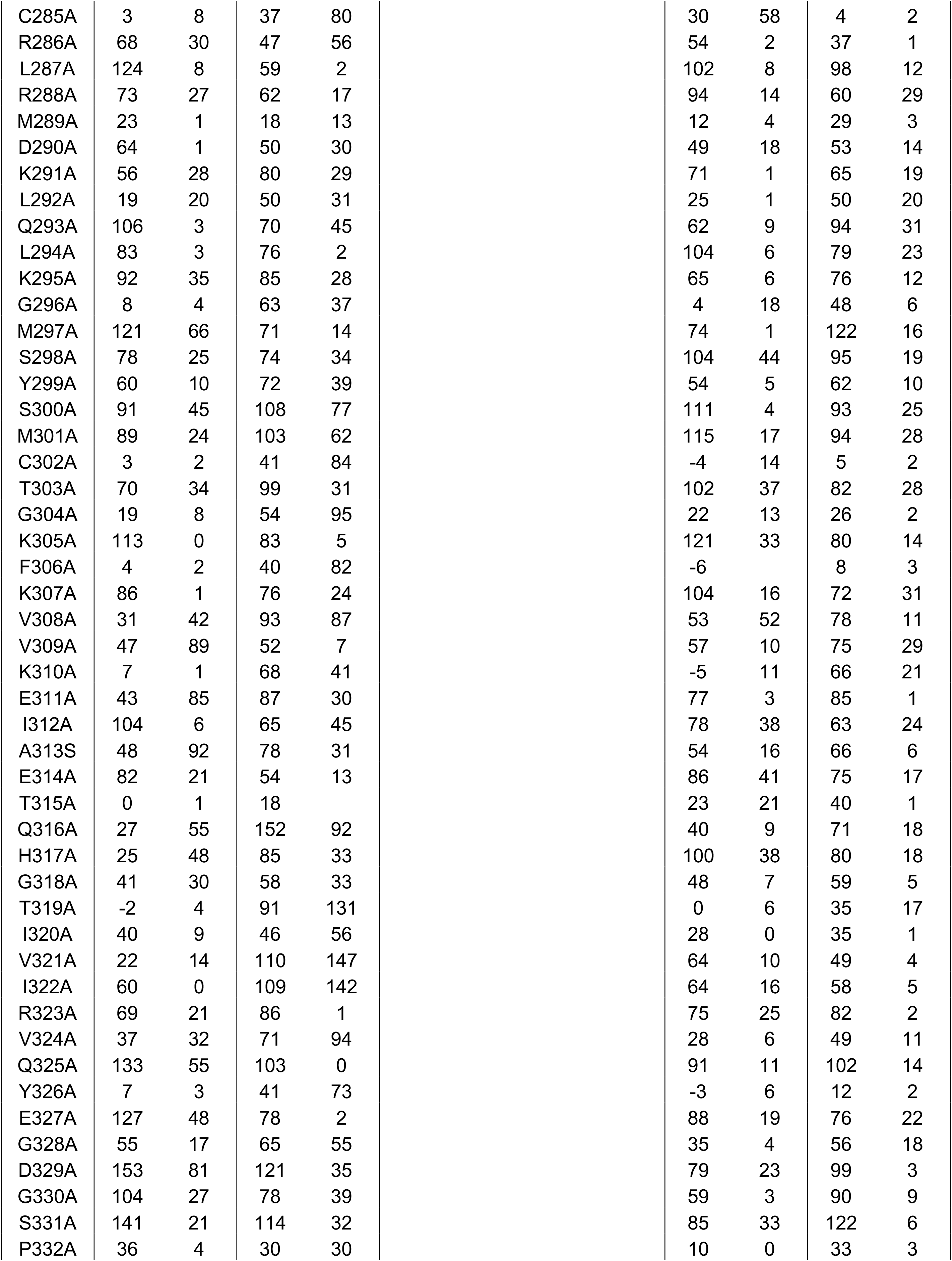
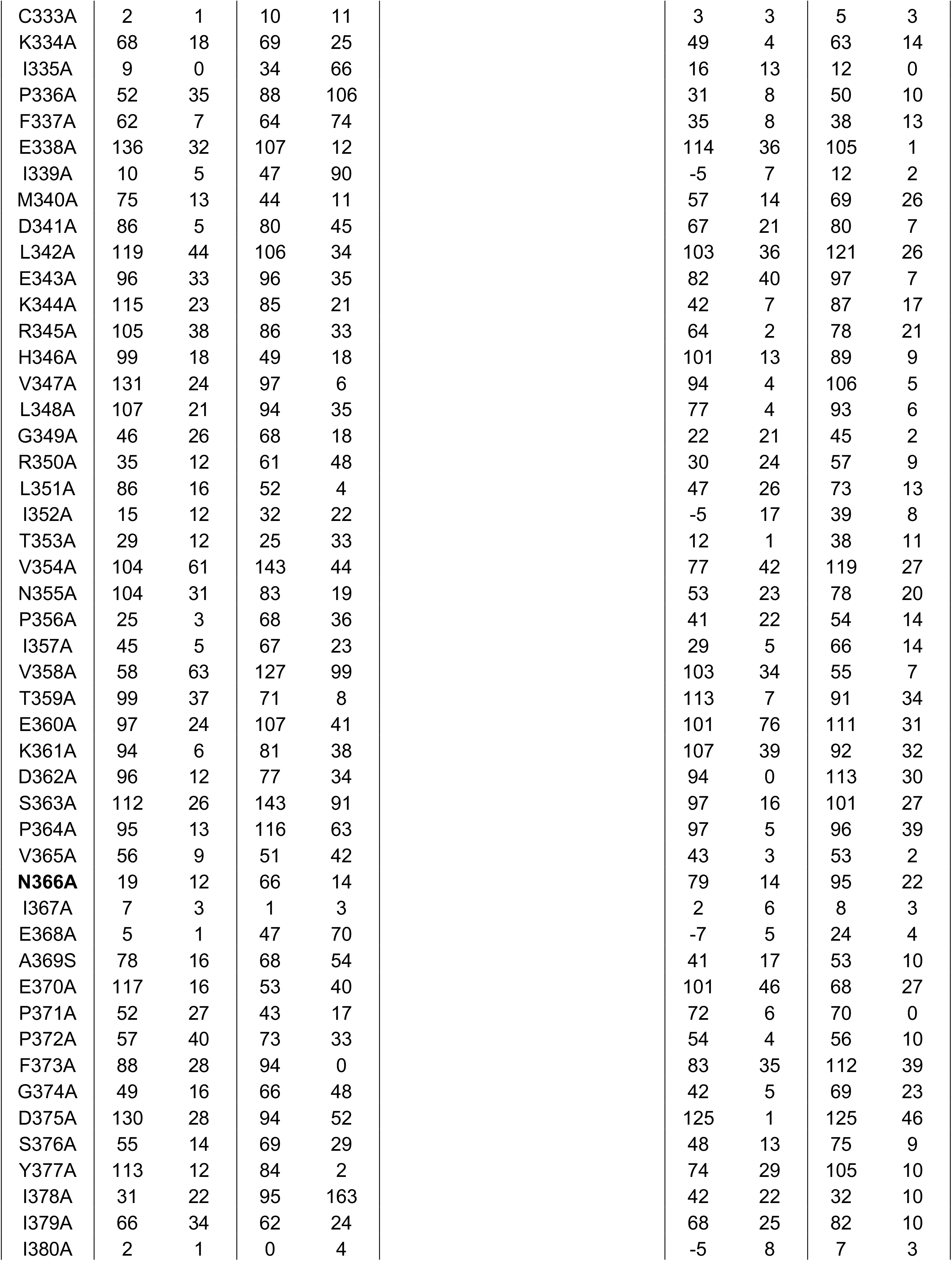
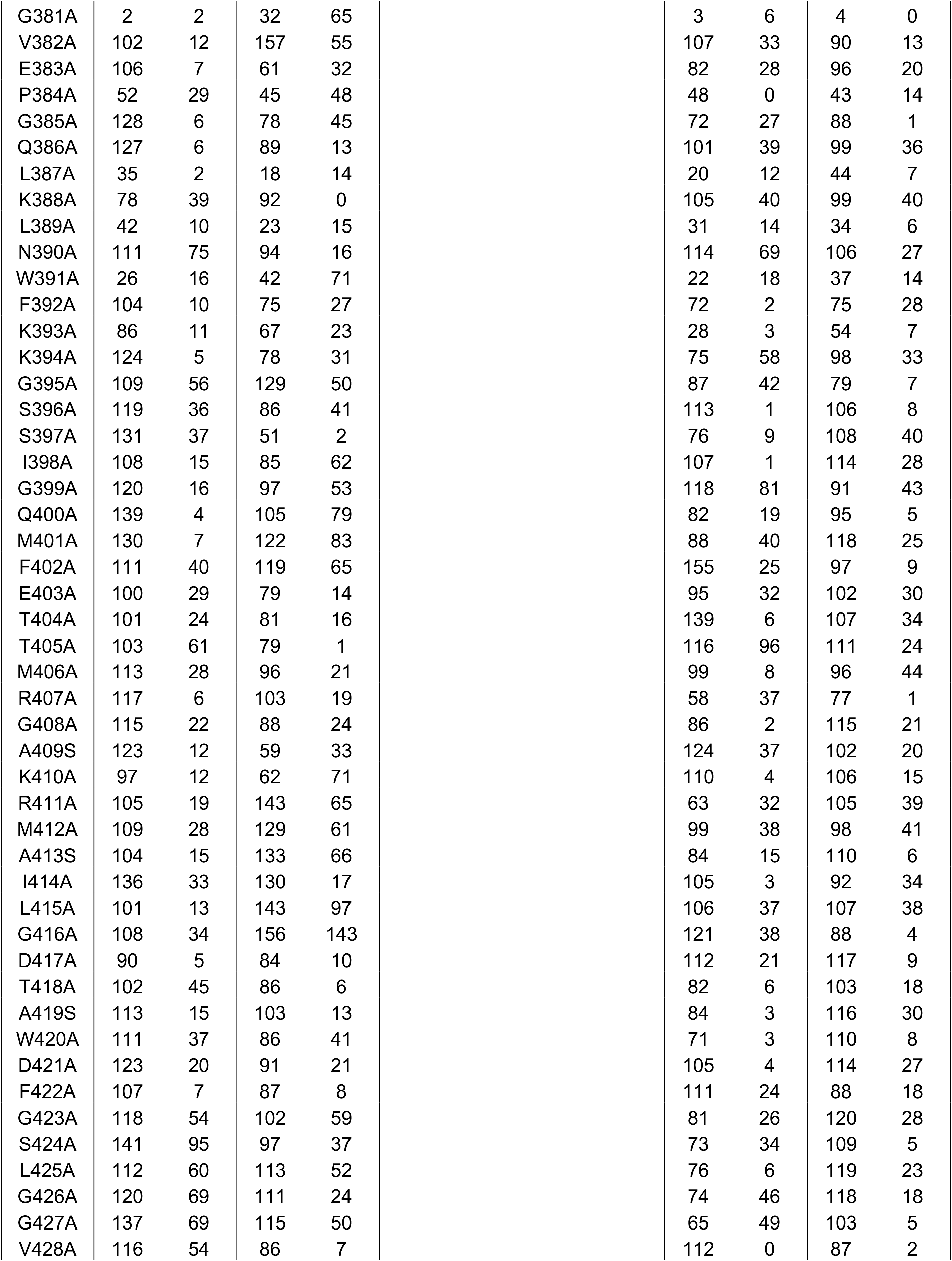
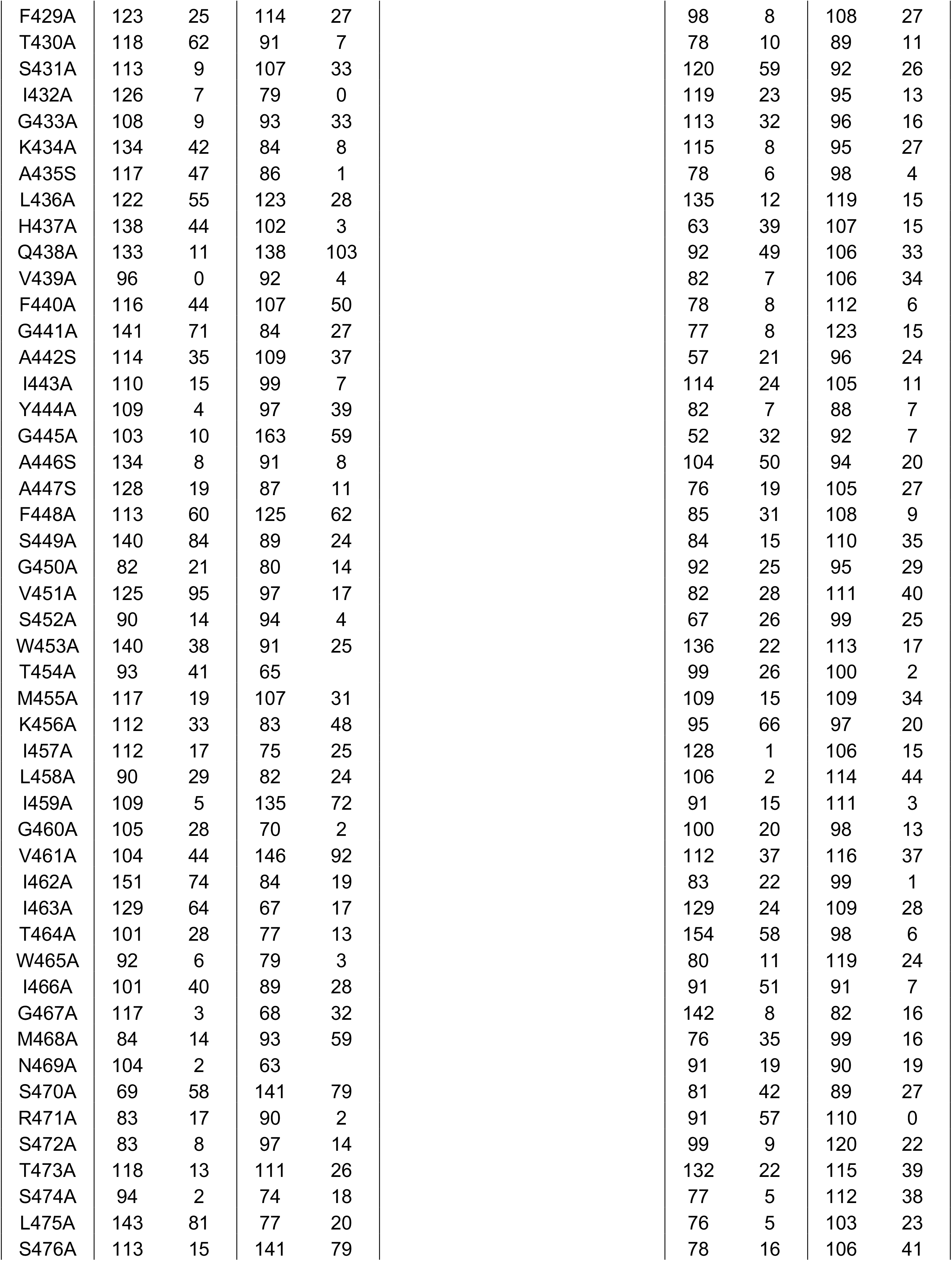
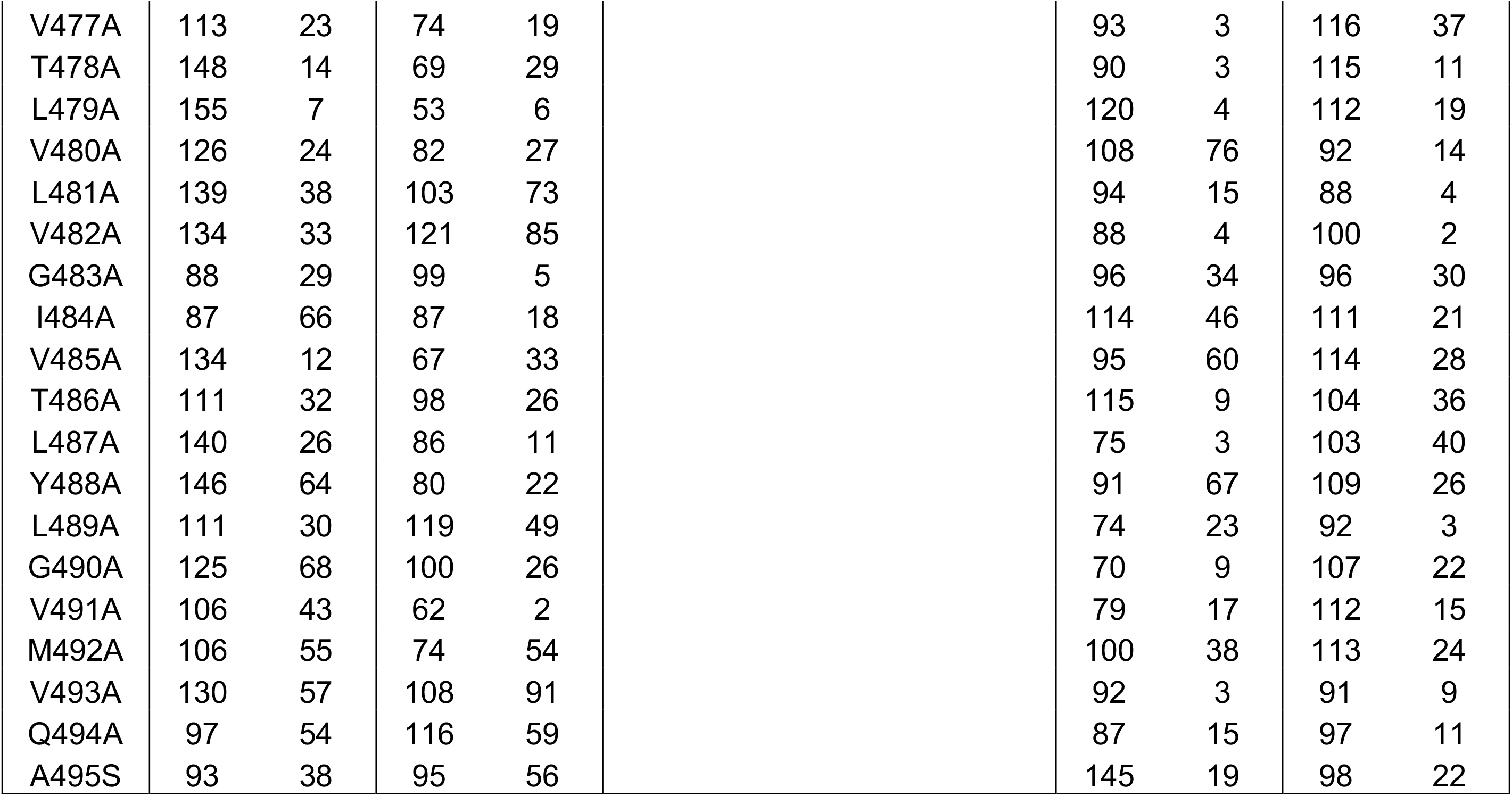
Percent mAb binding reactivity to DENV2 E protein alanine scanning mutagenesis library normalized to wildtype DENV2.

|  | **J9** |  | **I7** |  | **M1** |  | **L8** |  | **EDE1 C10** |  | **FL** |  |
| --- | --- | --- | --- | --- | --- | --- | --- | --- | --- | --- | --- | --- |
| **Mutation** | **Mean** | **Range** | **Mean** | **Range** | **Mean** | **Range** | **Mean** | **Range** | **Mean** | **Range** | **Mean** | **Range** |
| M1A | -2 | 4 | 83 | 3 | 48 | 2 | 44 | 6 | 4 | 2 | 87 | 0 |
| **R2A** | 10 | 12 | 83 | 3 | 49 | 8 | 54 | 6 | 49 | 15 | 82 | 19 |
| C3A | 5 | 16 | 37 | 17 | 30 | 3 | 24 | 7 | -4 | 2 | 49 | 7 |
| **I4A** | -1 | 0 | 120 | 36 | 77 | 15 | 67 | 15 | 69 | 6 | 94 | 1 |
| G5A | -3 | 2 | 91 | 3 | 49 | 9 | 46 | 14 | -2 | 5 | 87 | 3 |
| M6A | 89 | 17 | 121 | 39 | 98 | 3 | 96 | 8 | 82 | 10 | 126 | 29 |
| S7A | 82 | 46 | 127 | 18 | 78 | 9 | 77 | 15 | 106 | 3 | 111 | 7 |
| N8A | 79 | 12 | 60 | 0 | 76 | 13 | 60 | 4 | 99 | 19 | 81 | 14 |
| R9A | 2 | 13 | 52 | 7 | 36 | 3 | 39 | 8 | -1 | 1 | 70 | 8 |
| D10A | 97 | 45 | 132 | 2 | 88 | 28 | 100 | 8 | 119 | 27 | 116 | 4 |
| F11A | 14 | 24 | 76 | 77 |  |  |  |  | -3 | 17 | 51 | 10 |
| V12A | 75 | 10 | 90 | 62 |  |  |  |  | 118 | 5 | 76 | 9 |
| E13A | 63 | 38 | 79 | 22 | 75 | 5 | 74 | 4 | 90 | 16 | 83 | 6 |
| G14A | 87 | 44 | 164 | 37 | 86 | 5 | 102 | 3 | 92 | 19 | 86 | 16 |
| V15A | 90 | 23 | 100 | 14 | 86 | 17 | 94 | 2 | 141 | 18 | 111 | 13 |
| S16A | 101 | 9 | 149 | 76 | 93 | 27 | 89 | 20 | 135 | 51 | 104 | 6 |
| G17A | 91 | 3 | 132 | 77 | 81 | 7 | 76 | 24 | 130 | 29 | 98 | 22 |
| G18A | 131 | 29 | 126 | 24 | 94 | 1 | 120 | 23 | 120 | 34 | 116 | 7 |
| S19A | 113 | 39 | 117 | 45 | 87 | 11 | 85 | 29 | 108 | 62 | 97 | 17 |
| W20A | 119 | 45 | 83 | 23 | 66 | 7 | 63 | 19 | 77 | 25 | 99 | 19 |
| V21A | 59 | 4 | 49 | 3 | 66 | 17 | 65 | 4 | 93 | 23 | 81 | 10 |
| D22A | 17 | 7 | 14 | 13 | 23 | 2 | 28 | 4 | 34 | 33 | 32 | 2 |
| I23A | 15 | 4 | 5 | 7 | 8 | 5 | 13 | 4 | 7 | 4 | 26 | 1 |
| V24A | 66 | 22 | 93 | 0 | 76 | 9 | 80 | 21 | 87 | 41 | 86 | 2 |
| L25A | 6 | 8 | -5 | 5 | 2 | 3 | 2 | 1 | 1 | 1 | 13 | 1 |
| E26A | 74 | 18 | 96 | 24 | 80 | 3 | 80 | 6 | 116 | 23 | 95 | 5 |
| H27A | 44 | 9 | 73 | 10 | 72 | 23 | 80 | 22 | 80 | 3 | 93 | 10 |
| G28A | 34 | 1 | 53 | 50 |  |  |  |  | 68 | 21 | 50 | 6 |
| S29A | 130 | 6 | 137 | 9 | 77 | 26 | 88 | 16 | 79 | 17 | 108 | 18 |
| C30A | 2 | 5 | 27 | 14 | 21 | 8 | 23 | 4 | -1 | 5 | 43 | 5 |
| V31A | 47 | 1 | 27 | 7 | 30 | 15 | 37 | 9 | 52 | 14 | 51 | 6 |
| T32A | 4 | 7 | 6 | 1 | 7 | 5 | 7 | 0 | -1 | 5 | 21 | 3 |
| T33A | 118 | 16 | 116 | 15 | 93 | 9 | 95 | 3 | 98 | 57 | 112 | 23 |
| M34A | 2 | 2 | 34 | 5 | 30 | 6 | 34 | 12 | 3 | 3 | 66 | 6 |
| A35S | 88 | 22 | 122 | 13 | 71 | 22 | 69 | 8 | 66 | 6 | 99 | 10 |
| K36A | 120 | 29 | 115 | 18 | 83 | 20 | 83 | 1 | 69 | 1 | 106 | 5 |
| N37A | 68 | 11 | 68 | 14 | 50 | 10 | 56 | 6 | 67 | 14 | 78 | 6 |
| K38A | 54 | 6 | 71 | 28 | 55 | 15 | 61 | 1 | 75 | 12 | 77 | 4 |
| P39A | 2 | 1 | 27 | 7 | 19 | 3 | 22 | 4 | 4 | 3 | 41 | 3 |
| T40A | -1 | 0 | 55 | 11 | 37 | 2 | 34 | 1 | 4 | 6 | 62 | 4 |
| L41A | -3 | 0 | 2 | 1 | 2 | 1 | 5 | 2 | -4 | 1 | 15 | 2 |
| D42A | 5 | 6 | 19 | 0 | 19 | 7 | 19 | 1 | 0 | 2 | 43 | 0 |
| F43A | 25 | 5 | 6 | 10 | 16 | 9 | 19 | 6 | 22 | 0 | 25 | 2 |
| E44A | 81 | 4 | 118 | 5 | 87 | 9 | 81 | 5 | 72 | 41 | 102 | 7 |
| L45A | -2 | 3 | -5 | 10 | 0 | 2 | 1 | 0 | 0 | 4 | 6 | 1 |
| I46A | 19 | 8 | 25 | 3 | 33 | 7 | 30 | 10 | 44 | 0 | 57 | 7 |
| **K47A** | 12 | 8 | 64 | 2 | 57 | 20 | 61 | 1 | 60 | 13 | 84 | 1 |
| T48A | 55 | 7 | 91 | 21 | 80 | 36 | 81 | 1 | 68 | 16 | 89 | 9 |
| E49A | 74 | 11 | 68 | 33 | 62 | 16 | 62 | 15 | 85 | 17 | 79 | 4 |
| A50S | 55 | 6 | 74 | 44 | 60 | 23 | 60 | 17 | 55 | 15 | 83 | 1 |
| K51A | 46 | 20 | 45 | 17 | 60 | 11 | 54 | 10 | 58 | 0 | 74 | 3 |
| Q52A | 108 | 27 | 99 | 41 | 82 | 16 | 107 | 16 | 114 | 63 | 106 | 7 |
| P53A | 98 | 37 | 170 | 0 | 90 | 35 | 93 | 3 | 102 | 9 | 124 | 1 |
| A54S | 91 | 10 | 150 | 6 | 87 | 28 | 89 | 15 | 115 | 14 | 108 | 1 |
| T55A | 63 | 31 | 60 | 15 | 45 | 23 | 60 | 5 | 64 | 14 | 85 | 9 |
| L56A | 15 | 9 | -4 | 5 | 6 | 3 | 5 | 1 | 14 | 4 | 25 | 1 |
| R57A | 7 | 1 | -6 | 8 | 0 | 1 | 4 | 4 | 0 | 1 | 8 | 4 |
| K58A | 53 | 20 | 30 | 1 | 33 | 16 | 31 | 3 | 60 | 5 | 58 | 3 |
| Y59A | -2 | 2 | -5 | 7 | 0 | 2 | 2 | 2 | -3 | 0 | 5 | 1 |
| C60A | -3 | 0 | -5 | 7 | -1 | 2 | 1 | 1 | 2 | 4 | 2 | 1 |
| I61A | 3 | 7 | -4 | 4 | 0 | 3 | 2 | 3 | 0 | 5 | 6 | 1 |
| E62A | 79 | 36 | 76 | 1 | 63 | 25 | 54 | 14 | 73 | 2 | 94 | 4 |
| A63S | 31 | 13 | 6 | 9 | 6 | 2 | 8 | 5 | 14 | 1 | 31 | 5 |
| K64A | 149 | 6 | 186 | 4 | 100 | 11 | 116 | 27 | 130 | 20 | 130 | 8 |
| L65A | 80 | 44 | 37 | 11 | 54 | 11 | 59 | 9 | 64 | 4 | 72 | 8 |
| T66A | 55 | 15 | 49 | 11 | 53 | 19 | 58 | 12 | 93 | 5 | 78 | 7 |
| N67A | 38 | 2 | 12 | 7 | 20 | 2 | 19 | 0 | 62 | 0 | 37 | 1 |
| T68A | 90 | 61 | 109 | 9 | 76 | 32 | 80 | 9 | 93 | 18 | 90 | 4 |
| T69A | 58 | 5 | 35 | 11 | 30 | 6 | 34 | 4 | 86 | 1 | 45 | 1 |
| T70A | 42 | 7 | 36 | 10 | 27 | 8 | 31 | 1 | 61 | 16 | 52 | 3 |
| E71A | 149 | 89 | 182 | 54 | 94 | 22 | 112 | 12 | 162 | 21 | 110 | 21 |
| S72A | 119 | 0 | 122 | 12 | 80 | 20 | 88 | 9 | 114 | 32 | 104 | 6 |
| R73A | 92 | 40 | 124 | 22 | 86 | 23 | 78 | 16 | 71 | 7 | 104 | 3 |
| C74A | 1 | 2 | 1 | 4 | -2 | 1 | 0 | 0 | 4 | 9 | 0 | 0 |
| P75A | 11 | 2 | 75 | 16 | 11 | 4 | 29 | 13 | 4 | 4 | 39 | 7 |
| T76A | 132 | 45 | 138 | 10 | 116 | 11 | 104 | 11 | 147 | 42 | 118 | 3 |
| Q77A | 109 | 38 | 117 | 20 | 65 | 8 | 75 | 7 | 72 | 28 | 103 | 21 |
| G78A | 67 | 16 | 52 | 11 | 46 | 21 | 59 | 11 | 44 | 9 | 82 | 1 |
| E79A | 0 | 2 | -2 | 9 | 0 | 5 | 0 | 0 | -1 | 1 | 3 | 1 |
| P80A | 80 | 33 | 92 | 9 | 54 | 5 | 60 | 14 | 79 | 38 | 70 | 4 |
| S81A | 110 | 38 | 106 | 9 | 76 | 6 | 80 | 25 | 92 | 10 | 99 | 19 |
| L82A | 40 | 11 | 41 | 13 | 12 | 8 | 23 | 5 | 57 | 11 | 45 | 5 |
| N83A | 119 | 2 | 195 | 37 | 100 | 5 | 114 | 7 | 151 | 66 | 130 | 47 |
| E84A | 64 | 18 | 79 | 35 | 59 | 16 | 74 | 29 | 65 | 30 | 83 | 21 |
| E85A | 25 | 13 | 30 | 2 | 16 | 3 | 22 | 5 | 29 | 4 | 42 | 4 |
| Q86A | 71 | 8 | 90 | 21 | 104 | 8 | 91 | 19 | 72 | 59 | 109 | 8 |
| D87A | -2 | 2 | -2 | 0 | -1 | 2 | 2 | 1 | 0 | 3 | 6 | 0 |
| K88A | 106 | 12 | 138 | 52 | 113 | 4 | 108 | 8 | 108 | 12 | 106 | 1 |
| R89A | 74 | 12 | 42 | 1 | 44 | 4 | 42 | 3 | 89 | 6 | 79 | 20 |
| F90A | 7 | 9 | 9 | 3 | 4 | 3 | 7 | 1 | 7 | 8 | 21 | 1 |
| V91A | 53 | 21 | 14 | 8 | 16 | 2 | 16 | 4 | 41 | 22 | 45 | 2 |
| C92A | 1 | 10 | 3 | 1 | -2 | 2 | 0 | 0 | -6 | 0 | 3 | 3 |
| K93A | 137 | 24 | 63 | 3 | 73 | 3 | 78 | 10 | 93 | 44 | 93 | 14 |
| H94A | 33 | 12 | 30 | 6 | 18 | 6 | 20 | 2 | 31 | 18 | 40 | 3 |
| S95A | 54 | 14 | 98 | 23 | 72 | 6 | 84 | 19 | 116 | 26 | 102 | 4 |
| M96A | 90 | 2 | 70 | 11 | 75 | 14 | 83 | 6 | 60 | 1 | 86 | 4 |
| V97A | 42 | 15 | 49 | 15 | 46 | 21 | 49 | 6 | 74 | 14 | 72 | 5 |
| D98A | 1 | 3 | 50 | 3 | 25 | 8 | 33 | 0 | 10 | 4 | 62 | 4 |
| R99A | 5 | 2 | 73 | 71 |  |  |  |  | -7 | 9 | 58 | 7 |
| G100A | 2 | 1 | 73 | 58 |  |  |  |  | 5 | 23 | 36 | 8 |
| W101A | -3 | 1 | 70 | 8 | -1 | 1 | 4 | 7 | -4 | 1 | 4 | 4 |
| **G102A** | 0 | 4 | 85 | 31 | 69 | 14 | 50 | 4 | 102 | 34 | 97 | 8 |
| N103A | -3 | 3 | 24 | 4 | 2 | 2 | 8 | 3 | -3 | 8 | 25 | 1 |
| G104A | 26 | 3 | 33 | 4 | 12 | 2 | 16 | 5 | 2 | 4 | 35 | 7 |
| C105A | -1 | 7 | 6 | 5 | 0 | 1 | 1 | 1 | -3 | 1 | 4 | 0 |
| G106A | 161 | 62 | 189 | 12 | 10 | 3 | 111 | 17 | 172 | 12 | 130 | 3 |
| L107A | 103 | 37 | 123 | 9 | 101 | 20 | 114 | 3 | 105 | 21 | 113 | 9 |
| F108A | 0 | 6 | 78 | 4 | -2 | 1 | 13 | 18 | -3 | 1 | 102 | 7 |
| G109A | 43 | 10 | 50 | 1 | 22 | 6 | 29 | 3 | 34 | 9 | 41 | 0 |
| K110A | 45 | 1 | 83 | 15 | 55 | 2 | 51 | 4 | 50 | 1 | 69 | 8 |
| G111A | 51 | 28 | 85 | 31 | 37 | 7 | 67 | 7 | 65 | 0 | 76 | 8 |
| G112A | 101 | 31 | 169 | 82 | 116 | 8 | 96 | 1 | 126 | 52 | 124 | 7 |
| I113A | 48 | 5 | 44 | 21 | 29 | 11 | 37 | 13 | 73 | 22 | 59 | 10 |
| V114A | 33 | 12 | 20 | 11 | 14 | 1 | 20 | 12 | 29 | 28 | 39 | 4 |
| T115A | 6 | 5 | 4 | 8 | 11 | 8 | 10 | 1 | 15 | 2 | 35 | 0 |
| C116A | 0 | 6 | -2 | 10 | 0 | 2 | 1 | 3 | -2 | 3 | 2 | 2 |
| A117S | 93 | 10 | 85 | 7 | 73 | 3 | 82 | 0 | 76 | 37 | 104 | 3 |
| M118A | 63 | 6 | 53 | 3 | 39 | 8 | 42 | 1 | 85 | 1 | 64 | 23 |
| F119A | 0 | 1 | -7 | 6 | 1 | 2 | 0 | 1 | -2 | 3 | 2 | 2 |
| R120A | 127 | 50 | 85 | 6 |  |  |  |  | 97 | 19 | 136 | 21 |
| K123A | 88 | 5 | 67 | 83 |  |  |  |  | 82 | 10 | 77 | 20 |
| N124A | 110 | 54 | 210 | 61 | 139 | 39 | 131 | 0 | 116 | 62 | 153 | 8 |
| M125A | 60 | 11 | 59 | 17 | 45 | 11 | 47 | 10 | 78 | 5 | 71 | 1 |
| E126A | 100 | 29 | 84 | 6 |  |  |  |  | 58 | 20 | 101 | 21 |
| G127A | 51 | 13 | 13 | 6 | 17 | 10 | 20 | 6 | 48 | 14 | 41 | 1 |
| K128A | 94 | 16 | 68 | 7 | 54 | 9 | 63 | 14 | 60 | 16 | 84 | 3 |
| V129A | 54 | 3 | 75 | 22 | 60 | 16 | 68 | 9 | 77 | 37 | 72 | 3 |
| V130A | 49 | 6 | 17 | 5 | 21 | 9 | 30 | 8 | 48 | 6 | 50 | 5 |
| Q131A | 110 | 9 | 113 | 26 | 96 | 3 | 88 | 4 | 139 | 86 | 132 | 16 |
| P132A | 113 | 37 | 118 | 20 | 74 | 14 | 84 | 15 | 87 | 15 | 98 | 11 |
| E133A | 89 | 14 | 69 | 11 | 79 | 3 | 74 | 32 | 92 | 37 | 106 | 2 |
| N134A | 101 | 61 | 68 | 7 | 59 | 2 | 65 | 16 | 85 | 40 | 97 | 4 |
| L135A | 48 | 30 | 25 | 2 | 25 | 11 | 24 | 12 | 19 | 4 | 52 | 5 |
| E136A | 81 | 24 | 37 | 6 | 48 | 19 | 51 | 20 | 39 | 8 | 57 | 2 |
| Y137A | 10 | 0 | 3 | 8 | 15 | 5 | 15 | 2 | 10 | 7 | 30 | 2 |
| T138A | 21 | 1 | 15 | 0 | 18 | 9 | 19 | 4 | 20 | 4 | 37 | 3 |
| I139A | 12 | 5 | 7 | 9 | 7 | 5 | 9 | 0 | 11 | 6 | 18 | 4 |
| V140A | 100 | 32 | 159 | 51 | 101 | 18 | 112 | 7 | 143 | 34 | 118 | 0 |
| I141A | 49 | 2 | 33 | 16 | 36 | 9 | 37 | 12 | 61 | 31 | 52 | 1 |
| T142A | 75 | 15 | 83 | 35 | 62 | 1 | 73 | 13 | 76 | 22 | 87 | 5 |
| P143A | 47 | 10 | 66 | 21 | 51 | 18 | 51 | 15 | 56 | 5 | 75 | 15 |
| H144A | 17 | 9 | 39 | 33 |  |  |  |  | 29 | 6 | 54 | 6 |
| **S145A** | 11 | 2 | 82 | 1 | 69 | 35 | 65 | 8 | 67 | 23 | 91 | 4 |
| G146A | -3 | 1 | 59 | 9 | 51 | 7 | 44 | 8 | 19 | 3 | 83 | 0 |
| E147A | 115 | 36 | 125 | 5 | 84 | 12 | 81 | 15 | 108 | 10 | 113 | 5 |
| E148A | 86 | 25 | 132 | 51 | 75 | 18 | 77 | 2 | 88 | 4 | 98 | 16 |
| **H149A** | 12 | 10 | 107 | 1 | 84 | 47 | 74 | 6 | 69 | 27 | 115 | 1 |
| A150S | 121 | 39 | 125 | 15 | 101 | 23 | 89 | 2 | 129 | 25 | 107 | 7 |
| V151A | 5 | 1 | 96 | 5 | 61 | 7 | 56 | 13 | 43 | 1 | 96 | 2 |
| G152A | -3 | 1 | 59 | 1 | 52 | 17 | 54 | 8 | 24 | 11 | 89 | 2 |
| **N153A** | 0 | 9 | 67 | 20 | 48 | 9 | 53 | 19 | 61 | 15 | 64 | 9 |
| D154A | 91 | 36 | 120 | 5 | 95 | 17 | 108 | 11 | 80 | 6 | 111 | 8 |
| **T155A** | -5 | 1 | 43 | 5 | 53 | 11 | 50 | 7 | 81 | 3 | 63 | 6 |
| G156A | 98 | 63 | 99 | 7 |  |  |  |  | 79 | 2 | 112 | 3 |
| K157A | 140 | 1 | 106 | 7 | 83 | 5 | 93 | 8 | 116 | 31 | 113 | 6 |
| H158A | 55 | 2 | 86 | 4 | 71 | 3 | 69 | 4 | 136 | 12 | 106 | 7 |
| G159A | 54 | 18 | 150 | 7 | 79 | 2 | 94 | 4 | 80 | 18 | 105 | 17 |
| K160A | 131 | 9 | 162 | 2 | 103 | 18 | 106 | 2 | 67 | 3 | 130 | 8 |
| E161A | 96 | 35 | 85 | 3 | 85 | 28 | 80 | 1 | 136 | 14 | 106 | 7 |
| I162A | 35 | 17 | 22 | 18 | 26 | 13 | 30 | 8 | 51 | 8 | 49 | 4 |
| K163A | 45 | 37 | 38 | 20 | 36 | 13 | 48 | 3 | 65 | 6 | 58 | 3 |
| I164A | 6 | 1 | 3 | 11 | 4 | 3 | 7 | 3 | 1 | 3 | 18 | 2 |
| T165A | 62 | 11 | 56 | 2 | 47 | 21 | 47 | 11 | 51 | 8 | 73 | 5 |
| P166A | 136 | 83 | 177 | 15 | 94 | 16 | 106 | 29 | 116 | 31 | 129 | 13 |
| Q167A | 138 | 76 | 129 | 15 | 77 | 35 | 90 | 14 | 88 | 10 | 97 | 4 |
| S168A | 96 | 29 | 127 | 21 | 82 | 33 | 81 | 25 | 131 | 67 | 102 | 12 |
| S169A | 109 | 45 | 111 | 3 |  |  |  |  | 70 | 63 | 121 | 9 |
| I170A | 104 | 47 | 103 | 32 | 82 | 17 | 83 | 32 | 118 | 45 | 100 | 9 |
| T171A | 114 | 30 | 142 | 36 | 101 | 32 | 103 | 15 | 150 | 49 | 144 | 1 |
| E172A | 94 | 59 | 77 | 33 | 65 | 13 | 69 | 20 | 72 | 37 | 91 | 11 |
| A173S | 57 | 28 | 50 | 16 | 47 | 13 | 52 | 9 | 69 | 27 | 70 | 3 |
| E174A | 94 | 3 | 103 | 14 | 76 | 29 | 64 | 18 | 123 | 12 | 91 | 8 |
| L175A | 0 | 1 | -3 | 7 | 2 | 2 | 5 | 3 | -1 | 3 | 14 | 0 |
| T176A | 120 | 55 | 125 | 44 | 95 | 16 | 83 | 13 | 112 | 17 | 120 | 15 |
| G177A | 25 | 22 | 23 | 6 | 28 | 5 | 30 | 6 | 30 | 6 | 50 | 1 |
| Y178A | 12 | 1 | 9 | 7 | 16 | 3 | 15 | 5 | 24 | 5 | 35 | 5 |
| G179A | 20 | 1 | 23 | 12 |  |  |  |  | 6 | 7 | 33 | 0 |
| T180A | 116 | 22 | 82 | 9 | 70 | 17 | 83 | 15 | 68 | 33 | 99 | 4 |
| V181A | 71 | 44 | 32 | 22 | 50 | 13 | 55 | 3 | 61 | 9 | 59 | 2 |
| T182A | 117 | 26 | 119 | 36 | 82 | 33 | 92 | 27 | 120 | 7 | 119 | 10 |
| M183A | 36 | 19 | 32 | 3 | 39 | 12 | 45 | 6 | 83 | 10 | 54 | 5 |
| E184A | 88 | 26 | 102 | 29 | 81 | 15 | 63 | 23 | 109 | 16 | 84 | 9 |
| C185A | 6 | 7 | -3 | 3 | 1 | 2 | 2 | 0 | -4 | 0 | 6 | 1 |
| S186A | 63 | 17 | 109 | 2 | 74 | 17 | 83 | 22 | 109 | 46 | 104 | 4 |
| P187A | 91 | 46 | 121 | 33 | 97 | 24 | 87 | 21 | 134 | 8 | 101 | 4 |
| R188A | 86 | 21 | 111 | 56 | 94 | 27 | 99 | 26 | 112 | 38 | 112 | 12 |
| T189A | 74 | 23 | 136 | 22 | 95 | 17 | 91 | 19 | 111 | 36 | 111 | 3 |
| G190A | 78 | 0 | 123 | 6 | 87 | 34 | 98 | 20 | 129 | 5 | 120 | 0 |
| L191A | 103 | 5 | 91 | 30 |  |  |  |  | 77 | 5 | 81 | 16 |
| D192A | 44 | 1 | 37 | 12 | 35 | 5 | 40 | 14 | 43 | 15 | 67 | 8 |
| F193A | 67 | 0 | 54 | 6 | 52 | 10 | 56 | 23 | 91 | 26 | 75 | 20 |
| N194A | 93 | 5 | 97 | 36 | 91 | 31 | 80 | 15 | 108 | 22 | 93 | 1 |
| E195A | 88 | 45 | 61 |  |  |  |  |  | 39 | 7 | 65 | 15 |
| M196A | 100 | 24 | 76 | 35 | 57 | 25 | 55 | 14 | 51 | 36 | 91 | 11 |
| V197A | 20 | 22 | 7 | 1 | 11 | 5 | 16 | 1 | 19 | 4 | 44 | 5 |
| L198A | 67 | 21 | 123 | 40 | 134 | 26 | 121 | 12 | 174 | 54 | 141 | 12 |
| L199A | 73 | 16 | 76 | 4 | 44 | 7 | 48 | 15 | 72 | 2 | 76 | 14 |
| Q200A | 61 | 22 | 55 | 6 | 47 | 14 | 50 | 16 | 87 | 7 | 81 | 13 |
| M201A | 47 | 19 | 38 | 10 | 24 | 5 | 28 | 8 | 58 | 15 | 55 | 11 |
| E202A | 79 | 13 | 121 | 2 | 91 | 18 | 81 | 1 | 87 | 31 | 97 | 1 |
| N203A | 53 | 11 | 37 | 10 | 40 | 1 | 43 | 9 | 68 | 3 | 60 | 10 |
| K204A | 96 | 10 | 20 | 4 | 43 | 14 | 46 | 11 | 100 | 8 | 63 | 0 |
| A205S | 96 | 3 | 74 | 12 |  |  |  |  | 81 | 9 | 92 | 38 |
| W206A | 37 | 12 | -3 | 6 | 15 | 5 | 19 | 5 | 46 | 17 | 34 | 12 |
| L207A | 93 | 52 | 98 | 11 | 76 | 15 | 74 | 23 | 86 | 44 | 100 | 17 |
| V208A | 37 | 15 | 22 | 26 | 32 | 14 | 27 | 7 | 35 | 1 | 45 | 2 |
| H209A | 91 | 5 | 90 | 35 | 79 | 16 | 77 | 12 | 96 | 18 | 96 | 10 |
| R210A | 42 | 3 | 39 | 4 | 31 | 7 | 28 | 6 | 26 | 3 | 52 | 9 |
| Q211A | 87 | 21 | 71 | 11 | 67 | 5 | 81 | 15 | 117 | 33 | 96 | 1 |
| W212A | 25 | 4 | -2 | 2 | 27 | 6 | 35 | 11 | 58 | 4 | 48 | 7 |
| F213A | 16 | 2 | 1 | 9 | 8 | 3 | 9 | 3 | 24 | 1 | 30 | 5 |
| L214A | 58 | 3 | 4 |  |  |  |  |  | 44 | 9 | 22 | 6 |
| D215A | 34 | 66 | 30 | 17 |  |  |  |  | 64 | 12 | 62 | 13 |
| L216A | -3 | 0 | -1 | 16 | -1 | 3 | 1 | 0 | 3 | 9 | 5 | 1 |
| P217A | 105 | 20 | 154 | 10 | 81 | 13 | 96 | 17 | 69 | 51 | 83 | 25 |
| L218A | 1 | 4 | -1 | 3 |  |  |  |  | -12 | 8 | 1 | 1 |
| P219A | 5 | 2 | 11 | 19 | 2 | 7 | 1 | 0 | -2 | 4 | 13 | 1 |
| W220A | 7 | 2 | -2 | 7 | 2 | 3 | 2 | 0 | 1 | 4 | 11 | 2 |
| L221A | 80 | 8 | 84 | 6 | 67 | 16 | 51 | 48 | 77 | 48 | 93 | 4 |
| P222A | 47 | 89 | 123 | 70 |  |  |  |  | 80 | 3 | 84 | 10 |
| G223A | 75 | 10 | 53 | 12 | 70 | 8 | 76 | 5 | 55 | 4 | 86 | 11 |
| A224S | 85 | 7 | 167 | 13 | 121 | 20 | 109 | 7 | 106 | 58 | 112 | 3 |
| D225A | 50 | 12 | 67 | 8 | 73 | 22 | 73 | 22 | 67 | 10 | 80 | 19 |
| T226A | 93 | 26 | 110 | 47 |  |  |  |  | 116 | 22 | 88 | 16 |
| Q227A | 90 | 59 | 94 | 2 | 90 | 27 | 94 | 20 | 92 | 61 | 107 | 6 |
| G228A | 94 | 16 | 84 | 6 | 87 | 12 | 82 | 14 | 94 | 41 | 99 | 6 |
| S229A | 84 | 75 | 113 | 10 | 95 | 37 | 95 | 19 | 77 | 28 | 138 | 21 |
| N230A | 118 | 65 | 80 | 9 |  |  |  |  | 98 | 18 | 80 | 1 |
| W231A | -1 | 2 | -1 | 1 |  |  |  |  | -5 | 8 | -1 | 2 |
| I232A | 81 | 163 | 86 | 49 |  |  |  |  | 82 | 38 | 86 | 34 |
| Q233A | 124 | 70 | 147 | 77 |  |  |  |  | 85 | 2 | 91 | 3 |
| K234A | 117 | 107 | 87 | 8 |  |  |  |  | 63 | 12 | 88 | 19 |
| E235A | 30 | 62 | 40 | 68 |  |  |  |  | 37 | 3 | 28 | 1 |
| T236A | 96 | 31 | 96 | 22 |  |  |  |  | 73 | 19 | 90 | 53 |

|  |  |  |  |  |  |  |  |  |
| --- | --- | --- | --- | --- | --- | --- | --- | --- |
| L237A | 89 | 36 | 51 | 100 | 39 | 23 | 31 | 5 |
| V238A | 24 | 48 | 31 | 51 | 51 | 3 | 38 | 10 |
| T239A | 63 | 127 | 127 | 74 | 59 | 55 | 114 | 26 |
| F240A | 34 | 70 | 45 | 71 | 29 | 21 | 20 | 3 |
| K241A | 55 | 104 | 96 | 56 | 73 | 14 | 95 | 21 |
| **N242A** | 2 | 6 | 139 | 19 | 85 | 28 | 107 | 4 |
| P243A | 97 | 13 | 111 | 84 | 84 | 26 | 74 | 18 |
| H244A | 68 | 12 | 82 | 11 | 80 | 8 | 60 | 2 |
| A245S | 97 | 3 | 88 | 4 | 104 | 20 | 80 | 16 |
| K246A | 56 | 22 | 98 | 39 | 108 | 27 | 118 | 46 |
| K247A | 104 | 35 | 118 | 1 | 106 | 11 | 89 | 17 |
| Q248A | 74 | 10 | 78 | 71 | 72 | 14 | 56 | 13 |
| D249A | 84 | 36 | 74 | 36 | 63 | 21 | 83 | 11 |
| V250A | 118 | 34 | 75 | 26 | 78 | 25 | 72 | 13 |
| V251A | 130 | 33 | 119 | 39 | 110 | 43 | 100 | 18 |
| V252A | 101 | 4 | 135 | 79 | 102 | 12 | 133 | 34 |
| L253A | 114 | 3 | 85 | 19 | 57 | 7 | 91 | 19 |
| G254A | 77 | 15 | 84 | 26 | 86 | 29 | 84 | 4 |
| S255A | 54 | 24 | 16 | 12 | 41 | 1 | 43 | 16 |
| Q256A | 89 | 22 | 6 | 9 | 60 | 45 | 71 | 10 |
| E257A | 101 | 19 | 74 | 3 | 73 | 6 | 73 | 14 |
| G258A | 54 | 17 | 92 | 38 | 55 | 0 | 90 | 19 |
| A259S | 93 | 22 | 61 | 2 | 60 | 7 | 69 | 2 |
| M260A | 18 | 7 | 46 | 84 | 6 | 11 | 15 | 8 |
| H261A | 109 | 28 | 77 | 32 | 60 | 1 | 82 | 16 |
| T262A | 61 | 97 | 98 | 49 | 141 | 3 | 108 | 37 |
| A263S | 69 | 51 | 51 | 56 | 64 | 0 | 62 | 14 |
| L264A | 40 | 1 | -3 | 1 | 21 | 16 | 25 | 10 |
| T265A | 99 | 32 | 34 | 67 | 115 | 99 | 129 | 31 |
| G266A | 70 | 25 | 3 | 4 | 97 | 1 | 82 | 3 |
| A267S | 89 | 2 | 72 | 46 | 94 | 13 | 84 | 31 |
| T268A | 106 | 18 | 80 | 47 | 82 | 6 | 84 | 24 |
| E269A | 64 | 5 | 76 | 47 | 83 | 27 | 77 | 29 |
| I270A | 116 | 42 | 64 | 27 | 95 | 3 | 94 | 13 |
| Q271A | 98 | 70 | 75 | 22 | 130 | 22 | 92 | 27 |
| M272A | 73 | 17 | 65 | 21 | 146 | 18 | 97 | 29 |
| S273A | 112 | 10 | 99 | 18 | 87 | 27 | 100 | 32 |
| S274A | 103 | 13 | 74 | 5 | 85 | 19 | 97 | 16 |
| G275A | 135 | 20 | 66 | 35 | 90 | 0 | 57 | 8 |
| N276A | 128 | 7 | 76 | 28 | 108 | 17 | 96 | 51 |
| L277A | 56 | 46 | 97 | 10 | 111 | 90 | 127 | 11 |
| L278A | 44 | 35 | 126 | 55 | 92 | 19 | 96 | 11 |
| F279A | 28 | 15 | 76 | 31 | 86 | 78 | 70 | 19 |
| T280A | 52 | 31 | 113 | 37 | 130 | 19 | 127 | 19 |
| G281A | 8 | 0 | 83 | 100 | 43 | 36 | 48 | 20 |
| H282A | 104 | 15 | 128 | 54 | 142 | 4 | 130 | 32 |
| L283A | 36 | 18 | 52 | 48 | 46 | 1 | 69 | 23 |
| K284A | 77 | 4 | 80 | 5 | 58 | 33 | 84 | 21 |
| C285A | 3 | 8 | 37 | 80 | 30 | 58 | 4 | 2 |
| R286A | 68 | 30 | 47 | 56 | 54 | 2 | 37 | 1 |
| L287A | 124 | 8 | 59 | 2 | 102 | 8 | 98 | 12 |
| R288A | 73 | 27 | 62 | 17 | 94 | 14 | 60 | 29 |
| M289A | 23 | 1 | 18 | 13 | 12 | 4 | 29 | 3 |
| D290A | 64 | 1 | 50 | 30 | 49 | 18 | 53 | 14 |
| K291A | 56 | 28 | 80 | 29 | 71 | 1 | 65 | 19 |
| L292A | 19 | 20 | 50 | 31 | 25 | 1 | 50 | 20 |
| Q293A | 106 | 3 | 70 | 45 | 62 | 9 | 94 | 31 |
| L294A | 83 | 3 | 76 | 2 | 104 | 6 | 79 | 23 |
| K295A | 92 | 35 | 85 | 28 | 65 | 6 | 76 | 12 |
| G296A | 8 | 4 | 63 | 37 | 4 | 18 | 48 | 6 |
| M297A | 121 | 66 | 71 | 14 | 74 | 1 | 122 | 16 |
| S298A | 78 | 25 | 74 | 34 | 104 | 44 | 95 | 19 |
| Y299A | 60 | 10 | 72 | 39 | 54 | 5 | 62 | 10 |
| S300A | 91 | 45 | 108 | 77 | 111 | 4 | 93 | 25 |
| M301A | 89 | 24 | 103 | 62 | 115 | 17 | 94 | 28 |
| C302A | 3 | 2 | 41 | 84 | -4 | 14 | 5 | 2 |
| T303A | 70 | 34 | 99 | 31 | 102 | 37 | 82 | 28 |
| G304A | 19 | 8 | 54 | 95 | 22 | 13 | 26 | 2 |
| K305A | 113 | 0 | 83 | 5 | 121 | 33 | 80 | 14 |
| F306A | 4 | 2 | 40 | 82 | -6 |  | 8 | 3 |
| K307A | 86 | 1 | 76 | 24 | 104 | 16 | 72 | 31 |
| V308A | 31 | 42 | 93 | 87 | 53 | 52 | 78 | 11 |
| V309A | 47 | 89 | 52 | 7 | 57 | 10 | 75 | 29 |
| K310A | 7 | 1 | 68 | 41 | -5 | 11 | 66 | 21 |
| E311A | 43 | 85 | 87 | 30 | 77 | 3 | 85 | 1 |
| I312A | 104 | 6 | 65 | 45 | 78 | 38 | 63 | 24 |
| A313S | 48 | 92 | 78 | 31 | 54 | 16 | 66 | 6 |
| E314A | 82 | 21 | 54 | 13 | 86 | 41 | 75 | 17 |
| T315A | 0 | 1 | 18 |  | 23 | 21 | 40 | 1 |
| Q316A | 27 | 55 | 152 | 92 | 40 | 9 | 71 | 18 |
| H317A | 25 | 48 | 85 | 33 | 100 | 38 | 80 | 18 |
| G318A | 41 | 30 | 58 | 33 | 48 | 7 | 59 | 5 |
| T319A | -2 | 4 | 91 | 131 | 0 | 6 | 35 | 17 |
| I320A | 40 | 9 | 46 | 56 | 28 | 0 | 35 | 1 |
| V321A | 22 | 14 | 110 | 147 | 64 | 10 | 49 | 4 |
| I322A | 60 | 0 | 109 | 142 | 64 | 16 | 58 | 5 |
| R323A | 69 | 21 | 86 | 1 | 75 | 25 | 82 | 2 |
| V324A | 37 | 32 | 71 | 94 | 28 | 6 | 49 | 11 |
| Q325A | 133 | 55 | 103 | 0 | 91 | 11 | 102 | 14 |
| Y326A | 7 | 3 | 41 | 73 | -3 | 6 | 12 | 2 |
| E327A | 127 | 48 | 78 | 2 | 88 | 19 | 76 | 22 |
| G328A | 55 | 17 | 65 | 55 | 35 | 4 | 56 | 18 |
| D329A | 153 | 81 | 121 | 35 | 79 | 23 | 99 | 3 |
| G330A | 104 | 27 | 78 | 39 | 59 | 3 | 90 | 9 |
| S331A | 141 | 21 | 114 | 32 | 85 | 33 | 122 | 6 |
| P332A | 36 | 4 | 30 | 30 | 10 | 0 | 33 | 3 |
| C333A | 2 | 1 | 10 | 11 | 3 | 3 | 5 | 3 |
| K334A | 68 | 18 | 69 | 25 | 49 | 4 | 63 | 14 |
| I335A | 9 | 0 | 34 | 66 | 16 | 13 | 12 | 0 |
| P336A | 52 | 35 | 88 | 106 | 31 | 8 | 50 | 10 |
| F337A | 62 | 7 | 64 | 74 | 35 | 8 | 38 | 13 |
| E338A | 136 | 32 | 107 | 12 | 114 | 36 | 105 | 1 |
| I339A | 10 | 5 | 47 | 90 | -5 | 7 | 12 | 2 |
| M340A | 75 | 13 | 44 | 11 | 57 | 14 | 69 | 26 |
| D341A | 86 | 5 | 80 | 45 | 67 | 21 | 80 | 7 |
| L342A | 119 | 44 | 106 | 34 | 103 | 36 | 121 | 26 |
| E343A | 96 | 33 | 96 | 35 | 82 | 40 | 97 | 7 |
| K344A | 115 | 23 | 85 | 21 | 42 | 7 | 87 | 17 |
| R345A | 105 | 38 | 86 | 33 | 64 | 2 | 78 | 21 |
| H346A | 99 | 18 | 49 | 18 | 101 | 13 | 89 | 9 |
| V347A | 131 | 24 | 97 | 6 | 94 | 4 | 106 | 5 |
| L348A | 107 | 21 | 94 | 35 | 77 | 4 | 93 | 6 |
| G349A | 46 | 26 | 68 | 18 | 22 | 21 | 45 | 2 |
| R350A | 35 | 12 | 61 | 48 | 30 | 24 | 57 | 9 |
| L351A | 86 | 16 | 52 | 4 | 47 | 26 | 73 | 13 |
| I352A | 15 | 12 | 32 | 22 | -5 | 17 | 39 | 8 |
| T353A | 29 | 12 | 25 | 33 | 12 | 1 | 38 | 11 |
| V354A | 104 | 61 | 143 | 44 | 77 | 42 | 119 | 27 |
| N355A | 104 | 31 | 83 | 19 | 53 | 23 | 78 | 20 |
| P356A | 25 | 3 | 68 | 36 | 41 | 22 | 54 | 14 |
| I357A | 45 | 5 | 67 | 23 | 29 | 5 | 66 | 14 |
| V358A | 58 | 63 | 127 | 99 | 103 | 34 | 55 | 7 |
| T359A | 99 | 37 | 71 | 8 | 113 | 7 | 91 | 34 |
| E360A | 97 | 24 | 107 | 41 | 101 | 76 | 111 | 31 |
| K361A | 94 | 6 | 81 | 38 | 107 | 39 | 92 | 32 |
| D362A | 96 | 12 | 77 | 34 | 94 | 0 | 113 | 30 |
| S363A | 112 | 26 | 143 | 91 | 97 | 16 | 101 | 27 |
| P364A | 95 | 13 | 116 | 63 | 97 | 5 | 96 | 39 |
| V365A | 56 | 9 | 51 | 42 | 43 | 3 | 53 | 2 |
| **N366A** | 19 | 12 | 66 | 14 | 79 | 14 | 95 | 22 |
| I367A | 7 | 3 | 1 | 3 | 2 | 6 | 8 | 3 |
| E368A | 5 | 1 | 47 | 70 | -7 | 5 | 24 | 4 |
| A369S | 78 | 16 | 68 | 54 | 41 | 17 | 53 | 10 |
| E370A | 117 | 16 | 53 | 40 | 101 | 46 | 68 | 27 |
| P371A | 52 | 27 | 43 | 17 | 72 | 6 | 70 | 0 |
| P372A | 57 | 40 | 73 | 33 | 54 | 4 | 56 | 10 |
| F373A | 88 | 28 | 94 | 0 | 83 | 35 | 112 | 39 |
| G374A | 49 | 16 | 66 | 48 | 42 | 5 | 69 | 23 |
| D375A | 130 | 28 | 94 | 52 | 125 | 1 | 125 | 46 |
| S376A | 55 | 14 | 69 | 29 | 48 | 13 | 75 | 9 |
| Y377A | 113 | 12 | 84 | 2 | 74 | 29 | 105 | 10 |
| I378A | 31 | 22 | 95 | 163 | 42 | 22 | 32 | 10 |
| I379A | 66 | 34 | 62 | 24 | 68 | 25 | 82 | 10 |
| I380A | 2 | 1 | 0 | 4 | -5 | 8 | 7 | 3 |
| G381A | 2 | 2 | 32 | 65 | 3 | 6 | 4 | 0 |
| V382A | 102 | 12 | 157 | 55 | 107 | 33 | 90 | 13 |
| E383A | 106 | 7 | 61 | 32 | 82 | 28 | 96 | 20 |
| P384A | 52 | 29 | 45 | 48 | 48 | 0 | 43 | 14 |
| G385A | 128 | 6 | 78 | 45 | 72 | 27 | 88 | 1 |
| Q386A | 127 | 6 | 89 | 13 | 101 | 39 | 99 | 36 |
| L387A | 35 | 2 | 18 | 14 | 20 | 12 | 44 | 7 |
| K388A | 78 | 39 | 92 | 0 | 105 | 40 | 99 | 40 |
| L389A | 42 | 10 | 23 | 15 | 31 | 14 | 34 | 6 |
| N390A | 111 | 75 | 94 | 16 | 114 | 69 | 106 | 27 |
| W391A | 26 | 16 | 42 | 71 | 22 | 18 | 37 | 14 |
| F392A | 104 | 10 | 75 | 27 | 72 | 2 | 75 | 28 |
| K393A | 86 | 11 | 67 | 23 | 28 | 3 | 54 | 7 |
| K394A | 124 | 5 | 78 | 31 | 75 | 58 | 98 | 33 |
| G395A | 109 | 56 | 129 | 50 | 87 | 42 | 79 | 7 |
| S396A | 119 | 36 | 86 | 41 | 113 | 1 | 106 | 8 |
| S397A | 131 | 37 | 51 | 2 | 76 | 9 | 108 | 40 |
| I398A | 108 | 15 | 85 | 62 | 107 | 1 | 114 | 28 |
| G399A | 120 | 16 | 97 | 53 | 118 | 81 | 91 | 43 |
| Q400A | 139 | 4 | 105 | 79 | 82 | 19 | 95 | 5 |
| M401A | 130 | 7 | 122 | 83 | 88 | 40 | 118 | 25 |
| F402A | 111 | 40 | 119 | 65 | 155 | 25 | 97 | 9 |
| E403A | 100 | 29 | 79 | 14 | 95 | 32 | 102 | 30 |
| T404A | 101 | 24 | 81 | 16 | 139 | 6 | 107 | 34 |
| T405A | 103 | 61 | 79 | 1 | 116 | 96 | 111 | 24 |
| M406A | 113 | 28 | 96 | 21 | 99 | 8 | 96 | 44 |
| R407A | 117 | 6 | 103 | 19 | 58 | 37 | 77 | 1 |
| G408A | 115 | 22 | 88 | 24 | 86 | 2 | 115 | 21 |
| A409S | 123 | 12 | 59 | 33 | 124 | 37 | 102 | 20 |
| K410A | 97 | 12 | 62 | 71 | 110 | 4 | 106 | 15 |
| R411A | 105 | 19 | 143 | 65 | 63 | 32 | 105 | 39 |
| M412A | 109 | 28 | 129 | 61 | 99 | 38 | 98 | 41 |
| A413S | 104 | 15 | 133 | 66 | 84 | 15 | 110 | 6 |
| I414A | 136 | 33 | 130 | 17 | 105 | 3 | 92 | 34 |
| L415A | 101 | 13 | 143 | 97 | 106 | 37 | 107 | 38 |
| G416A | 108 | 34 | 156 | 143 | 121 | 38 | 88 | 4 |
| D417A | 90 | 5 | 84 | 10 | 112 | 21 | 117 | 9 |
| T418A | 102 | 45 | 86 | 6 | 82 | 6 | 103 | 18 |
| A419S | 113 | 15 | 103 | 13 | 84 | 3 | 116 | 30 |
| W420A | 111 | 37 | 86 | 41 | 71 | 3 | 110 | 8 |
| D421A | 123 | 20 | 91 | 21 | 105 | 4 | 114 | 27 |
| F422A | 107 | 7 | 87 | 8 | 111 | 24 | 88 | 18 |
| G423A | 118 | 54 | 102 | 59 | 81 | 26 | 120 | 28 |
| S424A | 141 | 95 | 97 | 37 | 73 | 34 | 109 | 5 |
| L425A | 112 | 60 | 113 | 52 | 76 | 6 | 119 | 23 |
| G426A | 120 | 69 | 111 | 24 | 74 | 46 | 118 | 18 |
| G427A | 137 | 69 | 115 | 50 | 65 | 49 | 103 | 5 |
| V428A | 116 | 54 | 86 | 7 | 112 | 0 | 87 | 2 |
| F429A | 123 | 25 | 114 | 27 | 98 | 8 | 108 | 27 |
| T430A | 118 | 62 | 91 | 7 | 78 | 10 | 89 | 11 |
| S431A | 113 | 9 | 107 | 33 | 120 | 59 | 92 | 26 |
| I432A | 126 | 7 | 79 | 0 | 119 | 23 | 95 | 13 |
| G433A | 108 | 9 | 93 | 33 | 113 | 32 | 96 | 16 |
| K434A | 134 | 42 | 84 | 8 | 115 | 8 | 95 | 27 |
| A435S | 117 | 47 | 86 | 1 | 78 | 6 | 98 | 4 |
| L436A | 122 | 55 | 123 | 28 | 135 | 12 | 119 | 15 |
| H437A | 138 | 44 | 102 | 3 | 63 | 39 | 107 | 15 |
| Q438A | 133 | 11 | 138 | 103 | 92 | 49 | 106 | 33 |
| V439A | 96 | 0 | 92 | 4 | 82 | 7 | 106 | 34 |
| F440A | 116 | 44 | 107 | 50 | 78 | 8 | 112 | 6 |
| G441A | 141 | 71 | 84 | 27 | 77 | 8 | 123 | 15 |
| A442S | 114 | 35 | 109 | 37 | 57 | 21 | 96 | 24 |
| I443A | 110 | 15 | 99 | 7 | 114 | 24 | 105 | 11 |
| Y444A | 109 | 4 | 97 | 39 | 82 | 7 | 88 | 7 |
| G445A | 103 | 10 | 163 | 59 | 52 | 32 | 92 | 7 |
| A446S | 134 | 8 | 91 | 8 | 104 | 50 | 94 | 20 |
| A447S | 128 | 19 | 87 | 11 | 76 | 19 | 105 | 27 |
| F448A | 113 | 60 | 125 | 62 | 85 | 31 | 108 | 9 |
| S449A | 140 | 84 | 89 | 24 | 84 | 15 | 110 | 35 |
| G450A | 82 | 21 | 80 | 14 | 92 | 25 | 95 | 29 |
| V451A | 125 | 95 | 97 | 17 | 82 | 28 | 111 | 40 |
| S452A | 90 | 14 | 94 | 4 | 67 | 26 | 99 | 25 |
| W453A | 140 | 38 | 91 | 25 | 136 | 22 | 113 | 17 |
| T454A | 93 | 41 | 65 |  | 99 | 26 | 100 | 2 |
| M455A | 117 | 19 | 107 | 31 | 109 | 15 | 109 | 34 |
| K456A | 112 | 33 | 83 | 48 | 95 | 66 | 97 | 20 |
| I457A | 112 | 17 | 75 | 25 | 128 | 1 | 106 | 15 |
| L458A | 90 | 29 | 82 | 24 | 106 | 2 | 114 | 44 |
| I459A | 109 | 5 | 135 | 72 | 91 | 15 | 111 | 3 |
| G460A | 105 | 28 | 70 | 2 | 100 | 20 | 98 | 13 |
| V461A | 104 | 44 | 146 | 92 | 112 | 37 | 116 | 37 |
| I462A | 151 | 74 | 84 | 19 | 83 | 22 | 99 | 1 |
| I463A | 129 | 64 | 67 | 17 | 129 | 24 | 109 | 28 |
| T464A | 101 | 28 | 77 | 13 | 154 | 58 | 98 | 6 |
| W465A | 92 | 6 | 79 | 3 | 80 | 11 | 119 | 24 |
| I466A | 101 | 40 | 89 | 28 | 91 | 51 | 91 | 7 |
| G467A | 117 | 3 | 68 | 32 | 142 | 8 | 82 | 16 |
| M468A | 84 | 14 | 93 | 59 | 76 | 35 | 99 | 16 |
| N469A | 104 | 2 | 63 |  | 91 | 19 | 90 | 19 |
| S470A | 69 | 58 | 141 | 79 | 81 | 42 | 89 | 27 |
| R471A | 83 | 17 | 90 | 2 | 91 | 57 | 110 | 0 |
| S472A | 83 | 8 | 97 | 14 | 99 | 9 | 120 | 22 |
| T473A | 118 | 13 | 111 | 26 | 132 | 22 | 115 | 39 |
| S474A | 94 | 2 | 74 | 18 | 77 | 5 | 112 | 38 |
| L475A | 143 | 81 | 77 | 20 | 76 | 5 | 103 | 23 |
| S476A | 113 | 15 | 141 | 79 | 78 | 16 | 106 | 41 |

|  |  |  |  |  |  |  |  |  |  |
| --- | --- | --- | --- | --- | --- | --- | --- | --- | --- |
| V477A | 113 | 23 | 74 | 19 |  | 93 | 3 | 116 | 37 |
| T478A | 148 | 14 | 69 | 29 |  | 90 | 3 | 115 | 11 |
| L479A | 155 | 7 | 53 | 6 |  | 120 | 4 | 112 | 19 |
| V480A | 126 | 24 | 82 | 27 |  | 108 | 76 | 92 | 14 |
| L481A | 139 | 38 | 103 | 73 |  | 94 | 15 | 88 | 4 |
| V482A | 134 | 33 | 121 | 85 |  | 88 | 4 | 100 | 2 |
| G483A | 88 | 29 | 99 | 5 |  | 96 | 34 | 96 | 30 |
| I484A | 87 | 66 | 87 | 18 |  | 114 | 46 | 111 | 21 |
| V485A | 134 | 12 | 67 | 33 |  | 95 | 60 | 114 | 28 |
| T486A | 111 | 32 | 98 | 26 |  | 115 | 9 | 104 | 36 |
| L487A | 140 | 26 | 86 | 11 |  | 75 | 3 | 103 | 40 |
| Y488A | 146 | 64 | 80 | 22 |  | 91 | 67 | 109 | 26 |
| L489A | 111 | 30 | 119 | 49 |  | 74 | 23 | 92 | 3 |
| G490A | 125 | 68 | 100 | 26 |  | 70 | 9 | 107 | 22 |
| V491A | 106 | 43 | 62 | 2 |  | 79 | 17 | 112 | 15 |
| M492A | 106 | 55 | 74 | 54 |  | 100 | 38 | 113 | 24 |
| V493A | 130 | 57 | 108 | 91 |  | 92 | 3 | 91 | 9 |
| Q494A | 97 | 54 | 116 | 59 |  | 87 | 15 | 97 | 11 |
| A495S | 93 | 38 | 95 | 56 |  | 145 | 19 | 98 | 22 |

**Table S2.**
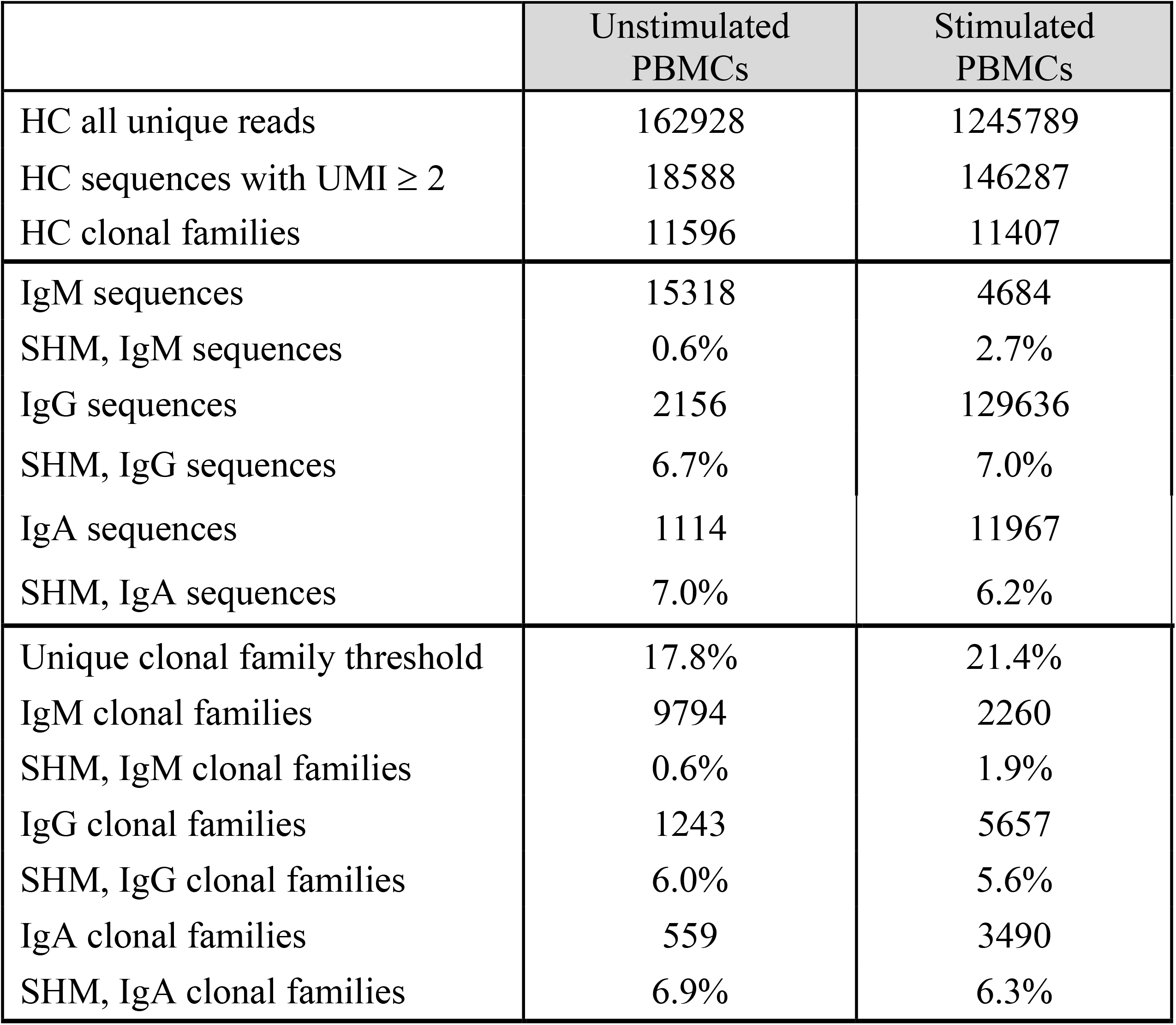
Summary of Ig sequences, clonal families and their corresponding mean SHM in unstimulated PBMCs vs. stimulated PBMCs from DENV-infected patient 013.

**Table S3.** Sequences from unstimulated PBMCs or stimulated PBMCs related to the clonal families of single cell plasmablasts with reactivity to dengue virus isolated from patient 013.

| Clonal family<br>(mAbs) | Germline | Isotype | Reads for all clonotypes |  |
| --- | --- | --- | --- | --- |
|  |  |  | Unstimulated PBMCs | Stimulated PBMCs |
| CF10<br>(C1, A7, I11, L9, P2, G5) | IGHV3-30-3 | total | 143 | 216292 |
|  |  | IgM | 8 | 427 |
|  |  | IgG | 112 | 199151 |
|  |  | IgA | 23 | 16714 |
| CF5<br>(C4, J2, N2) | IGHV1-69 | total | 422 | 141308 |
|  |  | IgM | 9 | 91 |
|  |  | IgG | 413 | 140117 |
|  |  | IgA | 0 | 1100 |
| CF13<br>(N8, F4) | IGHV4-34 | total | 32 | 82814 |
|  |  | IgM | 0 | 195 |
|  |  | IgG | 32 | 76605 |
|  |  | IgA | 0 | 6014 |
| CF7<br>(J8, J9) | IGHV1-69 | total | 579 | 43179 |
|  |  | IgM | 10 | 98 |
|  |  | IgG | 568 | 42315 |
|  |  | IgA | 1 | 764 |
| CF9<br>(K11, L3, M4, M11, O4) | IGHV1-69 | total | 673 | 35793 |
|  |  | IgM | 12 | 103 |
|  |  | IgG | 656 | 34444 |
|  |  | IgA | 5 | 1246 |
| CF8<br>(J3, 403_P4) | IGHV1-69 | total | 10 | 7349 |
|  |  | IgM | 0 | 16 |
|  |  | IgG | 10 | 7001 |
|  |  | IgA | 0 | 332 |
| CF3<br>(405_P4, I13) | IGHV1-18 | total | 10 | 2993 |
|  |  | IgM | 1 | 6 |
|  |  | IgG | 9 | 2899 |
|  |  | IgA | 0 | 88 |
| CF1<br>(B10, M1, D8) | IGHV4-39 | total | 4 | 1087 |
|  |  | IgM | 1 | 11 |
|  |  | IgG | 3 | 1049 |
|  |  | IgA | 0 | 27 |
| CF2<br>(H3, M6) | IGHV1-69 | total | 0 | 1087 |
|  |  | IgM | 0 | 7 |
|  |  | IgG | 0 | 1066 |
|  |  | IgA | 0 | 14 |
| CF11<br>(E9, I8) | IGHV3-30-3 | total | 0 | 19 |
|  |  | IgM | 0 | 0 |
|  |  | IgG | 0 | 19 |
|  |  | IgA | 0 | 0 |

**Figure S1.**
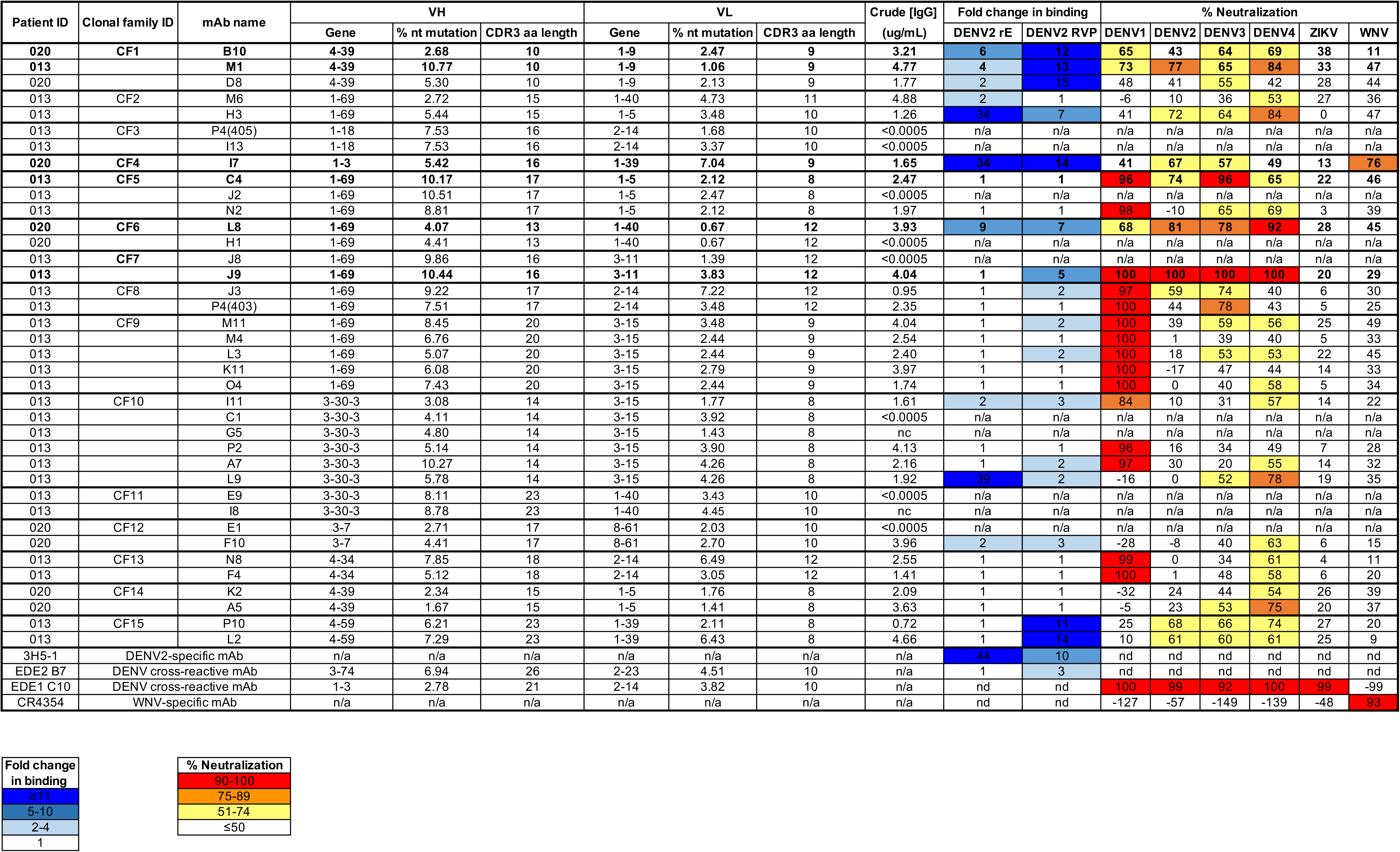
Characteristics of plasmablast-derived mAbs from DENV-infected patients. MAb sequences were identified from single plasmablasts of DENV-infected patient 013 and patient 020, as previously described (Zanini et al., 2018). The patient from which corresponding mAb sequences were identified is listed in the first column, followed by mAb clonal family ID, mAb name, and gene usage, % nucleotide (nt) SHM, and CDR3 amino acid (aa) length for the variable heavy (VH) and light (VL) chain genes. VH and VL sequences were cloned into IgG1 expression vectors and transfected into mammalian cells. Neat crude IgG1-containing culture supernatant was tested for binding to recombinant DENV2 E protein ectodomain and DENV2 RVPs by ELISA, and for neutralizing activity against the indicated RVPs. MAbs 3H5-1 (2 µg/mL), EDE2 B7(2 µg/mL) and EDE1 C10 (10 µg/mL), and CR4354 (2 µg/mL) were used as controls. MAb binding activity is expressed as fold-change in absorbance values over negative control wells containing media only. The heatmap (light to dark blue) indicates strength of binding, as defined in the key below the table. A value of 1 indicates no increase in binding relative to negative control wells. Percent neutralization was calculated using the formula: (% infection in the absence of IgG1 - % infection in the presence of IgG1) / (% infection in the absence of IgG1) x 100. The heatmap (yellow to red) indicates the range of neutralization potencies as indicated in the key below the table. Results are representative of 2 independent experiments. Under the crude IgG column, a value of <0.0005 indicates undetectable levels of IgG1 in crude culture supernatant. Antibodies selected for further characterization are shown in bold. “N/a,” not applicable; “nc,” not successfully cloned; “nd,” not determined.

**Figure S2.**
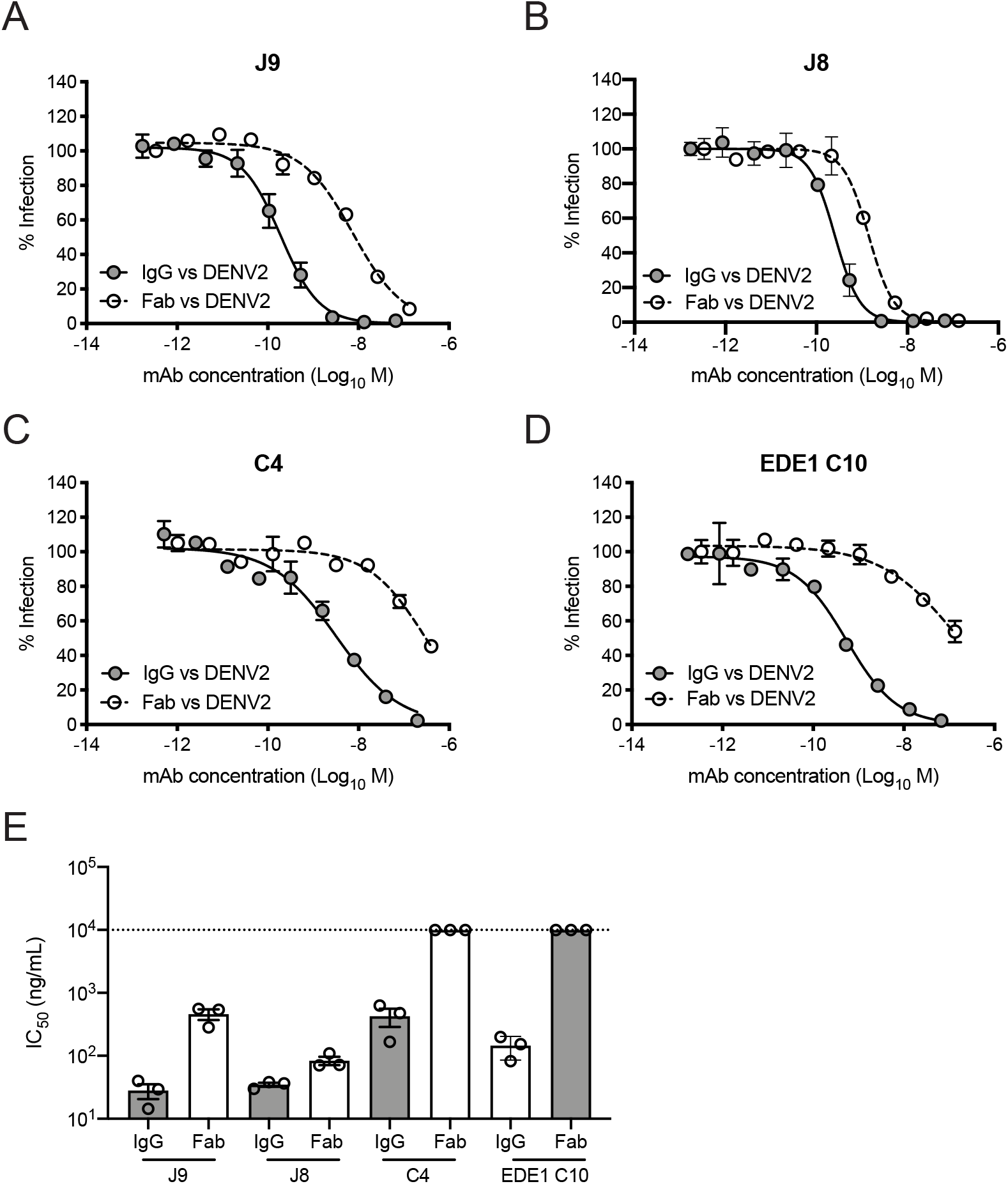
Neutralization potency of IgG and Fab fragments. **(A)** J9, **(B)** J8, **(C)** C4, and **(D)** EDE1 C10 were tested as Fab fragments or full-length IgG for neutralization of DENV2 RVPs. Dose-response neutralization curves represent 3 independent experiments, each performed in duplicate. Data points and error bars indicate the mean and range, respectively. **(E)** Mean IC_50_ values of the indicated IgG or Fab fragment from 3 independent experiments represented by data points. Error bars represent the SD. Values at the dotted horizontal line indicates that 50% neutralization was not achieved at the highest concentration of IgG or Fab tested. Fabs were tested at 2x excess molar concentration relative to IgG.

**Figure S3.**
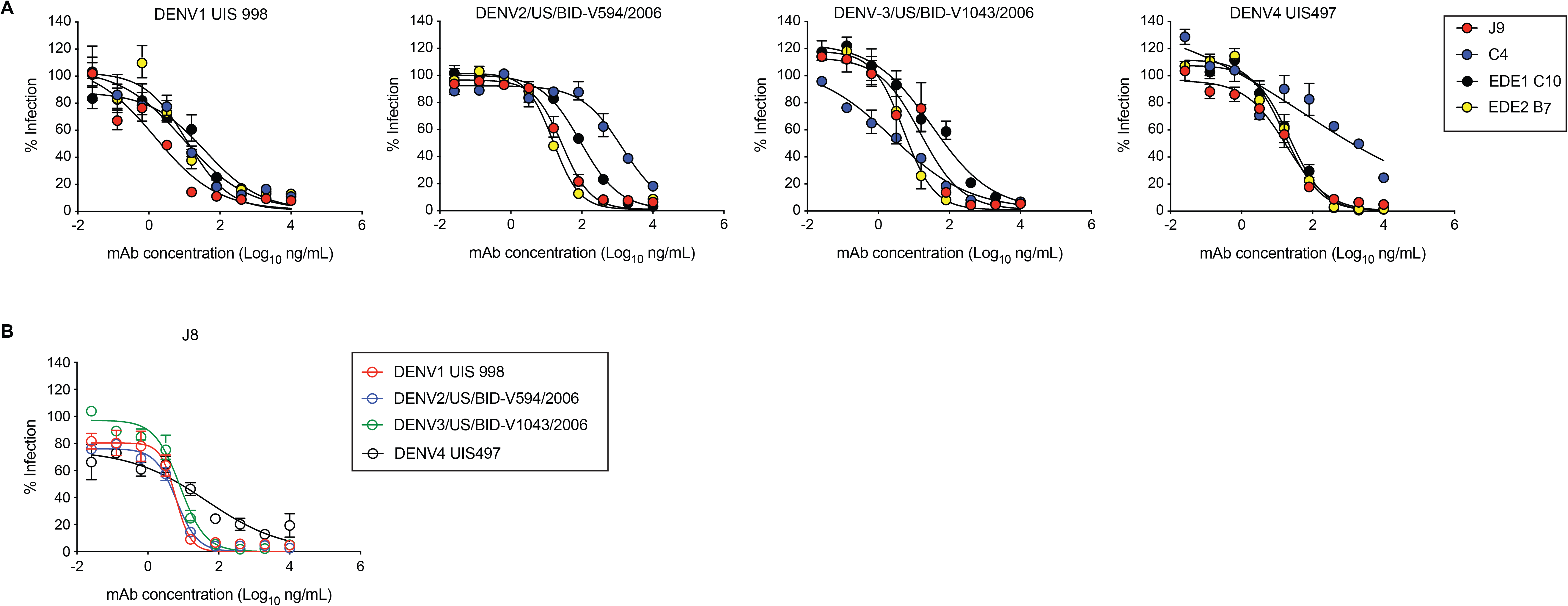
Neutralization potency of mAbs against contemporary DENV1-4 strains. Neutralization of contemporary DENV1-4 strains by **(A)** J9, C4, and **(B)** J8 was determined by intracellular staining with AF488-conjugated 4G2 at 48 h post-infection. Error bars indicate the range of duplicate infections. Infectivity levels were normalized to those in the absence of neutralizing antibody. Dose-response neutralization curves represent one experiment.

**Figure S4.**
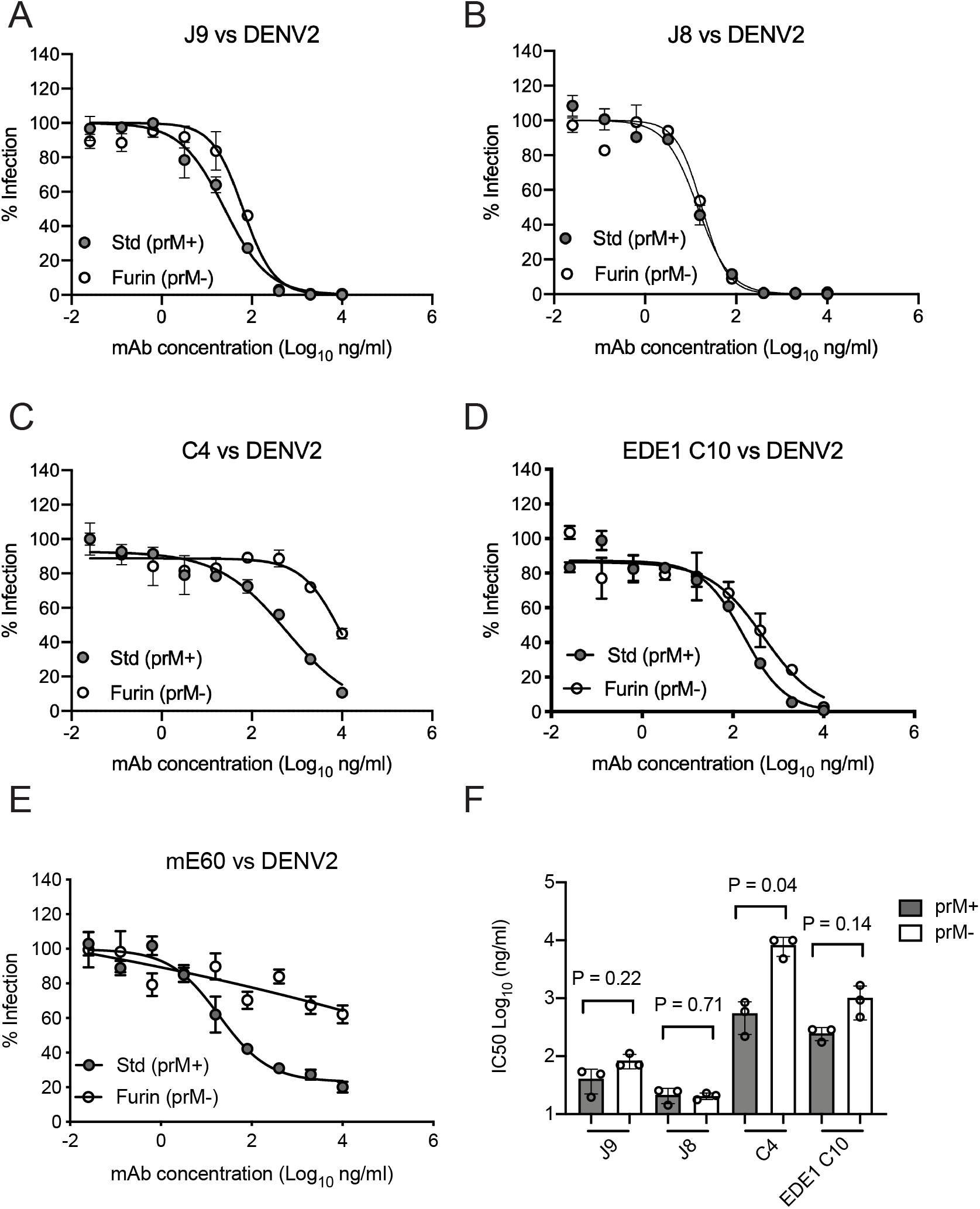
mAb neutralization of standard and mature RVP preparations. **(A)** Dose-response neutralization curves for the indicated mAbs against DENV2 RVPs prepared under standard conditions (Std) or in the presence of overexpressed furin to generate mature RVPs (Furin). Data points and error bars indicate the mean and range of infectivity in duplicate wells, respectively. **(B)** Mean IC_50_ values of mAbs against standard or mature RVPs from 3 independent experiments depicted by data points. Error bars indicate the SD. P-values were obtained from two-tailed paired t-tests.

**Figure S5.**
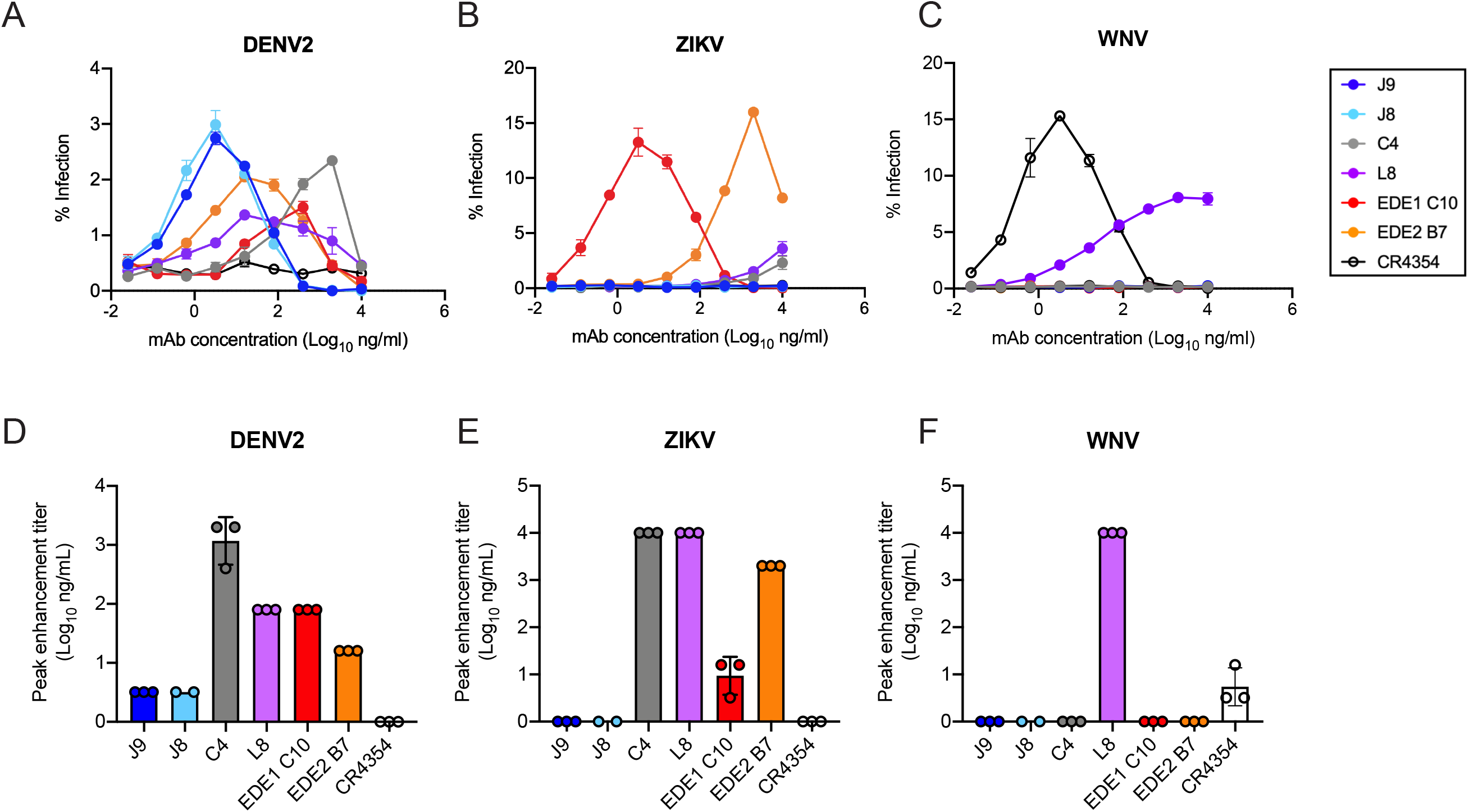
Antibody-dependent enhancement (ADE) of DENV2, ZIKV and WNV infection. Serial dilutions of the indicated mAbs were tested for ADE of **(A)** DENV2, **(B**) ZIKV or **(C)** WNV RVP infection of K562 cells. Error bars indicate the mean of infection in duplicate wells. Bar graphs represent average mAb concentrations at peak enhancement of **(D)** DENV2, **(E)** ZIKV or **(F)** WNV RVP infection obtained from 2-3 independent experiments, each represented by a data point. Where indicated, error bars represent the SD.

**Figure S6.**
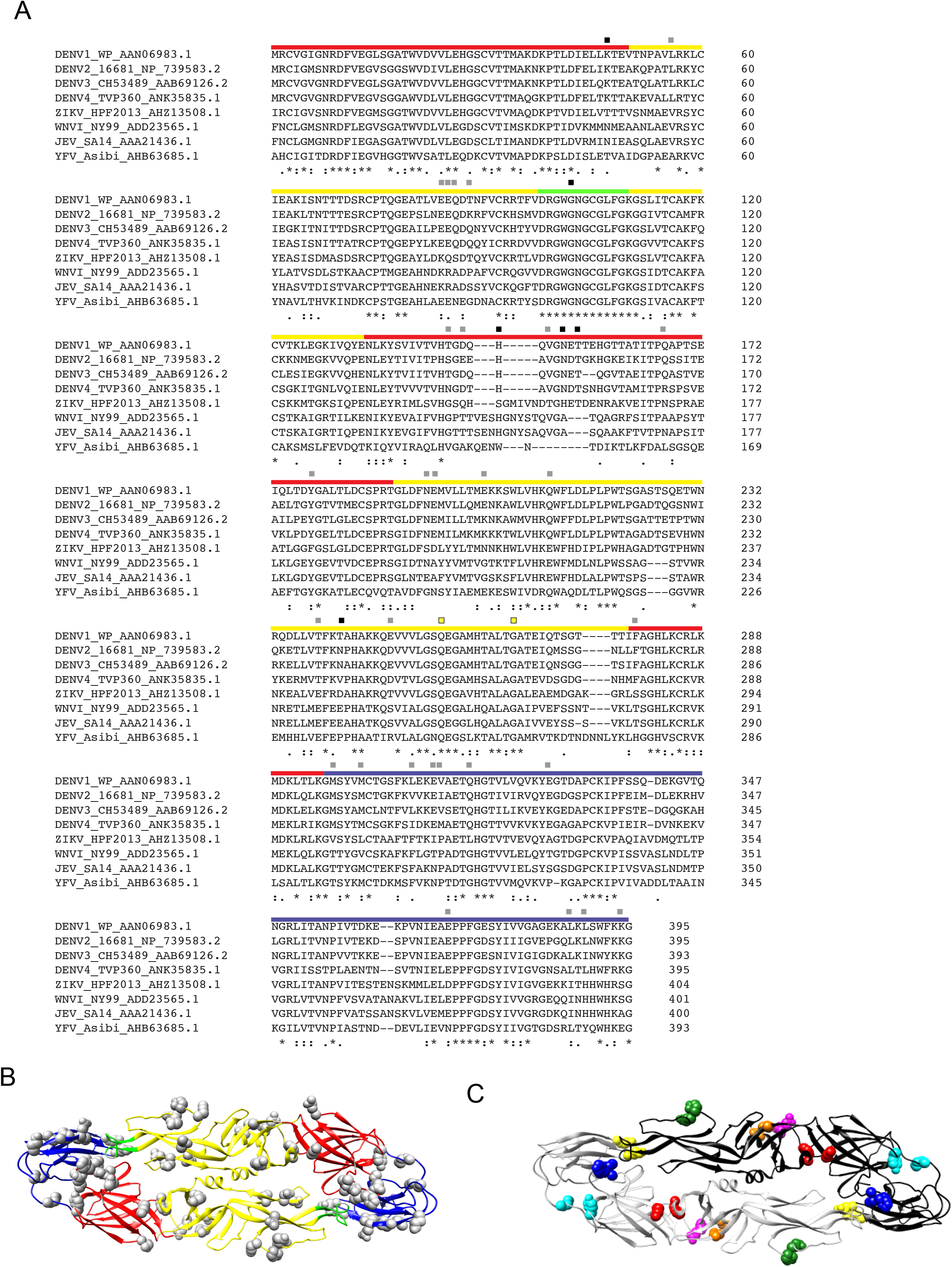
DENV2 E protein mutagenesis. **(A)** Alignment of DENV1-4, ZIKV, and WNV E ectodomain amino acid residues obtained using ClustalW2. Red, yellow, green, and blue bars above the alignment indicate residues within E protein DI, DII, DII-fusion loop, and DIII, respectively. Squares above colored bars indicate residues selected for mutagenesis and generation of RVP variants tested for sensitivity to J9 neutralization: gray squares = no effect on neutralization sensitivity; black squares = reduced sensitivity to J9 neutralization. Yellow squares indicate residues conserved across flaviviruses and important for mAb I7 binding. The sequence used for ZIKV H/PF/2013 differs at two amino acids (residue 246 K>R and 345 M>I from GenBank accession number AHZ13508.1, as previously described (Dowd et al., 2016). **(B)** Ribbon structure of the DENV2 E dimer (PDB 1OAN) with 34 individually mutated residues shown as gray spheres. E protein domains are color-coded as in (A). **(C)** Ribbon structure of the DENV2 E dimer (PDB 1OAN) with one monomer shown in black, and the other in gray. Colored spheres indicate the locations of paired mutations: K47T+V151T (red); L56V+Q211E (orange); E85Q+Q86S (green); H149S+V151T (blue); Y178F+M287V (cyan); N194S+E195D (purple); Q316L+K394S (yellow).

**Figure S7.**
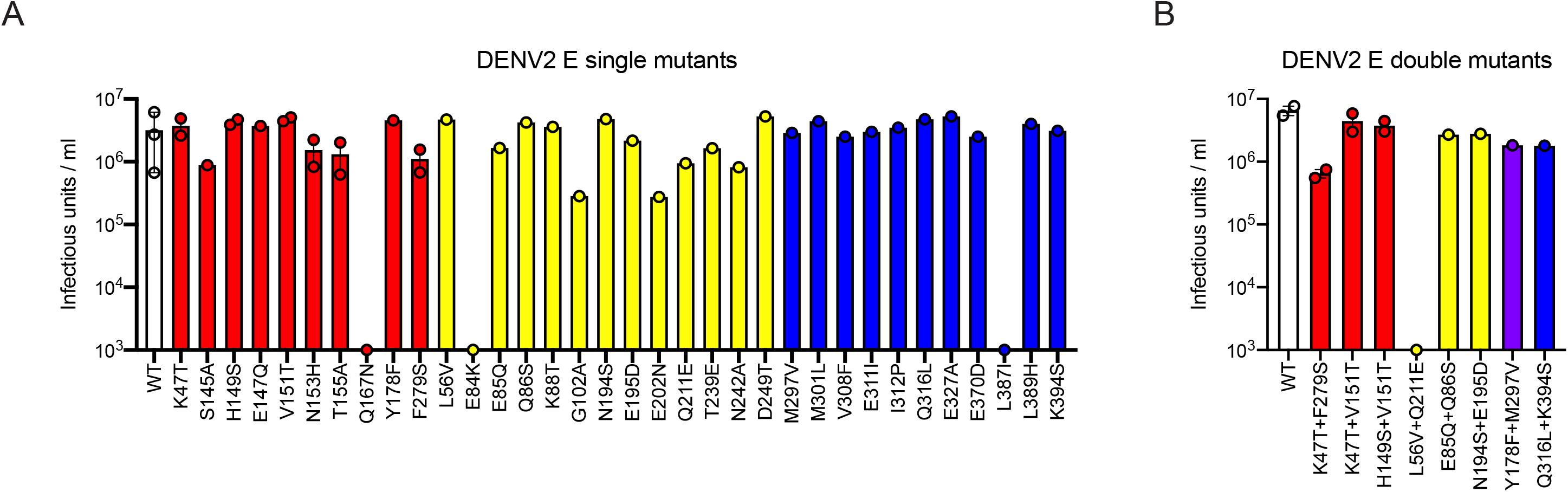
Infectious titer of DENV2 RVPs encoding E protein mutations. Infectious titers of DENV2 RVP encoding **(A)** single or **(B)** double E protein mutations. For each graph, white bars show the infectious titer of WT DENV2 RVP, and red, yellow, and blue bars represent mutations in DI, DII, and DIII, respectively. In (B), the purple bar represents a paired mutation at one residue in DI (Y178F) and another in DIII (M297V). Titers are based on one or two independent RVP preparations, as indicated by data points. Where present, error bars represent the range of infectivity from 2 independent RVP preparations.

**Figure S8.**
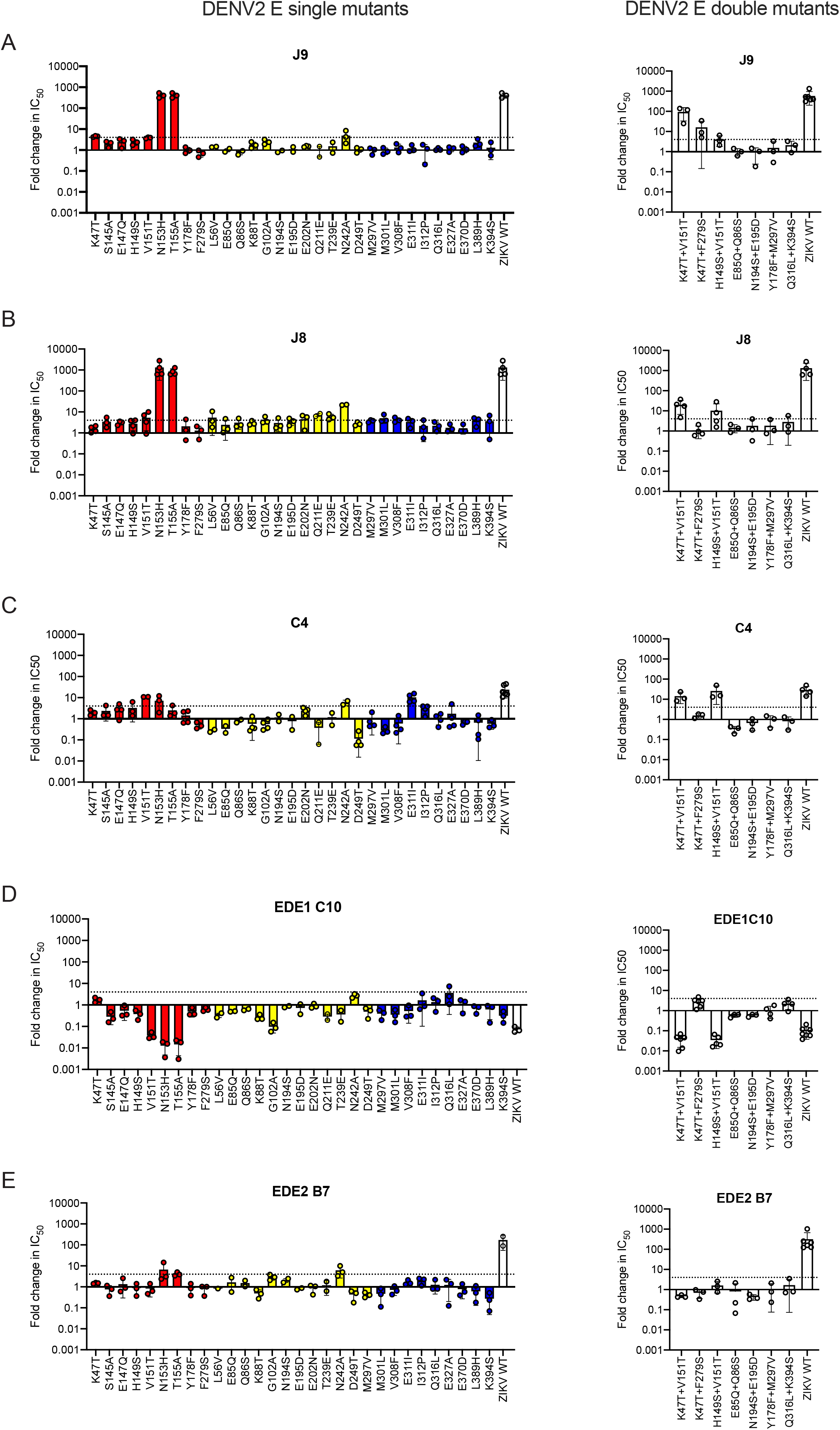
Effect of E protein mutations mAb neutralization potency. We screened a panel of DENV2 RVP variants encoding single (left) or double (right) E protein mutations for sensitivity to neutralization by mAbs **(A)** J9, **(B)** J8, **(C)** C4, **(D)** EDE1 C10, and **(E)** EDE2 B7. Bar graphs represent average fold-change in IC_50_ relative to WT DENV2 RVP obtained from at least 2 independent experiments, as indicated by data points. Error bars indicate the range (n = 2) or SD (n > 2). The dotted line represents a 4-fold increase in IC_50_ relative to DENV2 WT. On the left panel, red, yellow, and blue bars indicate mutations are residues in DI, DII, and DIII, respectively. For each mAb, neutralization of WT ZIKV RVPs is included as a control.

**Figure S9.**
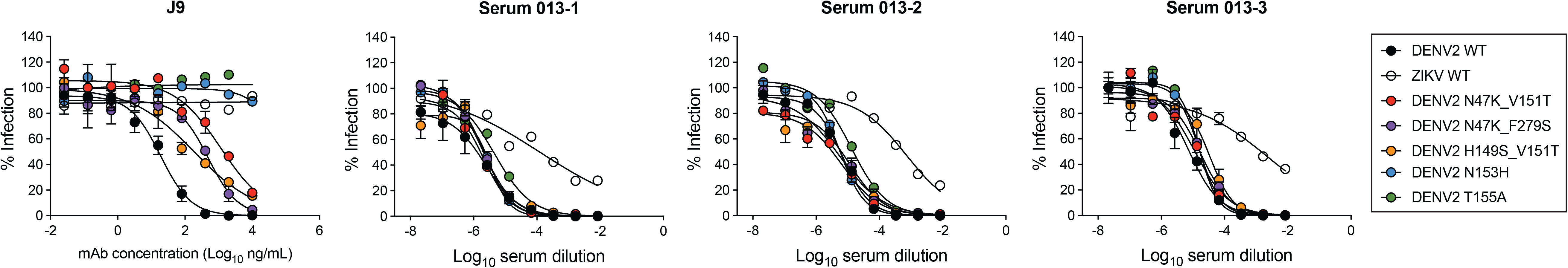
Effect of E protein mutations on patient 013 serum neutralization potency. Neutralizing activity of mAb J9 and longitudinal serum samples from patient 013 were tested against DENV2 RVP variants encoding E protein mutations that reduced J9 neutralization potency. ZIKV WT RVP was included as a control. Serum samples 013-1, 013-2 and 013-3 were collected 4, 8 and 22 days after onset of fever, respectively. Dose-response neutralization curves are representative of 3 independent experiments, each performed in duplicate. Error bars indicate the range of infectivity normalized to infection levels in the absence of antibody.

**Figure S10.**
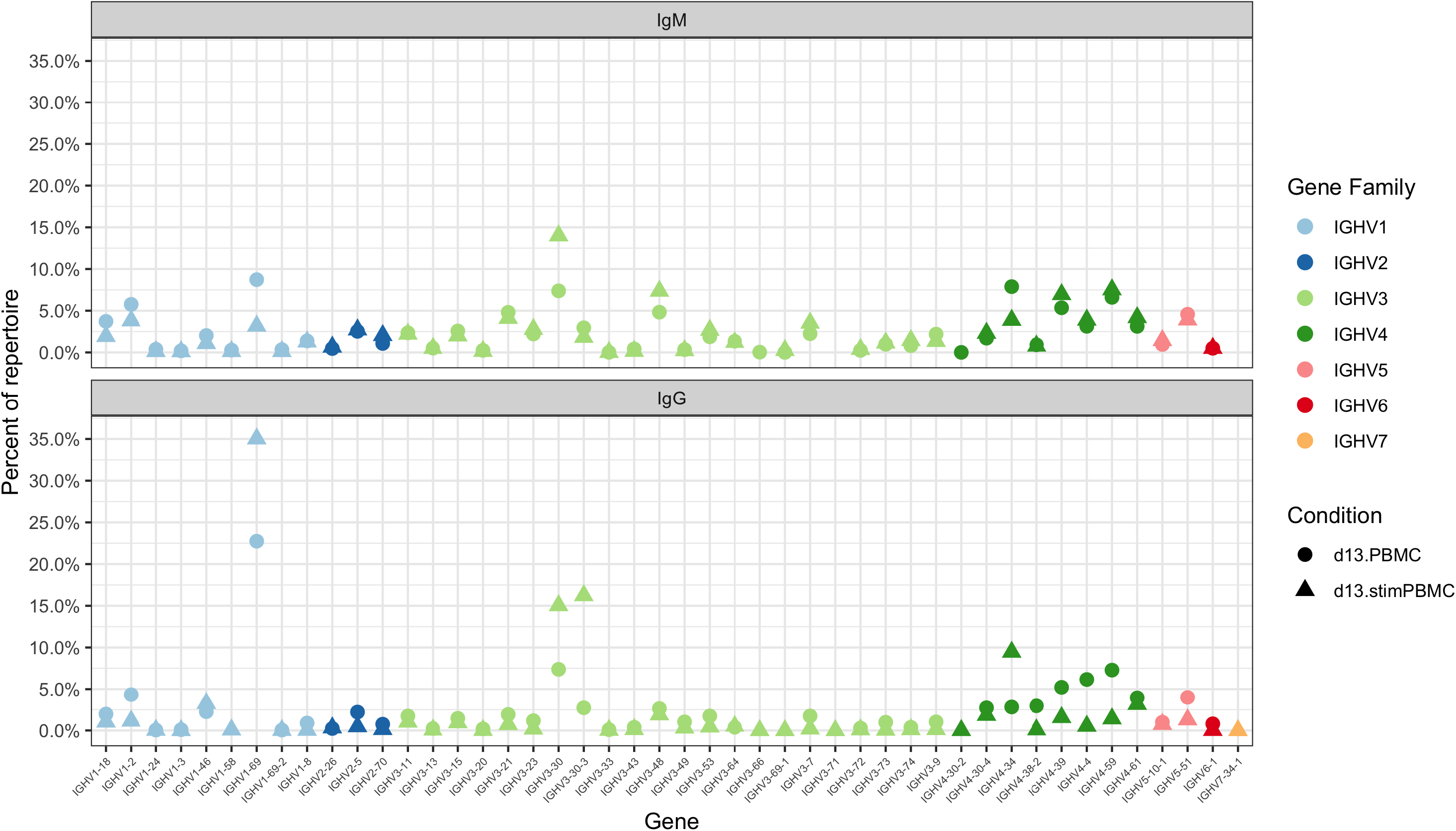
Patient 013 VH gene usage. Germline VH gene usage among the unstimulated (circles) and stimulated (triangles) PBMC repertoires of patient 013, with only IgM (top) and IgG (bottom) sequences shown. Genes are grouped by color into gene families, as indicated in the key.

**Figure S11.**
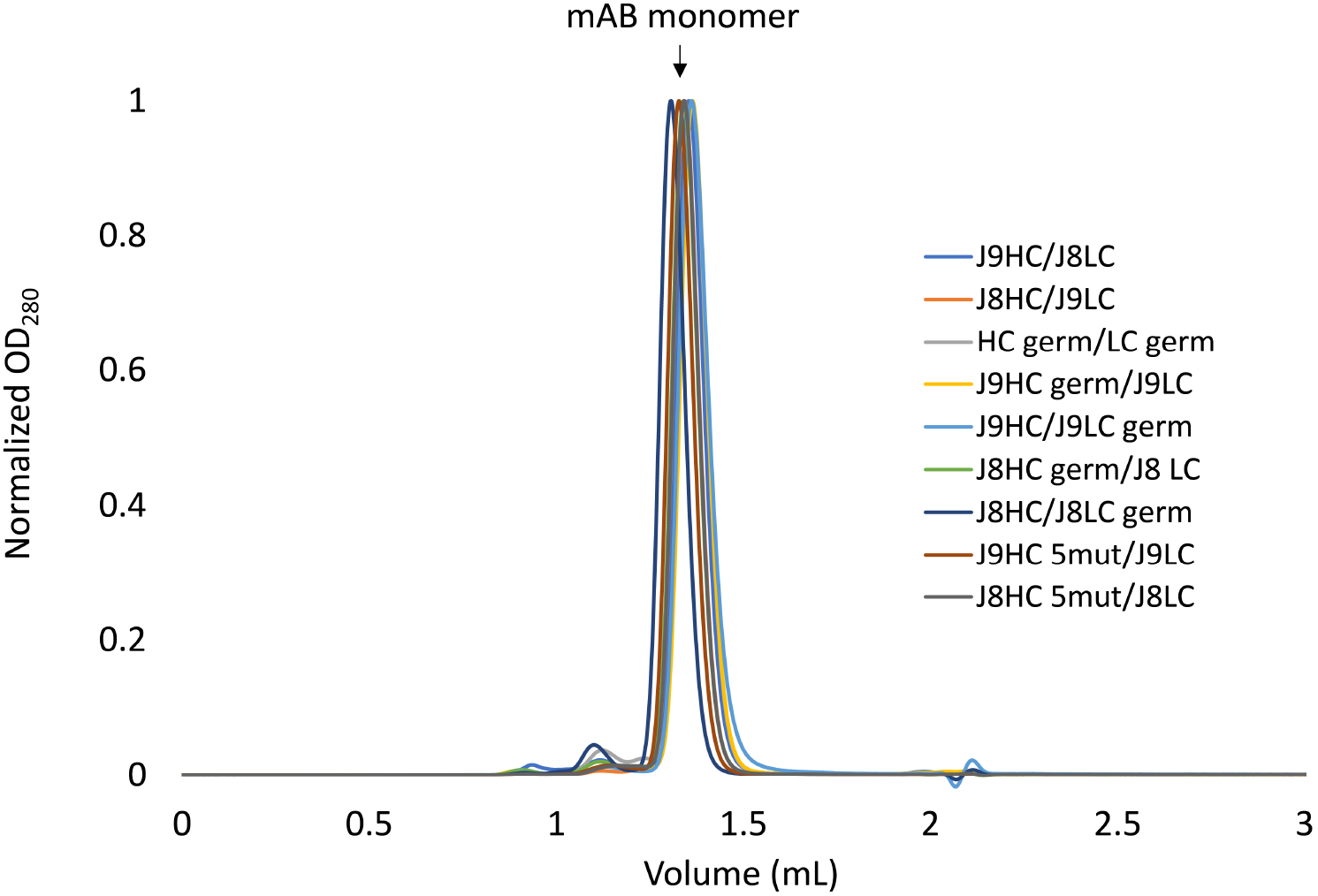
Analytical size exclusion chromatography of recombinant IgGs. Biophysical characterization of J9 and J8 IgG variants by analytical size exclusion chromatography (Superdex S200 Increase 3.2/300). A single major peak at 280 nm corresponding to monomeric IgG is observed.

